# Human ATG3 contains a non-canonical LIR motif crucial for its enzymatic activity in autophagy

**DOI:** 10.1101/2022.08.02.502437

**Authors:** Jakob Farnung, Matthias Muhar, Jin Rui Liang, Kateryna A. Tolmachova, Roger M. Benoit, Jacob E. Corn, Jeffrey W. Bode

## Abstract

Macroautophagy is one of two major degradation systems in eukaryotic cells. Regulation and control of autophagy is often achieved through the presence of short peptide sequences called LC3 interacting regions (LIR) in autophagy-involved proteins. Using a combination of new protein-derived activity-based probes, protein modelling and X-ray crystallography, we identified a non-canonical LIR motif in the human E2 enzyme responsible for LC3 lipidation, ATG3. The LIR motif is present in the flexible region of ATG3 and adopts an uncommon β-sheet structure binding to the backside of LC3. We show that the β-sheet conformation is crucial for its interaction with LC3. *In cellulo* studies provide evidence that LIR^ATG3^ is required for LC3 lipidation and ATG3∼LC3 thioester formation. Removal of LIR^ATG3^ negatively impacts the rate of thioester transfer from ATG7 to ATG3.

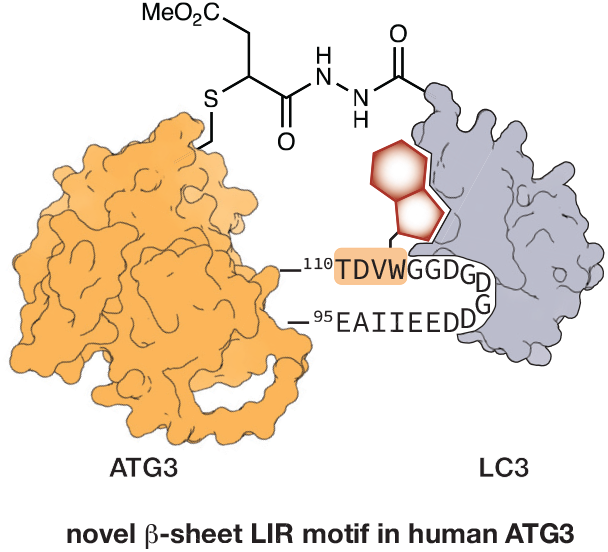

---

Macroautophagy (herein referred to as autophagy) is a major catabolic process in eukaryotic cells,^1^ responsible for the bulk degradation of various cellular components such as proteins^2^, cellular compartments^3^ and pathogens^4^. Akin to the ubiquitin proteasome system, two families of small protein modifiers are crucial regulatory elements of autophagy. Proteins of the LC3 or GABARAP families are conjugated via their C-terminal glycine residue to phosphatidylethanolamine-containing lipids.^5^ This process is catalyzed by an intricate enzymatic cascade in which ATG7 functions as an activating E1 enzyme using ATP to form a thioester with LC3/GABARAP and transferring them to an E2 enzyme, ATG3, via a trans-thioesterification reaction. ATG3 performs the lipidation of LC3/GABARAP in conjunction with an E3-like enzyme complex of isopeptide-linked ATG5-ATG12. Membrane tethering of LC3/GABARAP is crucial for membrane expansion of autophagosomes to engulf the autophagic cargo and for eventual lysosome fusion^6-9^. But the mechanistic details of LC3/GABARAP-lipidation by ATG3 remain enigmatic.

A conserved feature of the autophagy pathway is the recurring presence of short peptide motifs called LC3 interacting regions (LIR) in proteins associated with autophagy^10^. Proteins containing LIR motifs are recruited to LC3/GABARAP through hydrophobic interactions. The core sequence of LIR motifs, ΦxxΨ, is characterized by the presence of an aromatic residue (Φ) – Trp Phe and Tyr – followed by two variable positions and an aliphatic, hydrophobic amino acid (Ψ), generally Ile, Leu, or Val^11^. The LIR motif adopts an extended conformation forming an intermolecular β-sheet with β2 of LC3/GABARAP. A variety of LIR motifs have been identified in selective autophagy receptors, which employ them to recruit cargos to expanding autophagosomes^2^. In addition, these motifs can also be found in proteins involved in the attachment of LC3/GABARAP to membranes such as ATG4^12^, a protease required for processing of proLC3/proGABARAP and delipidation of LC3/GABARAP.

Few chemical probes for autophagy have been developed^13-15^. The majority are inhibitors of autophagy proteins functioning upstream of the lipidation cascade, such as wortmannin, a PI3K inhibitor that abrogates localization of lipidation enzymes to expanding autophagosomes^16,17^. Despite the similarity to the ubiquitin proteasome pathway, only few chemical probes exist for LC3/GABARAP lipidation^18,19^. Hemelaar *et al*. reported the synthesis of LC3/GABARAP activity-based probes by direct aminolysis and employed these probes to identify ATG4 as the processing protease of proLC3/proGABARAP^20^. However, access to these probes by direct aminolysis is generally hampered by harsh reaction conditions and inefficient conversion.

Herein, we report the facile preparation of GABARAP and LC3A activity-based probes (ABPs) using a recently established hydrazide acylation protocol^21^. Access to these ABPs led us to identify an unknown non-canonical LIR motif embedded in an unusual β-sheet conformation in human ATG3.

Further investigation of this motif with peptide binders, x-ray crystallography, and CRISPR-enabled *in cellulo* studies revealed that this LIR motif is crucial for the enzymatic function of ATG3.

## Results

### Preparation of GABARAP and LC3A activity-based probes

We recently reported access to Ubl proteins bearing thiol-reactive electrophiles at their C-terminus by chemoselective acylation of recombinant protein-hydrazides with carboxylic acid anhydrides. The lack of ABPs for autophagy inspired us to prepare GABARAP ABPs using this protocol. GABARAP was expressed as the C-terminal glycine deletion (ΔG116) to conserve the atomic register at the C-terminus after the introduction of electrophilic groups; the resulting acyl hydrazides mimic the native glycine and place the electrophile close to the reactive site of the native C-terminus (Extended Data Fig. 1). GABARAP(ΔG116)-NHNH_2_ was successfully converted to α-chlorocetyl probe **1** and methyl fumarate probe **2**, in analogy to probes that showed excellent activity in the UFM1-pathway (Fig. 1a)^22^.

**Fig. 1.**
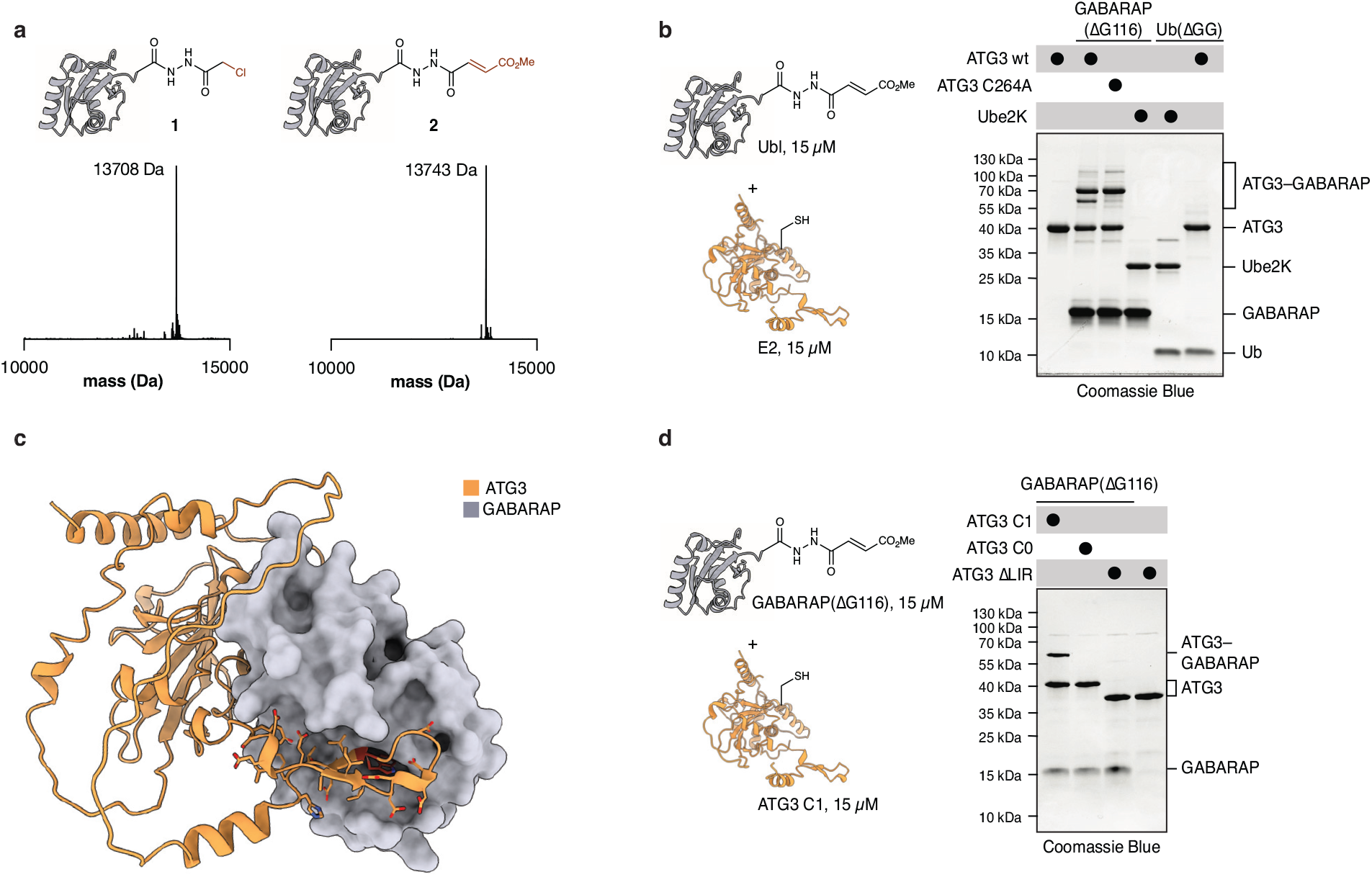
Modification of ATG3 with GABARAP ABPs depends on LIR^ATG3^. **a**, Characterization of GABARAP activity-based probes, **1** and **2**, obtained by hydrazide-acylation. Deconvoluted mass-spectrum (ESI) of GABARAP(ΔG116)–NHNH α-chloroacetyl **1**. Expected mass 13707 Da. Deconvoluted mass-spectrum (ESI) of GABARAP(ΔG116)–NHNH methyl-fumarate **2**. Expected mass 13743 Da. **b**, Reaction of **2** or Ub(ΔGG)–NHNH methyl fumarate probe (15 μM) with recombinant E2s ATG3 and Ube2K (15 μM). The reaction was allowed to proceed for 1 h and was analyzed by SDS-PAGE and Coomassie Blue staining. **c**, ColabFold-predicted structure of the protein-complex of ATG3 and GABARAP. Side-chain atoms are shown for the predicted LIR motif in ATG3 (90-112). **d**, Reaction of **2** (15 μM) with ATG3 variants C1, C0 and ΔLIR (15 μM). ATG3 C1 contains only active-site cysteine C264. ATG3 C0 contains no cysteines. ATG3 ΔLIR lacks amino acids 95-111. Reaction was allowed to proceed for 1 h and was analyzed by SDS-PAGE and Coomassie Blue staining. Full-gel images for **b** and **d** are available in the Source Data File.

To assess the reactivity of the probes, we incubated **1** and **2** (15 μM) with recombinantly expressed ATG3 (15 μM) and its catalytically inactive variant C264A (Fig. 1b, Supplementary Fig. 1). Probe **2** reacted with ATG3 efficiently leading to the formation of multiple ATG3–GABARAP bands. Mutation of active-site cysteine 264 to alanine caused loss of one ATG3–GABARAP band but did not abrogate additional bands for the complex, indicating unspecific modification of ATG3, likely due to the presence of multiple cysteine residues in flexible regions of ATG3. We, therefore, demonstrated that C264 is one of the *bona fide* target cysteines of probe **2** despite its cross-reactivity with other nearby cysteines. Probe **2** nonetheless reacted specifically with its cognate E2 as ubiquitin E2 Ube2K did not react with **2** and ATG3 wt did not react with Ub(ΔGG) methyl fumarate probe. We were intrigued by the cross-linking efficiency of **2** with ATG3 and to investigate the origin of this efficiency, we generated a C1 variant of ATG3 that only contains the catalytic cysteine 264 and a corresponding C0 variant containing no cysteines. Probe **2** showed excellent reactivity with ATG3 C1 and, as expected, no reaction was observed with ATG3 C0 (Fig. 1d, lanes 1-2). ATG3 C1 maintains its cross-linking efficiency compared to ATG3 wt indicating that cross-linking is not due to the presence of numerous cysteine residues in ATG3 but likely due to higher affinity between ATG3 and GABARAP. In general, the affinity of E2s for their cognate Ubl is quite low as the thioester transfer from the E1 to the E2 enzyme is facilitated by additional interactions from the UFD domain of the E1 enzyme^23^. The specificity and high reactivity of our probes with human ATG3 indicate that ATG3 may contain additional binding elements, resulting in a tighter interaction than generally observed for E2s and their Ubl cognates^24^.

### Human ATG3 contains a non-canonical LIR motif

Previous reports established that yeast ATG3 contains an ATG8 interaction motif required for interaction with ATG8, the yeast analogue of LC3 and GABARAP^25^. However, this motif is not conserved in human ATG3 and therefore did not explain the enhanced interaction of human ATG3 with GABARAP (Supplementary Fig. 2). As there was no structural information available on human ATG3 that could explain the higher affinity of ATG3 for GABARAP, we turned to computational methods. Artificial intelligence driven modelling such as AlphaFold has shown great promise in accessing structural data on proteins for which no structural data is available^26^. Recent improvements have even enabled researchers to model protein-protein interactions^27^. We employed an open-source modelling tool, ColabFold, to model the protein-protein interaction between ATG3 and GABARAP (Fig 1c, Supplementary Fig. 3-4)^28^. Intriguingly, ColabFold modelled a complex of ATG3 and GABARAP that resembled the canonical closed-conformation observed in E2-Ub thioester complexes^29^. Additionally, a section of the flexible region of ATG3, L94-Y111, folded into a short β-sheet that was bound to a groove formed by helices α2 and α3. The interaction site on GABARAP was equivalent with the canonical binding region of LIR motifs. Inspection of the β-sheet sequence shows a sequence motif, W^107^VDT^110^, reminiscent of core LIR motifs but containing threonine instead of the canonical Ile, Leu or Val residues. The binding mode of the WDVT motif was similar to binding modes observed for previously investigated LIR motifs; W107 binds to hydrophobic pocket (HP) 1 and T110 to HP2. Therefore, the WVDT motif likely represents a non-canonical LIR motif previously undiscovered in human ATG3, in which T110 binds to GABARAP instead of the canonical aliphatic amino acids. The β-sheet forms further interactions with GABARAP outside of the core LIR motif. Several ionic interactions between Asp and Glu residues in the peptide and Lys / Arg residues surrounding the binding site seem to further stabilize the interaction.

LIR^ATG3^ was predicted to be embedded in a β-sheet, a conformation scarcely found in reported LIR motifs. Most LIR motifs are embedded in an extended conformation that forms an intermolecular β-sheet with β2 of LC3/GABARAP. The presence of LIR-motifs within β-sheets has only recently been reported for the pathogenic protein RavZ,^30^ and FNIP^31^ (vide infra). However, several non-canonical LIR motifs have been reported. For example, UBA5 binds GABARAP using two aliphatic residues and an additional aromatic amino acid outside of the core LIR sequence^32^ and NDP52 exclusively binds to LC3C via its cLIR motif consisting of only aliphatic residues^33^. Nevertheless, even these non-canonical motifs bind in an extended linear fashion to LC3/GABARAP.

We hypothesized that the putative non-canonical LIR motif in ATG3 was responsible for its high affinity interaction with GABARAP, resulting in the efficient reaction of our probes with ATG3. We sought to test this with a cross-linking assay and chose ATG3 variants C1 and C0 (vide supra) to facilitate analysis. We expressed a variant of ATG3 C1 lacking the LIR motif (Δ95-111), ΔLIR. Incubation of **2** with ATG3 ΔLIR showed no complex formation by SDS-PAGE analysis. (Fig 1d). Removal of the LIR motif by deletion of residues 95-111 had the same effect as removal of the catalytic cysteine C264. The same LIR-dependence was observed for cross-linking with wild-type ATG3, which retained all of its cysteine residues (Extended Data Fig. 2). This observation was independent of the probe used; reaction of probe **1** and cross-linking with ATG3 were also strictly dependent on LIR^ATG3^. These findings support the ColabFold model that ATG3 contains an additional binding element for GABARAP through its LIR motif.

GABARAP is representative of one of two protein families conjugated by ATG3. The LC3-family is also tethered to membranes by ATG3. To test the general involvement of LIR^ATG3^ in ATG3 activity beyond GABARAP we also prepared LC3A probes **3** and **4**. We reacted these probes with ATG3 C1, C0 and ΔLIR. As observed for GABARAP, LC3A shows efficient reaction with ATG3 C1 but no reaction with either C0 or ΔLIR variants (Extended Data Fig. 3). These results indicate that LIR^ATG3^ has a general role in ATG3 binding to LC3/GABARAP.

### LIR^ATG3^ motif forms a β-sheet

To exclude an effect of LIR deletion on protein activity we performed a competition experiment with chemically prepared LIR (L94-H112) peptide **5**. Probe **2** (15 μM) was incubated with ATG3 C1 in the presence of increasing amounts of **5**. The peptide blocked ATG3 modification in a concentration dependent manner with an IC_50_ of 104 μM, indicating that the LIR motif is involved in binding GABARAP (Fig. 2a,b,d). We also envisioned that cyclization of the LIR peptide would recapitulate its β-sheet conformation and lead to tighter binding than the linear peptide due to preorganization^34^. Cyclic peptide **6** was prepared by solid-phase peptide synthesis followed by cyclization using selective cysteine alkylation (Extended Data Fig. 3)^35,36^. peptide **6** inhibited the reaction of **2** with ATG3 C1 with an IC_50_ of 24 μM (Fig. 2c,d). This IC_50_ is 5-fold lower than the IC_50_ observed for the linear peptide **5**, suggesting that the β-sheet conformation is indeed crucial for the interaction of the ATG3 LIR motif with GABARAP. CD spectroscopic analysis of the peptides showed that linear peptide **5** is not structured in solution. Upon cyclization **6** remains mostly unstructured with some transition to an organized structure (Supplementary Fig. 5). Additionally, we confirmed the interaction of LIR^ATG3^ peptides with GABARAP and LC3A by fluorescence polarization (Fig. 2e-g). Linear and cyclic peptides bind to both LC3A and GABARAP. As indicated by the competition experiments, cyclic peptides bound markedly tighter to GABARAP and LC3A than the linear peptides. The stronger affinity of the cyclic peptides supports the importance of the β-sheet conformation in binding to LC3A and GABARAP.

**Fig. 2.**
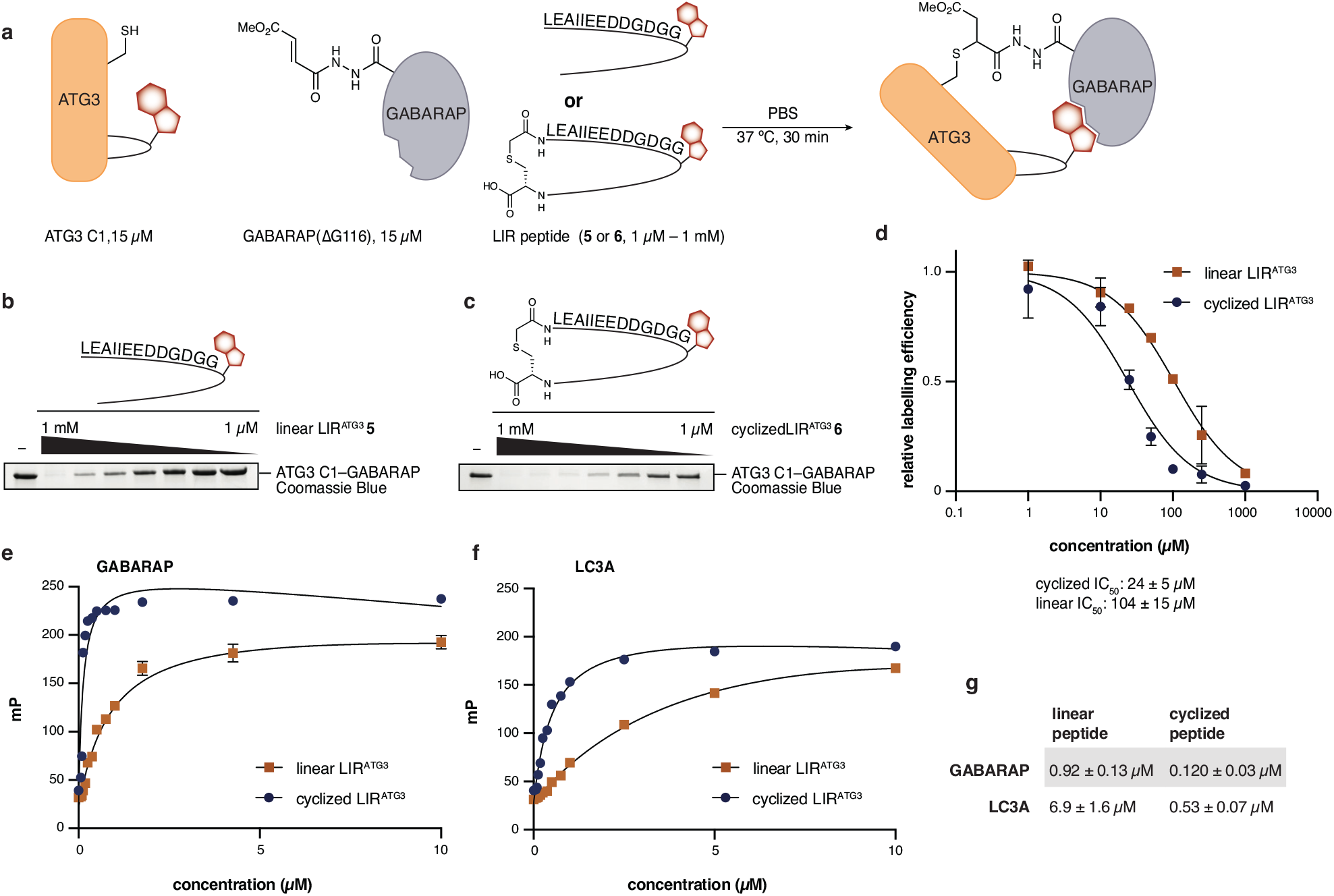
Characterization of LIR^ATG3^. **a**, Reaction scheme outlining competition assay used for **b, c** and **d. b**, Coomassie Blue-stained SDS-PAGE analysis for reaction of **2** with ATG3 C1 in the presence of varying concentrations of linear LIR^ATG3^ peptide **5** as outlined in **a. c**, Coomassie Blue-stained SDS-PAGE analysis for reaction of **2** with ATG3 C1 in the presence of varying concentrations of cyclized LIR^ATG3^ peptide **6** as outlined in **a. d**, Quantification of competition assays shown in **b** and **c**. Amount of GABARAP–ATG3 C1 complex was quantified by gel-densitometry and normalized to the reaction without peptide for each gel. IC_50_ was estimated by non-linear regression. *n* = 2,3 independent experiments with similar results. Data are presented as average values ± s.d. **e**, Fluorescence polarization binding-data of fluorescein-modified LIR^ATG3^ peptide binding to GABARAP. Either linear or cyclized LIR^ATG3^ peptide was used. The measurement was performed in triplicates and data shown as average values ± s.d. K_D_ was estimated using non-linear regression. **f**, Fluorescence polarization data of fluorescein-modified LIR^ATG3^ peptide binding to LC3A. Either linear or cyclized LIR^ATG3^ peptide was used. K_D_ was estimated using non-linear regression. The measurement was performed in triplicates and data shown as average values ± s.d. **g**, Table summarizing K_D_ values obtained in **e, f**. Source data and full-gel images for **b** and **c** are available in the Source Data File.

### Co-crystal structure of GABARAP and LIR^ATG3^

To further corroborate our findings, we co-crystallized GABARAP with an LIR^ATG3^ peptide (Y90-H112). We solved the co-crystal structure at a resolution of 2.6 Å and could resolve amino acids E95-T110 of the LIR^ATG3^ peptide (Extended Data Table 1, Fig. 3a). GABARAP adopted its previously described closed conformation and showed no structural rearrangements^37^. The structure of the LIR peptide and also its interactions with GABARAP were consistent with respect to the ColabFold prediction with a backbone-atom root-mean-square deviation of 0.61 Å (Fig. 3b). ATG3 W107 binds to HP1 *via* hydrophobic interactions to a deep pocket formed by GABARAP P30, K48, F104. The binding mode is identical to the binding mode observed for other LIR motifs (Fig. 3c). ATG3 V108 is buried by hydrophobic interactions with GABARAP K46 and Y49 and the upper strand of the LIR β-sheet. ATG3 D109 forms a surface exposed salt bridge with R28. ATG3 T110 binds to HP2 akin to the more conserved Ile, Leu, Val residues commonly observed in LIR motifs. The shallow pocket is formed by GABARAP Y49, V51, P52, L55, L63. To form the observed β-sheet conformation, residues N-terminal of W107 form a hairpin turn consisting of glycine and aspartic acid residues mainly binding via H-bonding and electrostatic interactions to GABARAP. ATG3 D104 binds to H9 via a hydrogen bond. Aspartic and glutamic acid residues E99, E100, D101 and D102 form salt bridges with various basic residues, K20, K24 and K48 (Fig. 3d). Following residue E99 a short β-sheet is formed for which backbone atoms could be resolved until E95. Overall, the interaction between the ATG3 LIR motif and GABARAP buries 660 Å^2^ [^38^].

**Fig. 3.**
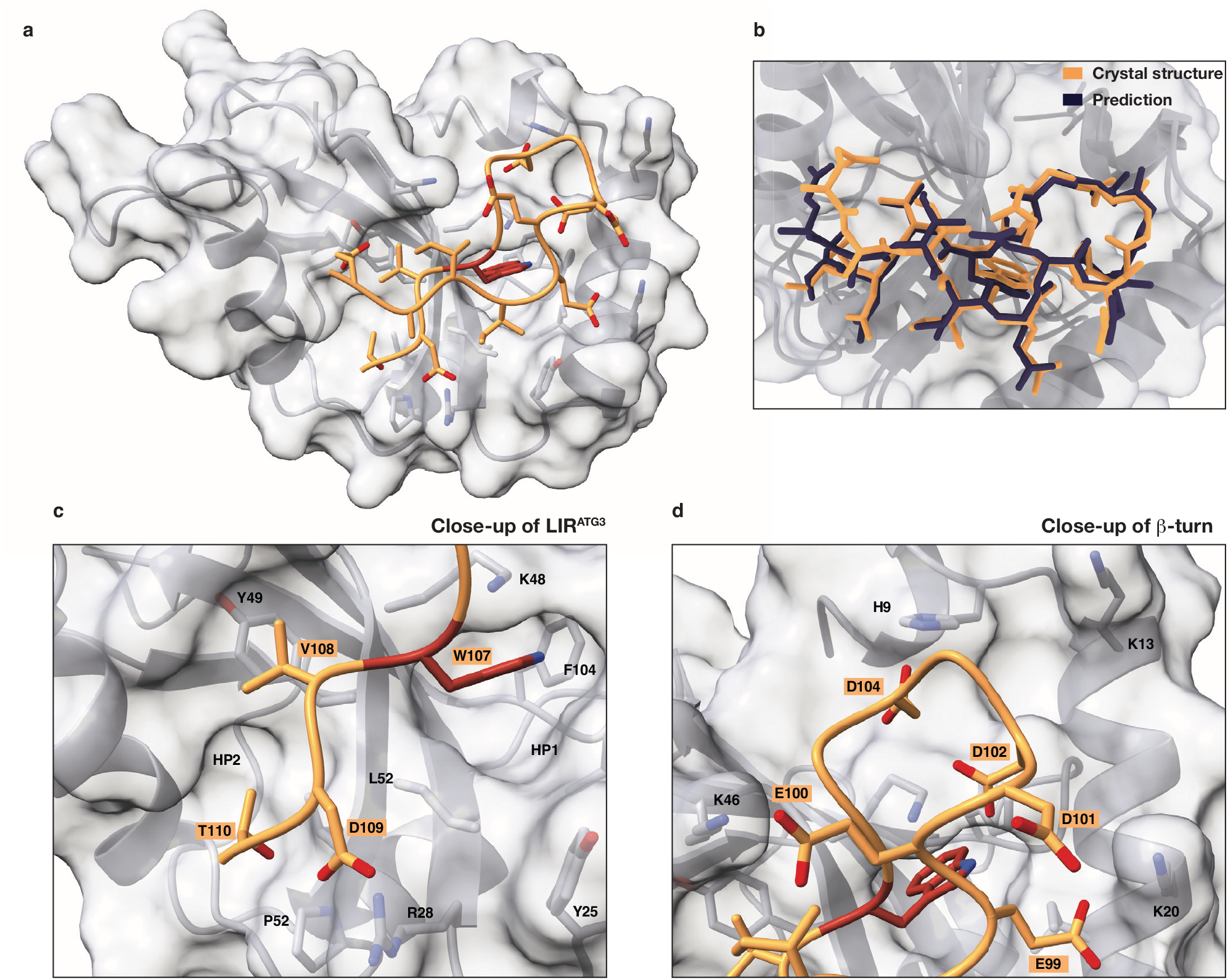
Co-crystal structure of GABARAP and LIR^ATG3^. **a**, Co-crystal structure of GABARAP (grey) and LIR^ATG3^ (orange). Side-chains in contact with LIR^ATG3^ are shown. For clarity W107 is colored in red. One copy of the GABARAP-LIR^ATG3^ complex was chosen from the asymmetric unit. Other copies in the asymmetric unit show similar structures of the complex. **b**, Zoom-in and overlap of LIR^ATG3^ (orange) with structure of LIR^ATG3^ predicted by ColabFold (dark blue); Backbone-atome RMSD is 0.61 Å. **c**, Zoom-in of core LIR^ATG3^ motif binding to hydrophobic pockets HP1 and HP2. **d**, Zoom-in of hairpin turn and the interaction of LIR^ATG3^ with basic residues in GABARAP. Residues E95-D104 have been omitted for clarity.

Based on the predicted ATG3-GABARAP model and our co-crystal structure we identified residues in the LIR motif involved in the ATG3-GABARAP interaction. To probe the contribution of LIR^ATG3^ residues to GABARAP binding we performed an Ala-screen using our previously established cross-linking assay as readout (Fig. 4). Cross-linking efficiency of probe **2** with ATG3 or its mutants is dependent on the affinity of the LIR motif. We expressed ATG3 alanine mutants of the identified residues and reacted them with probe **2**. As expected for LIR motifs, mutation of W107 almost completely abrogated reaction of **2** with ATG3. Similarly, V108A significantly reduced the cross-linking efficiency. This is in agreement with observations for other LIR motifs showing that the first variable position contributes significantly to the binding of LIR motifs^39^. Intriguingly, mutation of D109 or T110 have almost no effect on reaction efficiency. Additionally, the majority of the other screened residues have at least some impact on the conjugation with GABARAP. The hairpin turn binds mainly via electrostatic interactions to GABARAP. Increasing the salt concentration disrupts this interaction as shown by impeded cross-linking with ATG3, highlighting the importance of this part of LIR^ATG3^ for its interaction with GABARAP (Extended Data Fig. 4).

**Fig. 4.**
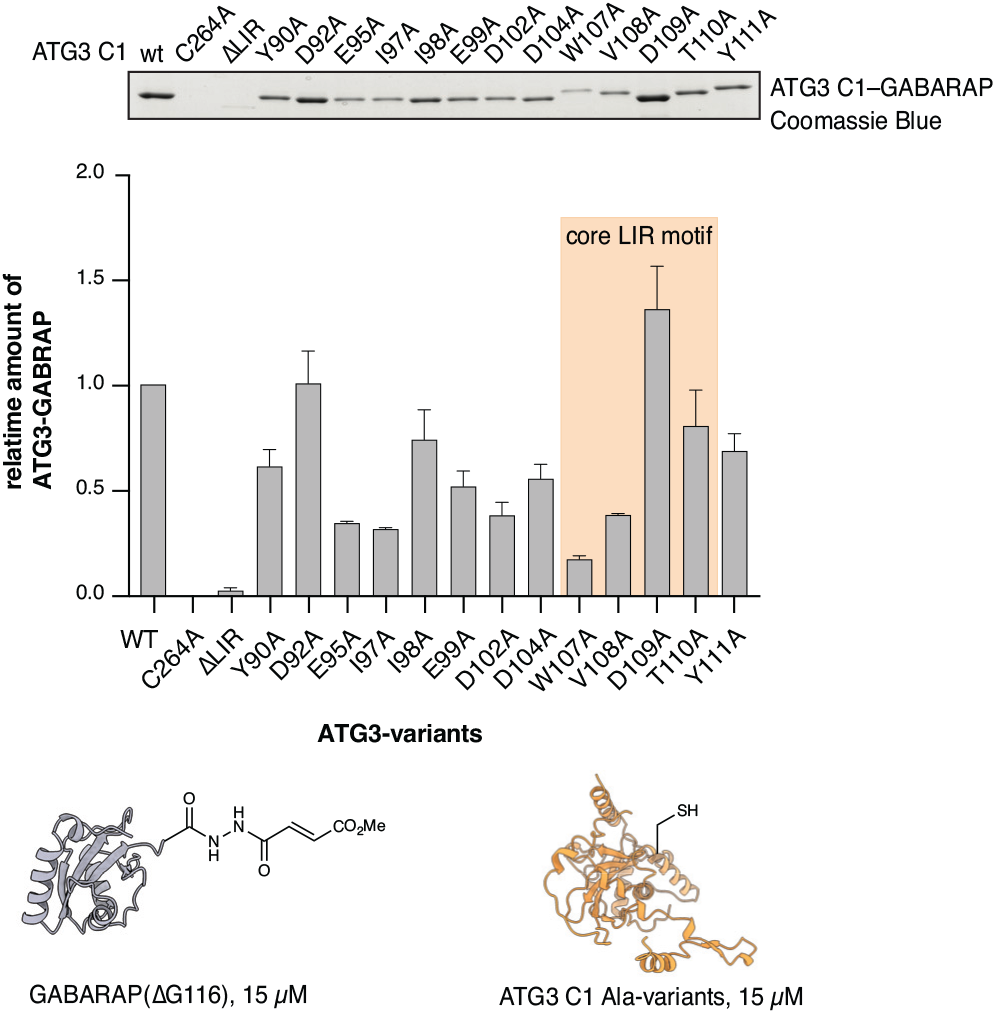
Dissection of interactions between GABARAP and LIR^ATG3^ by alanine scanning. GABARAP probe **2** (15 μM) was reacted with ATG3 C1 alanine-variants (15 μM) for 30 min. Coomassie Blue stained SDS-PAGE gel showing modification of ATG3 C1 variants with GABARAP methyl fumarate. The indicated ATG3 residues were mutated to alanine. Cross-linking efficiency was estimated by gel densitometry and normalized to the reaction with ATG3 C1. Data are presented as average values ± s.d. n=2 independent experiments. Source data and full-gel images are available in the Source Data File.

### LIR motif affects LC3 lipidation and thioester transfer *in cellulo*

LIR motifs are crucial to the function of various proteins involved in autophagy including selective autophagy receptors as well as enzymes involved in LC3 lipidation such as ATG7^40^ or ATG4^12^. The LIR motif in yeast ATG3 was shown to affect Atg8 lipidation *in vitro* and also affect the cytoplasm-to-vacuole pathway^25,41^.

To gauge the role of the LIR motif in human ATG3, we generated a homozygous HEK293T ATG3 knock-out (KO) cell line using CRISPR-Cas9. The knock-out was validated by immunoblotting against ATG3 and LC3A/B showing defective LC3 lipidation (Fig. 5a). Upon rescue with FLAG-ATG3 wild-type, LC3 was lipidated again as evidenced by the presence of the LC3–II band. However, expression of ATG3 C264A or ATG3 ΔLIR did not induce LC3 lipidation (Fig. 5b, Supplementary Figure 6). This finding indicates that the LIR motif in human ATG3 is a prerequisite for LC3 lipidation by ATG3.

**Fig. 5.**
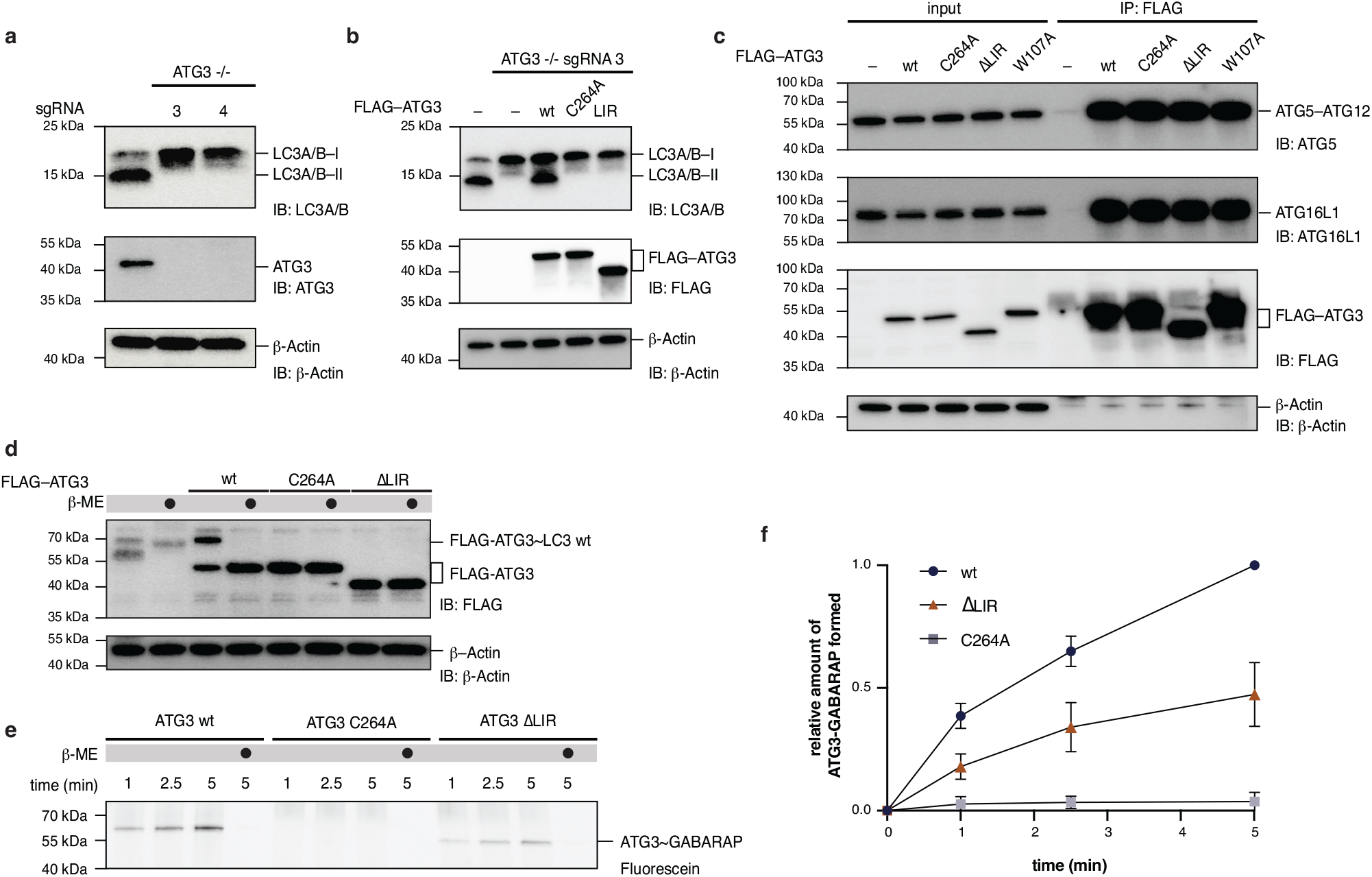
LIR^ATG3^ effects LC3 lipidation and thioester transfer. **a** Generation of homozygous knock-outs of HEK293T cells using CRISPR-Cas9 gene editing. Gene editing efficiency was validated by immunoblotting against ATG3 and LC3A/B. No lipidation was observed in knock-out cells. Analysis for two knock-out clones is shown. **b** Rescue with FLAG-ATG3 variants in HEK293T ATG3 –/– cells and autophagy induced by starvation for 1 h in the presence of chloroquine (40 *μ*M). Lipidation was assessed by immunoblotting against LC3A/B. Lipidation was observed upon re-expression of ATG3 wt but not ATG3 C264A or ΔLIR. As a positive control wild-type HEK293T cells were starved and analyzed in parallel to ATG3 –/– cells **c** Co-immunoprecipitation of FLAG-ATG3 variants (wt, C264A, ΔLIR, W107A). Interaction of ATG3 with ATG5–ATG12 and ATG16L1 was assessed by immunoblotting. No difference in pull-down efficiency was observed. **d** Detection of ATG3∼LC3A/B thioester *in cellulo*. FLAG-ATG3 variants were transiently expressed in HCT116 cells in which endogenous ATG3 was knocked down by CRISPR inhibition (CRISPRi). Autophagy was induced by starvation in the presence of chloroquine (40 *μ*M) for 1 h. Cell lysates were either prepared in the absence or in the presence of β-mercaptoethanol (β-ME). Samples containing reducing agent were boiled prior to SDS-PAGE analysis. The presence of a thioester intermediate was analyzed by anti-FLAG immunoblotting. ATG3∼LC3A/B thioester was only observed for ATG3 wt. **e** Pulse-chase assay of GABARAP transfer from ATG7 to ATG3. ATG7 was charged with fluorescein-labelled GABARAP. The reaction was quenched by addition of EDTA, and transfer initiated by addition of ATG3 variants (wt, C264A, ΔLIR). Reaction was monitored for the indicated time-points. Thioester-transfer was assessed by SDS-PAGE and in-gel fluorescence. **f** Quantification of in-gel fluorescence of **e**. Data are presented as average values ± s.d. and normalized to gel density of ATG3 wt 5 min. n=3 independent experiments. Source data and full-gel images for **a**-**e** are available in the Source Data File.

### How does LIR^ATG3^ influence ATG3’s activity?

We investigated the interaction of ATG3 with components of its enzymatic cascade by co-immunoprecipitation. We transiently expressed FLAG-ATG3 variants in HCT116 ATG3 knock-down cells generated using CRISPRi (Supplementary Fig. 7) and assessed their interaction with E3 complex subunits by anti-FLAG Co-IP.^42^ ATG3 wt, C264A, ΔLIR and W107A precipitated ATG5-ATG12 complex and ATG16L1 with similar efficiency (Fig 5c). Therefore, LIR^ATG3^ does not affect lipidation through defective binding to the E3 enzyme complex. Likewise, we performed pull-down of recombinantly expressed ATG3 variants with FLAG-ATG7 immobilized on anti-FLAG resin. No difference in pull-down efficiency was observed (Extended Data Figure 5). This indicates that LIR^ATG3^ or the lack thereof does not impact binding of ATG3 to ATG7 via its ATG7 interacting region (RIA7)^43^.

In order to assess the influence of LIR^ATG3^ on ATG3∼LC3 thioester formation, we expressed FLAG-ATG3 in HEK293T cells to analyze the presence of the ATG3∼LC3 thioester complex formation. Cell lysis under reducing and non-reducing conditions showed the presence of a thiol-sensitive band for wild-type ATG3. This band corresponds to the ATG3∼LC3 thioester complex, as no thioester was detected for ATG3 C264A (Fig. 5d). With ATG3 ΔLIR no higher MW thiol-sensitive band was detected. This observation is either due to inefficient trans-thioesterification between ATG7 and ATG3 or hydrolytic-instability of the ATG3 (ΔLIR)∼LC3/GABARAP complex. To further test the effect of LIR^ATG3^ on thioester transfer from ATG7 to ATG3 we performed a pulse-chase assay. Fluorescein-labelled GABARAP was charged onto ATG7, and thioester transfer initiated by addition of ATG3 variants. We observed a significant reduction in transfer-rate between ATG3 wt and ΔLIR (Fig 5e, f). Therefore, impeded thioester transfer at least partially explains the absence of ATG3∼LCA3/B upon LIR^ATG3^ or deletion.

## Discussion

Using a combination of chemical, biochemical, computational, and structural methods we have identified a previously unknown LIR motif in human ATG3. The core LIR motif, WVDT, is non-canonical and embedded in an unusual β-sheet conformation. Amino acids beyond the core LIR motif and the β-sheet conformation are involved in ATG3 binding to LC3/GABARAP. This uncommon β-sheet conformation for LIR motifs has only been reported for a few cases, including the FNIP tumor suppressor^31^, the *Legionella* effector protein RavZ^30^. A cursory search of reported LIR structures suggests that ALFY likely contains a β-sheet embedded LIR motif^44^. We provide conclusive evidence that the bent β-sheet conformation for LIR^ATG3^ is required for efficient binding to LC3/GABARAP and is not an artifact of co-crystallizing LIR peptide and LC3/GABARAP. This finding suggests that bent LIR conformations are more prevalent than currently appreciated. It is likely that a more wide-spread identification of this conformation has been precluded to this date by investigating LIR motifs in structural studies that were truncated in their N-terminal region and could not form the β-sheet conformation. This information will be useful in guiding bioinformatics methods to identify previously unidentified LIR motifs and extending the realm of LIR structures.

In addition, we show that LIR^ATG3^ is required for LC3 lipidation *in cellulo* and effects efficient thioester transfer from ATG7 to ATG3. While we observed no requirement of LIR^ATG3^ for interaction with either ATG5–ATG12, ATG16L1 or ATG7, the LIR motif influences the catalytic activity of the complex. ATG3’s flexible region contains not only the LIR motif but also two additional amino acid stretches required for ATG7 interaction, RIA7, and ATG12, RIA12. RIA7 and RIA12 partially overlap and are therefore mutually exclusive in their binding. It is probable that LIR^ATG3^ facilitates or influences binding of either region to their respective binding partner and therefore drives the lipidation reaction forward. It is also plausible that LIR^ATG3^ binds to and blocks the LIR-binding region of LC3/GABARAP from binding to the plethora of proteins bearing LIR motifs during its transfer from ATG7 to ATG3 to its substrate lipid. Interaction of LC3/GABARAP with one of these effector proteins during its transfer would likely negatively impact the efficiency of the lipidation reaction and stall efficient autophagosome expansion.

In conclusion, we describe a non-canonical LIR motif in human ATG3 crucial for its biological activity. Identification of this unusual LIR motif and its biological role provides new insights into the function of ATG3, one of the core proteins of autophagy.

## Methods

### Acylation of GABARAP and LC3A hydrazides

Proteins were diluted with acylation buffer (100 mM NaPhos, 50 mM NaCl, pH 3.0) to a concentration of 100 μM. The anhydride was dissolved in THF to a final concentration of 500 mM. The anhydride was added to the protein solution at the desired concentration (1 mM – 20 mM) and mixed. The reaction was allowed to proceed for ca. 5 min and analyzed by RP-HPLC or LC-MS analysis. The reaction was purified by dialysis at 4 ºC or by buffer exchange using desalting columns (Cytiva). Products could be used without further purification. For GABARAP(ΔG116)–NHNH α-chloroacetyl 200 equiv of α-chloroacetic acid anhydride were added. For GABARAP(ΔG116)–NHNH methyl fumarate 85 equiv methyl fumarate anhydride was added. For LC3A(ΔG120)–NHNH α-chloroacetyl 200 equiv α-chloroacetic acid anhydride was added. For LC3A(ΔG120)–NHNH methyl fumarate 20 equiv methyl fumarate anhydride was added.

### Generation of GABARAP and LC3A hydrazides from SUMO-GABARAP/LC3A–*Mxe*-GyrA–His_6_ fusions

SUMO-GABARAP/LC3A-*Mxe* GyrA-His_6_ fusion after Ni-NTA purification was dialyzed against intein cleavage buffer (25 mM HEPES, 500 mM NaCl, 1 mM EDTA pH 7.2 at 4 ºC). After dialysis hydrazine monohydrate (75 mM) and MESNa (10 mM) was added and the pH adjusted to 7.8. The reaction mixture was allowed to stand at room temperature and monitored using LC-MS analysis. Upon completion the solution was dialyzed against 25 mM HEPES, 500 mM NaCl pH 7.2 at 4 ºC. SENP1 (1 wt%) was added to cleave the SUMO-tag. The cleaved intein was removed using gravity Ni-NTA purification. The flow-through was collected and combined with resin washes. The combined fractions were dialyzed and further purified using cation exchange using a buffer of 25 mM Tris pH 8.0 and a gradient of 0–500 M NaCl in the same buffer over 25 column volumes. The product fractions were pooled, concentrated, and stored at –80 ºC until further use.

### Reaction of modified Ubls with E2s

Ubl probes (15 μM) were mixed with E2s (15 μM) in PBS. The mixture was incubated at 37 ºC in a water bath for 0.5-3 h. The reaction was quenched by addition of 2x Laemmli buffer and boiled at 95 ºC for 5 min. Samples were resolved by SDS-PAGE (8-16%, Bio-Rad) and visualized by Coomassie staining. If required, gel bands were quantified by gel densitometry using ImageLab (Bio-Rad).

### Co-crystallization of LIR^ATG3^ with GABARAP

GABARAP–OH in 25 mM Tris, 150 mM NaCl, pH 8.0 was concentrated using centrifugal filters (Amicon) to 730 μM ATG3 LIR peptide (Y90-H112) was resuspended in DMSO (50 mM) and 3 equivalents with respect to GABARAP were added. The reaction was dialyzed against 25 mM Tris, 150 mM NaCl, pH 8.0 at room temperature for 20 h. Crystallization was performed using hanging-drop vapour diffusion in 24-well crystallization flasks (Hampton Research). Protein solution (2 μL) was mixed with reservoir solution (2 μL) on siliconized cover-slips (Hampton Research). Crystallizations were performed at room temperature. Crystals were observed after 2 days. Crystals were obtained for 0.2 M NaOAc, 0.1 M sodium cacodylate, 30 vol% PEG 8,000, pH 6.5. Crystals were cryo-protected with 25 vol% glycerol and flash-frozen. Crystal diffraction was measured at the Swiss Light Source (Paul Scherrer Institute, Villigen, Switzerland) beamline X06DA (PXIII) equipped with the PILATUS 2M-F detector system (Dectris, Switzerland) at a wavelength of 1.0 Å, while the crystal was kept at 100 K. XDS software was used for data processing^45^. Molecular replacement using the GABARAP structure from PDB structure 6HB9 as the search model was performed using Phaser^32,46^. The asymmetric unit contained eight copies of the GABARAP–LIR^ATG3^ peptide complex. Refinement was performed iteratively using phenix.refine from the Phenix^47^ software package and Coot^48^. Protein structure visualization and analysis (RMSD, Coulombic potential, surface area) was performed using the UCSF ChimeraX software package^38^.

### *In cellulo* lipidation of LC3A/B

HEK293T ATG3 -/- cells were grown in a 6-well plate. pCMV-3xFLAG-ATG3 plasmids containing ATG3 wt, C264A or ΔLIR were transfected using Xtreme-Gene HP (Roche) in OptiMem. After 24 h media was changed to DMEM. After an additional 24 h the media was changed to HBSS supplemented with 40 μM chloroquine and incubated for 60 min. Cells were directly lysed using RIPA-buffer (50 mM Tris, 150 mM NaCl, 1 vol% Nonidet P-40, 0.5 wt% deoxycholate, 0.1 wt% SDS, pH 7.5) in the presence of protease inhibitors (Thermo-Fisher). Insoluble cell components were removed by centrifugation. Protein concentrations were measured by BCA according to the manufacturers protocol. Cell lysates were diluted to equal protein concentrations using RIPA-buffer. 10 μg of total protein was loaded and used for SDS-PAGE and Western bot analysis.

### Co-immunoprecipitation of ATG3

HCT116 ATG3 knock-down cells, generated by CRISPR inhibition, were grown in T25 flasks. pCMV-3xFLAG-ATG3 plasmids containing ATG3 wt, C264A or ΔLIR were transfected using Xtreme-Gene HP (Roche) in OptiMem. After 24 h media was changed to DMEM. After an additional 24 h the media was changed to HBSS supplemented with 40 μM chloroquine and incubated for 60 min. Cells were directly lysed using NP-40 buffer (50 mM Tris, 150 mM NaCl, 1 vol% nonidet-P40, pH 8.0) in the presence of protease inhibitors (Thermo-Fisher). Insoluble cell components were removed by centrifugation. An aliquot of lysate was removed for further analysis. The remaining cell lysate was incubated with Anti-FLAG M2 magnetic beads (20 μL, Sigma-Aldrich) for 2 h at 4 ºC. The resin was washed with NP40 lysis buffer and PBS. Proteins were eluted by addition of 4x non-reducing Laemmli buffer and boiling at 95 ºC for 5 min. β-mercaptoethanol was added to the eluted proteins. Input and elution were analyzed by SDS-PAGE and Western blot analysis.

### Analysis of *in cellulo* thioester formation

HEK293T ATG3 -/- cells were grown in a 6-well plate. pCMV-3xFLAG-ATG3 plasmids containing ATG3 wt, C264A or ΔLIR were transfected using Xtreme-Gene HP (Roche) in OptiMem. After 24 h media was changed to DMEM. After an additional 24 h the media was changed to HBSS supplemented with 40 μM chloroquine and incubated for 60 min. Cells were directly lysed using NP-40 buffer (50 mM Tris, 150 mM NaCl, 1 vol% nonidet-P40, pH 8.0) in the presence of protease inhibitors (Thermo-Fisher). Insoluble cell components were removed by centrifugation. Aliquots of cell lysate were removed and directly quenched by the addition of either 4x reducing Laemmli buffer or 4x non-reducing Laemmli buffer. Samples containing reducing Laemmli buffer were further boiled for 5 min at 95 ºC. Samples were analyzed by SDS-PAGE and Western blot analysis.

### Pulse-chase assay for GABARAP-thioester transfer to ATG3

Fluorescein-GABARAP (10 μM) was mixed with ATG7 (1 μM) in 25 mM Tris, 150 mM NaCl, pH 7.5. ATP-MgCl_2_ (5 mM) was added and the mixture was incubated at 4 ºC for 15 min. The solution was diluted 3-fold with 25 mM Tris, 150 mM NaCl, 50 mM EDTA, pH 7.5 and buffer exchanged using a Zeba-desalting column to 25 mM Tris, 150 mM NaCl, 50 mM EDTA, pH 7.5. The solution was added to recombinant ATG3 (0.5 μM) variants and incubated for 5 min. Aliquots were removed after 1, 2.5 and 5 min and mixed with either 4x non-reducing Laemmli buffer or 4x reducing Laemmli buffer. Samples were resolved using SDS-PAGE and analyzed by in-gel fluorescence and Stain-free visualization. ATG3∼GABARAP band was quantified by gel densitometry (Image-Lab, BioRad) and normalized to ATG3∼GABARAP band for wild-type ATG3 at 5 min.

## Data availability

Atomic coordinates and structure factors for the reported crystal structure have been deposited at the Protein Data Bank (PDB) with accession code 8AFI. Source data of uncropped and unprocessed gels are provided with this data.

## Acknowledgments

This work was supported by the Swiss National Science Foundation (Nos. 188634 and 310030_201160) and a European Research Council Synergy Grant under the European Union’s Horizon 2020 research and innovation programme (No. 856581─CHUbVi). JEC is supported by the NOMIS Foundation and the Lotte and Adolf Hotz-Sprenger Stiftung. We acknowledge the Molecular and Biomolecular Analysis Service of the Department of Chemistry and Applied Biosciences at ETH Zürich for mass spectrometry. We acknowledge the Protein Crystallization Center at University of Zürich and B. Blattmann for help with protein crystallization. We thank B. Schulman, S. M. Vos, L. Farnung, G Akimoto and R. Hofmann for helpful discussions. We thank the MX group for support at the Swiss Light Source (SLS) beamline and J. Beale for advice on data processing.

## Author Contributions

J.F. and J.W.B. conceived the project. J.F. designed and performed experiments, prepared materials, and analyzed data. M.M. prepared ATG3 knock-out cell-line. J.F. and J.R.L prepared ATG3 knock-down cell-line. J.F. and K.A.T. developed hydrazide modification. J.F. and R.B. performed structure determination by protein X-ray crystallography. J.W.B. designed experiments and analyzed data. J.F. and J.W.B. wrote the manuscript with help from all authors.

---

**Extended Data Fig. 1.**
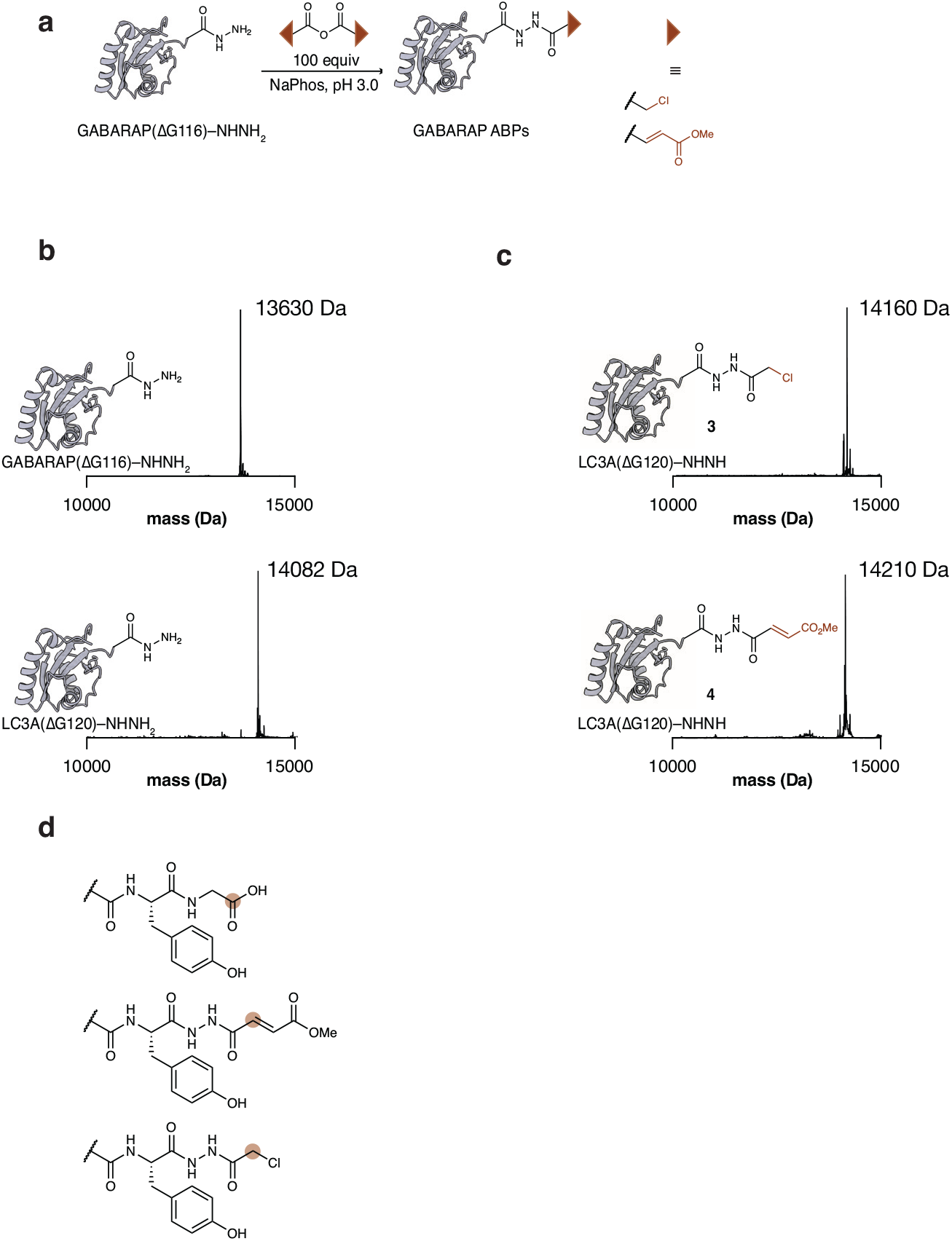
GABARAP and LC3A probe preparation by acyl hydrazide modification. **a**, Reaction scheme showing modification of GABARAP(ΔG116)–NHNH_2_ with symmetrical anhydrides at acidic pH. α-chloroacetyl and methyl fumarate probes were prepared. **b**, Deconvoluted mass-spectrum (ESI) of GABARAP(ΔG116)–NHNH_2_. Expected mass 13630 Da. Deconvoluted mass spectrum (ESI) of LC3A(ΔG120)–NHNH_2_. Expected mass 14082 Da. **c**, Deconvoluted mass spectrum (ESI) of LC3A(ΔG120)–NHNH α-chloroacetyl **3**. Expected mass 14158 Da. Deconvoluted mass spectrum (ESI) of LC3A(ΔG120)–NHNH methyl fumarate **4**. Expected mass 14210 Da. **d**, Alignment of native C-terminus of GABARAP and C-termini obtained by hydrazide modification. Site of modification is highlighted by brown circle.

**Extended Data Fig. 2.**
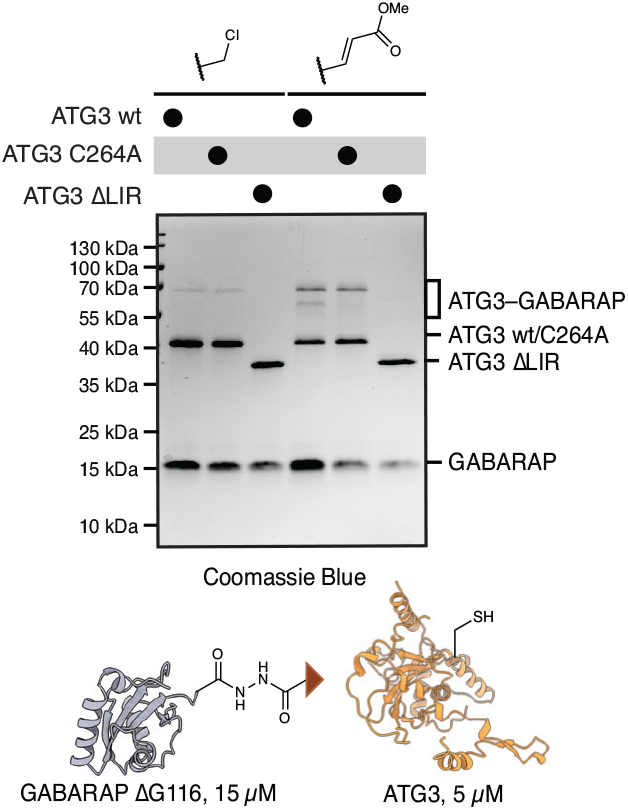
Modification of ATG3 wild-type is dependent on LIR^ATG3^. Coomassie Blue stained SDS-PAGE gel showing the reaction of GABARAP probes **1** and **2** (15 μM) with ATG3 wt, C264A and ΔLIR (5 μM). Source data and full-gel images are available in the Source Data File.

**Extended Data Fig. 3.**
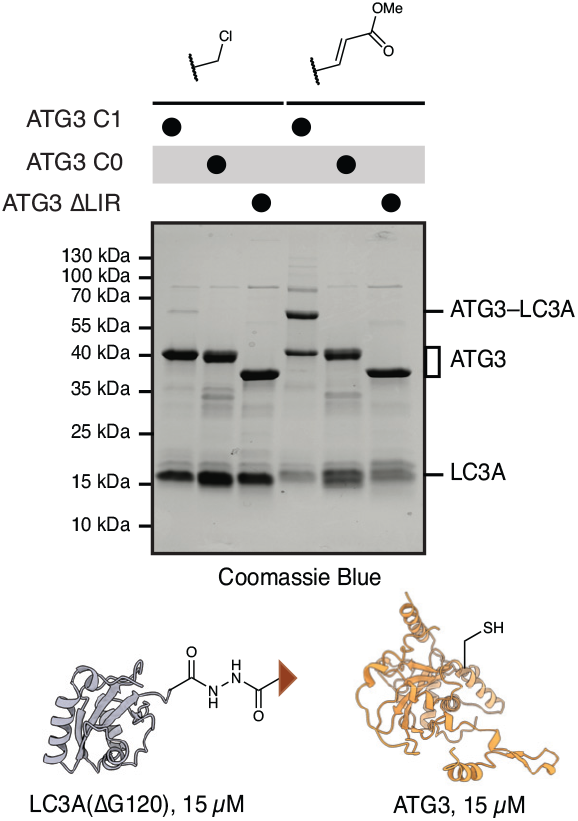
Modification of ATG3 with LC3A is dependent on LIR^ATG3^. Coomassie Blue stained SDS-PAGE gel showing the reaction of LC3A probes **3** and **4** (15 μM) with ATG3 C1, C0 and ΔLIR (15 μM). Source data and full-gel images are available in the Source Data File.

**Extended Data Fig. 4.**
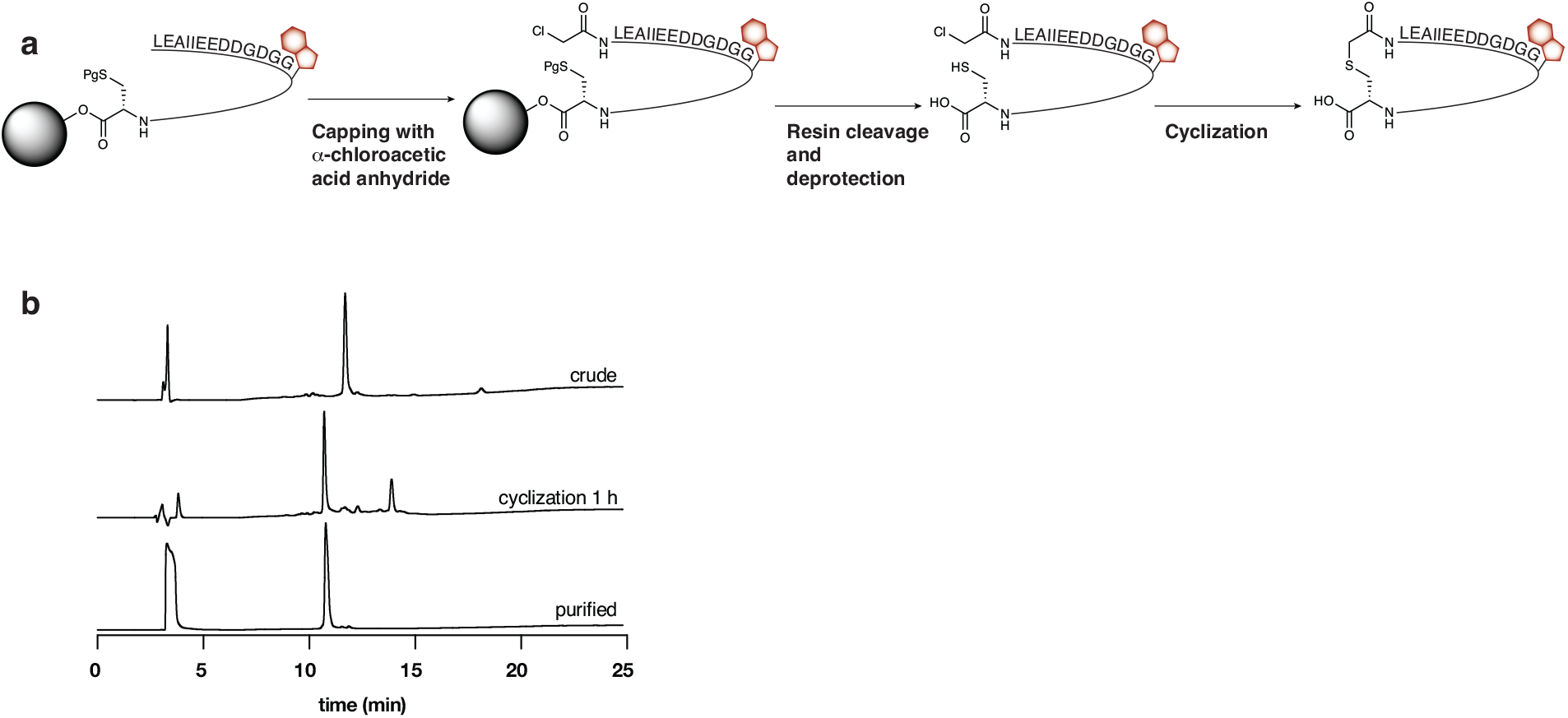
Preparation of cyclized LIR^ATG3^ peptide 6. **a**, Synthetic scheme describing the preparation of **6**. A cysteine residue is introduced at the C-terminus of the LIR motif. Following standard Fmoc-SPPS the LIR peptide is capped at its N-terminus with α-chloroacetic acid anhydride. Following resin cleavage, the peptide is cyclized by cysteine alkylation. **b**, Analytical RP-HPLC chromatograms showing the cyclization of the LIR peptide to provide **6**. Upper chromatogram shows the linear peptide prior to cyclization. Middle chromatogram shows the cyclization reaction after 1 h. Lower chromatogram shows purified peptide **6**.

**Extended Data Table 1.** Data collection and refinement statistics of co-crystal structure of GABARAP with LIR^ATG3^ peptide. Values in parentheses indicated values for high resolution shell.

| Parameters | 8AFI |
| --- | --- |
| **Data collection** |  |
| Wavelength (Å) | 1.0 |
| Space group | P1 |
| Cell dimensions |  |
| a, b, c (Å) | 62.80, 63.80, 73.90 |
| $\alpha$, $\beta$, $\gamma$ (°) | 64.50, 86.50, 70.30 |
| Resolution (Å) | 49.13 – 2.66 |
| R<sub>merge</sub> | 0.068 (1.13) |
| Completeness (%) | 94.7 |
| I/$\sigma$(I) | 14.19 (1.02) |
| CC<sub>1/2</sub> (%) | 99.9 (48.1) |
| Redundancy | 3.5 (3.4) |
| **Refinement** |  |
| Resolution (Å) | 47.14-2.66 (2.75-2.66) |
| Rwork/Rfree | 0.215/0.262 |
| No. reflections | 52724 |
| RMSD |  |
| bond lengths (Å) | 0.011 |
| bond angles (°) | 1.22 |
| No. atoms |  |
| Protein | 8485 |
| Ligand | 50 |
| water | 22 |
| B-factors |  |
| Protein | 90.67 |
| Ligand | 80.10 |
| water | 56.28 |
| Ramachandran plot |  |
| Favored | 96.87 |
| Allowed | 2.93 |
| Dis-allowed | 0.20 |

**Extended Data Fig. 5.**
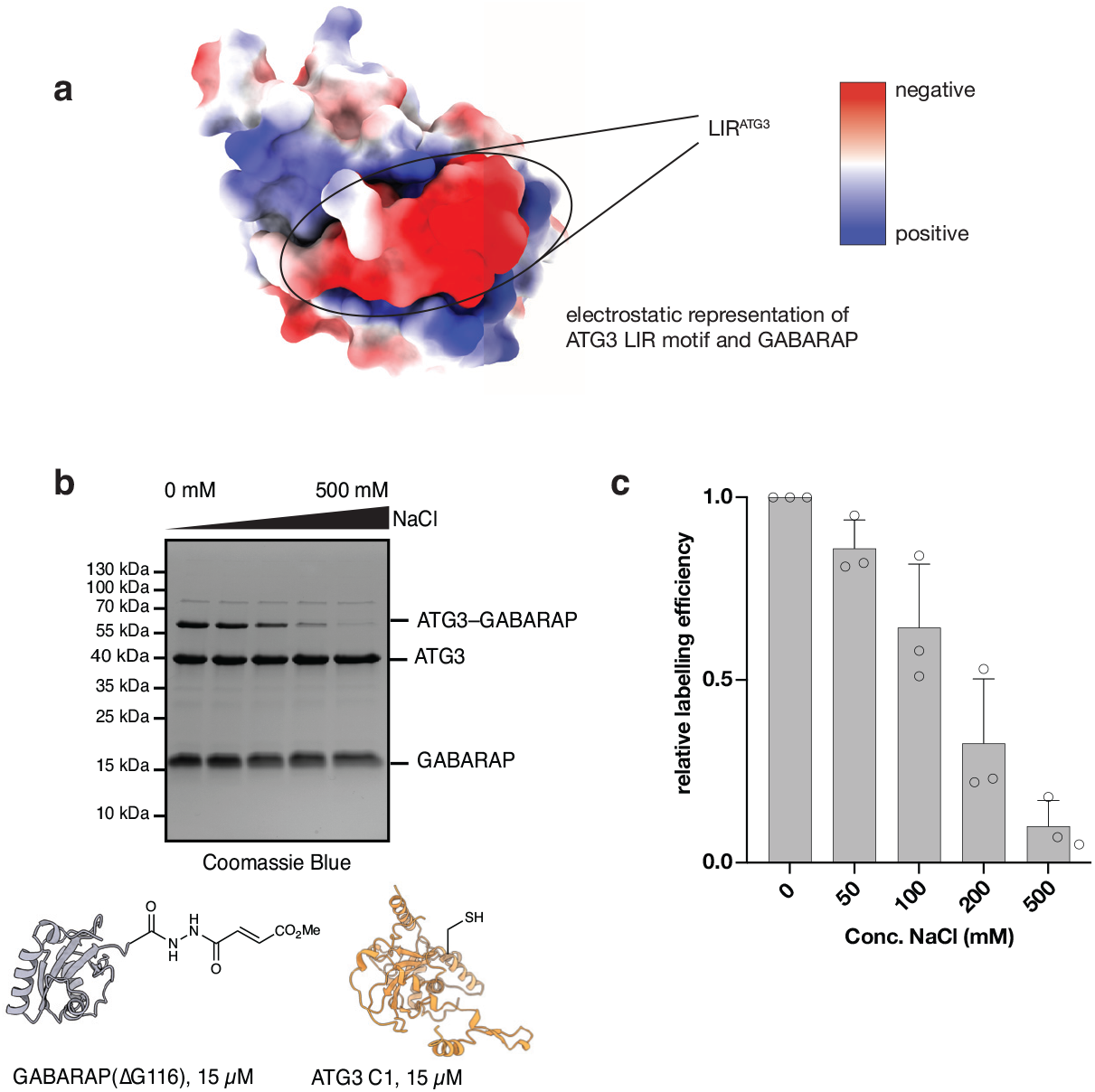
Interaction of ATG3 and GABARAP is salt-dependent. **a**, ColabFold prediction of ATG3-GABARAP complex (in Fig. 1c) shown with its Coulombic potential. Only LIR motif of ATG3 shown for clarity. LIR^ATG3^ is mostly negatively charged and binds to a strongly positively charged surface on GABARAP. **b**, Modification of ATG3 C1 with **2** in the presence of increasing concentrations of NaCl (0-500 mM). Coomassie Blue stained SDS-PAGE gel was analyzed by gel densitometry. **c**, Quantification of data shown in **b**. Data are presented as average values ± s.d. and normalized to gel density of ATG3 with 0 mM NaCl. n=3 independent experiments. Source data and full-gel images are available in the Source Data File.

**Extended Data Fig 6.**
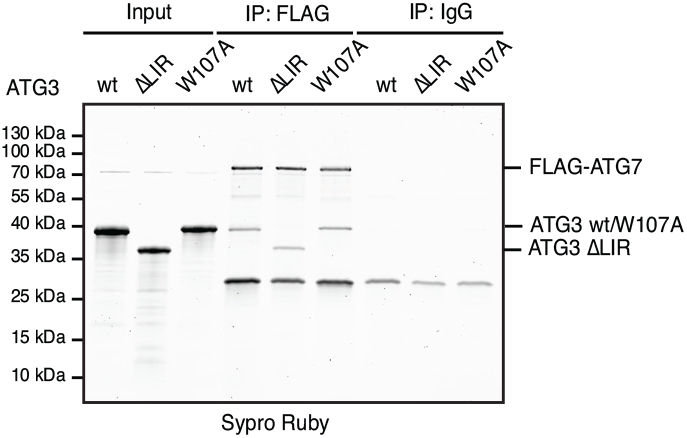
LIR^ATG3^ does not impact the interaction between ATG7 and ATG3. 3xFLAG-ATG7 was expressed in HCT116 cells with endogenous ATG3 knocked-down by CRISPRi and immobilized on Anti-FLAG beads following cell-lysis. Recombinant ATG3 variants (wt, ΔLIR, W107A) were incubated with immobilized ATG7. Bound proteins were eluted by boiling in non-reducing Laemmli buffer and analyzed by Sypro-Ruby staining of SDS-PAGE gel. Antibody fragments are visible after resin elution at 28 kDa. Source data and full-gel are available in the Source Data File.

## Notes

### Competing Interest Statement

The authors have declared no competing interest.

## References

1 Bento, C. F. et al. Mammalian Autophagy: How Does It Work? Annu Rev Biochem 85, 685–713, (2016).

2 Pankiv, S. et al. p62/SQSTM1 binds directly to Atg8/LC3 to facilitate degradation of ubiquitinated protein aggregates by autophagy. J Biol Chem 282, 24131–24145, (2007).

3 Novak, I. et al. Nix is a selective autophagy receptor for mitochondrial clearance. Embo Rep 11, 45–51, (2010).

4 Tumbarello, D. A. et al. The Autophagy Receptor TAX1BP1 and the Molecular Motor Myosin VI Are Required for Clearance of Salmonella Typhimurium by Autophagy. Plos Pathog 11, (2015).

5 Ichimura, Y. et al. A ubiquitin-like system mediates protein lipidation. Nature 408, 488–492, (2000).

6 Maruyama, T. et al. Membrane perturbation by lipidated Atg8 underlies autophagosome biogenesis. Nat Struct Mol Biol 28, 583–+, (2021).

7 Nguyen, T. N. et al. Atg8 family LC3/GABARAP proteins are crucial for autophagosome-lysosome fusion but not autophagosome formation during PINK1/Parkin mitophagy and starvation. J Cell Biol 215, 857–874, (2016).

8 Manil-Segalen, M. et al. The C. elegans LC3 Acts Downstream of GABARAP to Degrade Autophagosomes by Interacting with the HOPS Subunit VPS39 (vol 28, pg 43, 2014). Dev Cell 30, 110–110, (2014).

9 Weidberg, H. et al. LC3 and GATE-16/GABARAP subfamilies are both essential yet act differently in autophagosome biogenesis. Embo J 29, 1792–1802, (2010).

10 Noda, N. N. et al. Structural basis of target recognition by Atg8/LC3 during selective autophagy. Genes Cells 13, 1211–1218, (2008).

11 Birgisdottir, A. B., Lamark, T. & Johansen, T. The LIR motif - crucial for selective autophagy. J Cell Sci 126, 3237–3247, (2013).

12 Skytte Rasmussen, M. et al. ATG4B contains a C-terminal LIR motif important for binding and efficient cleavage of mammalian orthologs of yeast Atg8. Autophagy 13, 834–853, (2017).

13 Fan, S. J. et al. Inhibition of Autophagy by a Small Molecule through Covalent Modification of the LC3 Protein. Angew Chem Int Edit 60, 26105–26114, (2021).

14 Yang, A. M., Pantoom, S. & Wu, Y. W. Elucidation of the anti-autophagy mechanism of the Legionella effector RavZ using semisynthetic LC3 proteins. Elife 6, (2017).

15 Li, Y. T. et al. A semisynthetic Atg3 reveals that acetylation promotes Atg3 membrane binding and Atg8 lipidation. Nat Commun 8, (2017).

16 Norman, B. H. et al. Studies on the mechanism of phosphatidylinositol 3-kinase inhibition by wortmannin and related analogs. J Med Chem 39, 1106–1111, (1996).

17 Blommaart, E. F. C., Krause, U., Schellens, J. P. M., VreelingSindelarova, H. & Meijer, J. The phosphatidylinositol 3-kinase inhibitors wortmannin and LY294002 inhibit autophagy in isolated rat hepatocytes. Eur J Biochem 243, 240–246, (1997).

18 Sui, X. et al. Development and application of ubiquitin-based chemical probes. Chem Sci 11, 12633–12646, (2020).

19 Henneberg, L. T. & Schulman, B. A. Decoding the messaging of the ubiquitin system using chemical and protein probes. Cell Chem Biol 28, 889–902, (2021).

20 Hemelaar, J., Lelyveld, V. S., Kessler, B. M. & Ploegh, H. L. A single protease, Apg4B, is specific for the autophagy-related ubiquitin-like proteins GATE-16, MAP1-LC3, GABARAP, and Apg8L. J Biol Chem 278, 51841–51850, (2003).

21 Tolmachova, K. A., Farnung, J., Liang, J. R., Corn, J. E. & Bode, J. W. Facile Preparation of UFMylation Activity-Based Probes by Chemoselective Installation of Electrophiles at the C-Terminus of Recombinant UFM1. ACS Central Science, (2022).

22 Tolmachova, K. A., Farnung, J., Liang, J. R., Corn, J. E. & Bode, J. W. Facile Preparation of UFMylation Activity-Based Probes by Chemoselective Installation of Electrophiles at the C-Terminus of Recombinant UFM1. ACS Cent Sci 8, 756–762, (2022).

23 Huang, D. T. et al. Structural basis for recruitment of Ubc12 by an E2 binding domain in NEDD8’s E1. Mol Cell 17, 341–350, (2005).

24 Miura, T., Klaus, W., Gsell, B., Miyamoto, C. & Senn, H. Characterization of the binding interface between ubiquitin and class I human ubiquitin-conjugating enzyme 2B by multidimensional heteronuclear NMR spectroscopy in solution. J Mol Biol 290, 213–228, (1999).

25 Yamaguchi, M. et al. Autophagy-related Protein 8 (Atg8) Family Interacting Motif in Atg3 Mediates the Atg3-Atg8 Interaction and Is Crucial for the Cytoplasm-to-Vacuole Targeting Pathway. J Biol Chem 285, 29599–29607, (2010).

26 Jumper, J. et al. Highly accurate protein structure prediction with AlphaFold. Nature 596, 583–589, (2021).

27 Evans, R. et al. Protein complex prediction with AlphaFold-Multimer. bioRxiv, 2021.2010.2004.463034, (2022).

28 Mirdita, M. et al. ColabFold - Making protein folding accessible to all. bioRxiv, 2021.2008.2015.456425, (2022).

29 Pruneda, J. N., Stoll, K. E., Bolton, L. J., Brzovic, P. S. & Klevit, R. E. Ubiquitin in motion: structural studies of the ubiquitin-conjugating enzyme approximately ubiquitin conjugate. Biochemistry-Us 50, 1624–1633, (2011).

30 Kwon, D. H. et al. A novel conformation of the LC3-interacting region motif revealed by the structure of a complex between LC3B and RavZ. Biochem Bioph Res Co 490, 1093–1099, (2017).

31 Goodwin, J. M. et al. GABARAP sequesters the FLCN-FNIP tumor suppressor complex to couple autophagy with lysosomal biogenesis. Sci Adv 7, (2021).

32 Huber, J. et al. An atypical LIR motif within UBA5 (ubiquitin like modifier activating enzyme 5) interacts with GABARAP proteins and mediates membrane localization of UBA5. Autophagy 16, 256–270, (2020).

33 von Muhlinen, N. et al. LC3C, Bound Selectively by a Noncanonical LIR Motif in NDP52, Is Required for Antibacterial Autophagy. Mol Cell 48, 329–342, (2012).

34 Zorzi, A., Deyle, K. & Heinis, C. Cyclic peptide therapeutics: past, present and future. Curr Opin Chem Biol 38, 24–29, (2017).

35 Goto, Y. et al. Reprogramming the translation initiation for the synthesis of physiologically stable cyclic peptides. Acs Chem Biol 3, 120–129, (2008).

36 Bechtler, C. & Lamers, C. Macrocyclization strategies for cyclic peptides and peptidomimetics. Rsc Med Chem 12, 1325–1351, (2021).

37 Coyle, J. E., Qamar, S., Rajashankar, K. R. & Nikolov, D. B. Structure of GABARAP in two conformations: implications for GABA(A) receptor localization and tubulin binding. Neuron 33, 63–74, (2002).

38 Pettersen, E. F. et al. UCSF ChimeraX: Structure visualization for researchers, educators, and developers. Protein Sci 30, 70–82, (2021).

39 Wirth, M. et al. Molecular determinants regulating selective binding of autophagy adapters and receptors to ATG8 proteins. Nat Commun 10, 2055, (2019).

40 Noda, N. N. et al. Structural basis of Atg8 activation by a homodimeric E1, Atg7. Mol Cell 44, 462–475, (2011).

41 Sakoh-Nakatogawa, M., Kirisako, H., Nakatogawa, H. & Ohsumi, Y. Localization of Atg3 to autophagy-related membranes and its enhancement by the Atg8-family interacting motif to promote expansion of the membranes. Febs Lett 589, 744–749, (2015).

42 Kuma, A., Mizushima, N., Ishihara, N. & Ohsumi, Y. Formation of the approximately 350-kDa Apg12-Apg5.Apg16 multimeric complex, mediated by Apg16 oligomerization, is essential for autophagy in yeast. J Biol Chem 277, 18619–18625, (2002).

43 Ohashi, K. & Otomo, T. Identification and characterization of the linear region of ATG3 that interacts with ATG7 in higher eukaryotes. Biochem Bioph Res Co 463, 447–452, (2015).

44 Lystad, A. H. et al. Structural determinants in GABARAP required for the selective binding and recruitment of ALFY to LC3B-positive structures. Embo Rep 15, 557–565, (2014).

45 Kabsch, W. Xds. Acta Crystallogr D Biol Crystallogr 66, 125–132, (2010).

46 McCoy, A. J. et al. Phaser crystallographic software. J Appl Crystallogr 40, 658–674, (2007).

47 Afonine, P. V. et al. Towards automated crystallographic structure refinement with phenix.refine. Acta Crystallogr D Biol Crystallogr 68, 352–367, (2012).

48 Emsley, P. & Cowtan, K. Coot: model-building tools for molecular graphics. Acta Crystallogr D 60, 2126–2132, (2004).

