## Supplementary Information for "Human ATG3 contains a non-canonical LIR motif crucial for its enzymatic activity in autophagy"

### 1. Supplementary figures

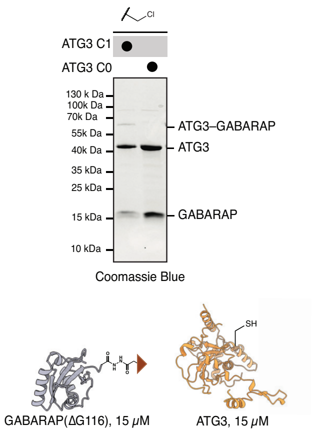

**Supplementary Figure 1. Reaction of GABARAP(ΔG116)–NHNH α-chloroacetyl probe 1 with ATG3 C1.** GABARAP probe **1** (15 μM) was incubated with ATG3 C1 or C0 (15 μM) for 1 h and analyzed by Coomassie-Blue staining and SDS-PAGE analysis. Cross-linking was only observed for ATG3 C1.

a

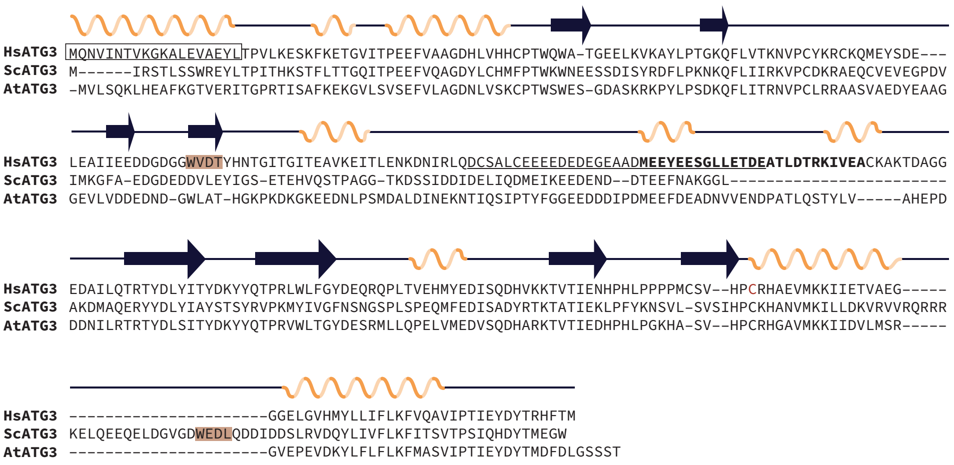

b

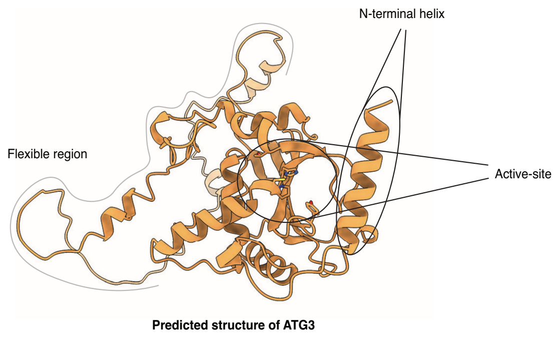

**Supplementary Figure 2. Sequence and structural elements of ATG3. a**, Sequence alignment of Homo Sapiens, Saccharomyces cerevisiae ATG3 and Arabidopsis thaliana ATG3. Sequences are aligned and structural elements of HsATG3 are shown. LIR$^{ATG3}$ is highlighted by brown box. N-terminal amphipathic helix is highlighted by box. RIA12 is highlighted by underlined text. RIA7 is highlighted by bold text. AIM$^{ATG3}$ in ScATG3 is highlighted by brown box. **b**, Predicted structure of HsATG3 using Colabfold. Structural elements are indicated.

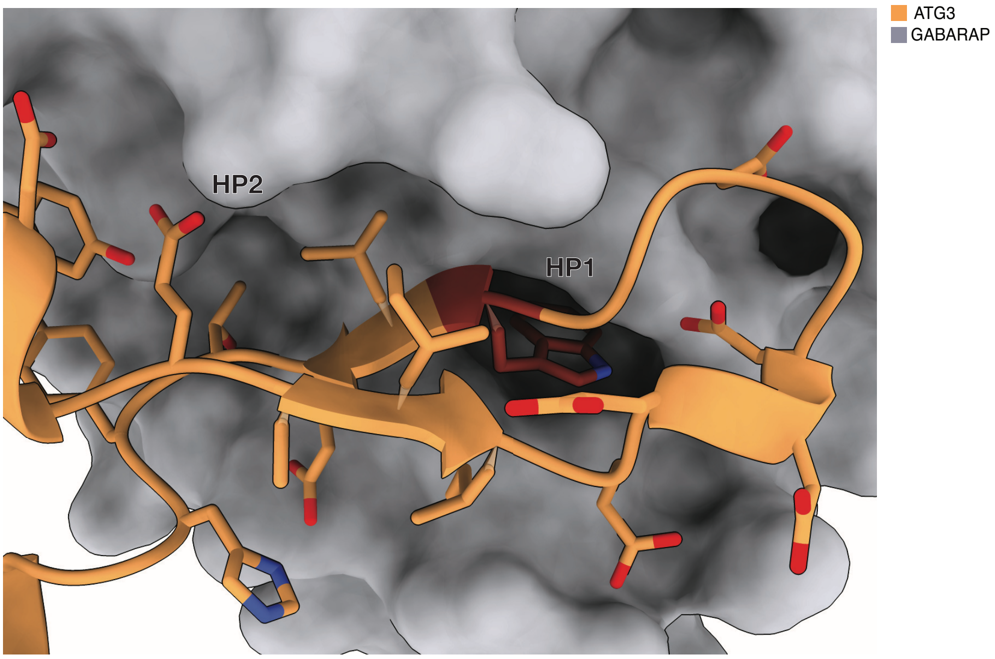

**Supplementary Figure 2. Zoom-in of predicted LIR motif in ATG3.** Zoom-in of predicted LIR motif in ATG3. Full-view shown in Figure 1c. W107 and T110 binding to hydrophobic pockets HP1 and HP2, respectively, are indicated. W107 is highlighted in red.

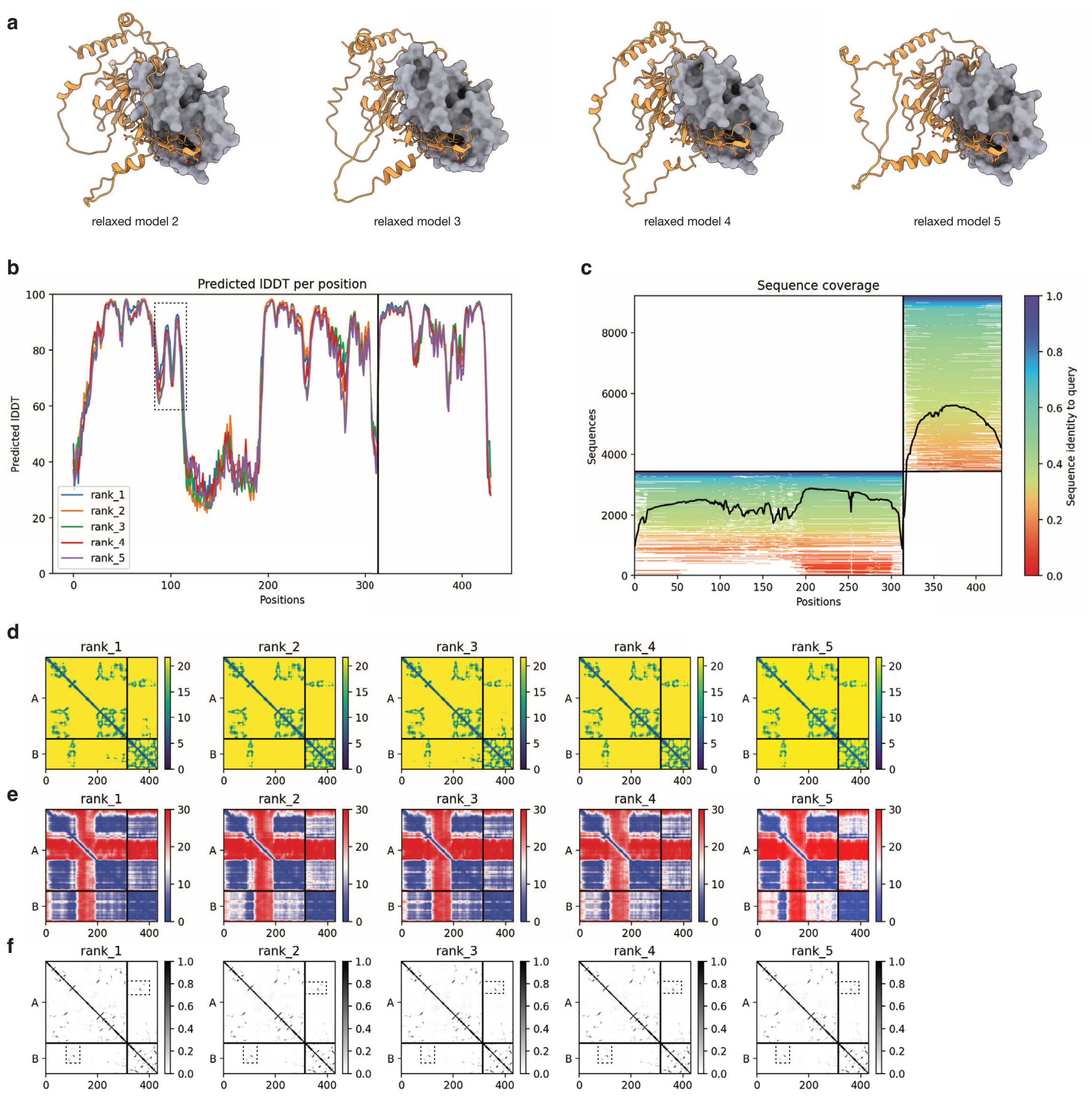

**Supplementary Figure 3. ColabFold prediction of GABARAP–ATG3 complex. a** Additional relaxed models predicted by ColabFold for GABARAP–ATG3 complex. Relaxed model 1 shown in Figure 1c. **b** Predicted lDDT values for relaxed models. LIR motif marked with dashed-box. **c** Multi-sequence alignment coverage for relaxed models. **d** Predicted distograms for all relaxed models. **e** Predicted errors of all relaxed models. **f** Predicted contacts of all relaxed models. LIR motif shown in dashed box.

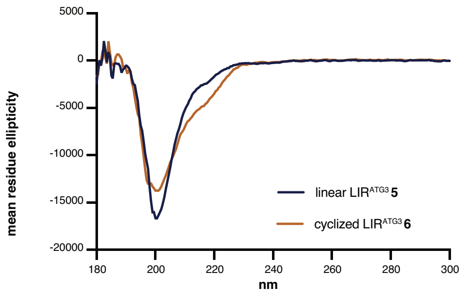

**Supplementary Figure 4. CD spectra of linear and cyclic LIR$^{ATG3}$ peptides 5 and 6.** CD spectra of LIR$^{ATG3}$ peptides **5** and **6** were recorded in 20 mM KPhos buffer (pH 7.2) at a concentration of 700 µM. Cyclized LIR$^{ATG3}$ peptide **6** shows a marked shift to a more organized structure as indicated by a decrease of the band intensity at 200 nm and an increase in band intensity at 215 nm.

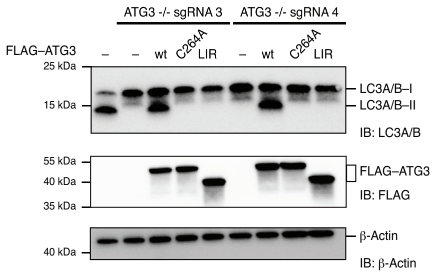

**Supplementary Figure 5. Reconstitution of LC3 lipidation with ATG3 variants in HEK293T ATG3 –/–.** FLAG-ATG3 variants were re-expressed in HEK293T ATG3 $^{-/-}$ cells and autophagy induced by starvation for 1 h in the presence of chloroquine (40 $\mu$M). Lipidation was assessed by immunoblotting against LC3A/B. Lipidation was observed upon re-expression of ATG3 wt but not ATG3 C264A or ΔLIR. As a positive control wild-type HEK293T cells were starved and analyzed in parallel to ATG3 –/– cells. Data for ATG3 –/– knockout with sgRNA 3 shown in Figure 6b.

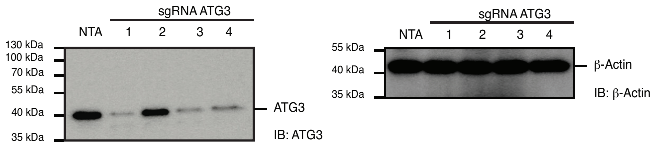

**Supplementary Figure 6. Western blot analysis of HCT116 ATG3 knock-down cells.** Western blot analysis anti-ATG3 and anti-β-actin. sgRNAs 1,3 and 4 cause robust knockdown of ATG3 compared to non-targeting sgRNA.

#### 2. General information

##### 2.1. Reagents and solvents

Chemical reagents were purchased from Sigma Aldrich (Buchs, Switzerland), Acros Organics (Geel, Belgium), TCI Europe (Zwijndrecht, Belgium) and used without further purification. $CH_2Cl_2$, DMF, EtOH, THF and MeOH were purchased from Fisher Scientific (Geel, Belgium) and Sigma Aldrich (Buchs, Switzerland) and used without further purification (reagent or HPLC grade). Milli-Q water was obtained from a Millipore purification system. Q5 High-Fidelity DNA Polymerase, PCR reagents, Gibson Assembly Master Mix, were purchased from New England BioLabs (Ipswich, MA, USA). DNase I was obtained from Roche Diagnostics GmbH (Mannheim, Germany). DNA purification kits were purchased from Fisher Scientific (Geel, Belgium) and Zymo Research (Irvine, CA, USA). Lysozyme (22500 U/mg) was obtained from Axon Lab AG (Baden, Switzerland). Kanamycin sulfate and ampicillin were obtained from AppliChem GmbH (Darmstadt, Germany). Ni-NTA agarose resin was obtained from Qiagen GmbH (Hilden, Germany). Bradford protein assays were performed using Protein Asaay Dye Reagent Concentrate from Bio-Rad. Dialysis tubing was obtained from Thermo Fisher Scientific (Waltham, MA, USA). Amicon Ultra centrifugal filters were purchased from Merck (Darmstadt, Germany), VivaSpin 500 centrifugal concentrators were purchased from Satorius Stedim

Lab (Stonehoude, UK), PD MiniTrap desalting columns were purchased from Cytiva. All buffers were prepared using Mili-Q water, sterile filtered (0.2 µm membrane filter), pH was adjusted at the temperature the buffer was used. Oligonucleotide synthesis and sequencing was carried out by Microsynth AG (Balgach, Switzerland). Synthetic genes were ordered from ATG Biosynthetics (Merzhausen, Germany). Cell culture reagents including Dulbecco's modified eagle medium (DMEM), OptiMEM, phosphate-buffered saline (PBS), trypsin-EDTA, HBSS and Fetal Bovine Serum (FBS) were purchased from Thermo Fisher Scientific. Transfection reagent Xtreme gene HP was obtained from Roche. ATG7 was purchased form R&D Systems (Boston, USA).

Fmoc-amino acids with suitable side-chain protecting groups, HATU (1-[bis(dimethyl-amino)methylene]-1H-1,2,3-triazolo[4,5-b]pyridinium 3-oxide hexafluorophosphate) were purchased from Merck-Milipore and ChemImpex. HPLC grade $CH_3CN$ from Sigma-Aldrich was used for analytical and preparative HPLC purification. Trifluoroacetic acid for HPLC analytical and preparative HPLC purification was purchased from ABCR. DMF (> 99.8%) from Sigma-Aldrich and N-methylpyrrolidine from ABCR were directly used without further purification for solid phase peptide synthesis. Other commercially available reagents and solvents were purchased from Sigma-Aldrich (Buchs, Switzerland), Acros Organics (Geel, Belgium) and TCI Europe (Zwijndrecht, Belgium).

#### 2.2. Mass spectrometric characterization

High-resolution mass spectra were recorded by the Molecular and Biomolecular Analysis Service (MoBiAS) at ETH Zurich with a Bruker maXis instrument (ESI-MS measurements) equipped with an ESI source and a Q-TOF detector. Reaction monitoring was performed on a Bruker microFLEX instrument (MALDI-TOF) using 4-hydroxy-α-cyanocinnamic acid as the matrix. LC-MS analysis was performed on a Waters Xevo G2-XS QTof. Mass analysis was performed using MassLynx. Sample peaks were integrated and deconvoluted using MaxEnt1.

#### 2.3. RP-HPLC analysis and purification

Peptides were analyzed and purified by reverse phase high performance liquid chromatography (RP-HPLC) on JASCO analytical and preparative instruments equipped with dial pump, mixed and in-line degasser, a variable wavelength UV detector (simultaneous detection of the eluent at 220 nm, 254 nm and 301 nm) and a Rheodyne injector with a 200 µL or 10 mL injection loop. Columns were heated to 60 °C using a Jetstream 2 column heater (analytical) or a $H_2O$ water bath (preparative). The mobile-phases for RP-HPLC were Milipore-$H_2O$ containing 0.1% (v/v) TFA and HPLC grade $CH_3N$ containing 0.1% (v/v) TFA. Analytical HPLC was performed at a flow rate of 1 mL/min. Analytical columns used were Shiseido Capcell Pak C18 (5 µm, 4.6 mm I.D. x 250 mm) or Agilent Eclipse XDB-C8 (5 µm, 4.6 mm x 150 mm). Preparative HPLC was performed on Shiseido Capcell Pak MGIII (5 µm, 20 mm I.D. x 250 mm) at a flow rate of 40 mL/min.

General *analytical* HPLC methods:

- flow 1 mL/min, isocratic 10% $CH_3CN$ for 3 min, then gradient from 5% to 95% $CH_3CN$ in 14 min

General *preparative* HPLC methods:

- flow 40 mL/min, isocratic 5% $CH_3CN$ for 5 min, then gradient from 5% to 65% $CH_3CN$ in 28 min.

#### 2.4. Fast protein liquid chromatography (FPLC)

Chromatographic protein purification was performed on an Äkta Pure system (GE Healthcare) at 4 °C. Buffers were prepared and pH-adjusted at 4 °C. Buffers were filtered (2 µm) and degassed prior to use. Ion exchange chromatography was performed using MonoS 5/50 GL, or MonoQ 5/50 GL. Size exclusion chromatography was performed on a Superdex 75 Increase 10/300 G, or HiLoad 16/600 Superdex 200. Protein elution was monitored at 280 nm and acquired using Unicorn (6.3.2.89).

#### 2.5. Solid-phase peptide synthesis

Loading of amino acids on solid support was performed as followed:
C-terminal carboxylic acid: The amino acid (1.20 equiv of desired loading) was dissolved in $CH_2Cl_2$ (200 mM). NMM (2 equiv) was added to the solution. The solution was given to preswollen chloro-trityl resin and shaken for 1h. The resin was washed with $CH_2Cl_2$ and DMF. Remaining chloro-trityl moieties were capped with $CH_2Cl_2$/MeOH/NMM (17:2:1, v:v:v) for 1 min. The capping step was repeated once. The resin was washed with $CH_2Cl_2$ and DMF. The resin was dried using a $N_2$ stream. The resin was stored at 4 °C.

Peptides were synthesized on a Multisyntech Syro I parallel synthesizer using Fmoc-SPPS chemistry. The following Fmoc amino acids with side-chain protection groups were used: Fmoc-Ala-OH, Fmoc-Arg(Pbf)-OH, Fmoc-Asn(Trt)-OH, Fmoc-Asp(OtBu)-OH, Fmoc-Cys(Trt)-OH, Fmoc-Gln(Trt)-OH, Fmoc-Glu(OtBu)-OH, Fmoc-Gly-OH, Fmoc-His(1-Trt)-OH, Fmoc-Ile-OH, Fmoc-Leu-OH, Fmoc-Lys(Boc)-OH, Fmoc-Lys(Alloc)-OH Fmoc-Nle-OH, Fmoc-Phe-OH, Fmoc-Pro-OH, Fmoc-Ser(tBu)-OH, Fmoc-Thr(tBu)-OH, Fmoc-Trp(Boc)-OH, Fmoc-Tyr(tBu)-OH, Fmoc-Val-OH.

General methods on Multisyntech Syro I parallel synthesizer:

- Amino acids were dissolved in DMF to a concentration of 0.5 M. HATU was dissolved in DMF to a concentration of 0.5 M. DIPEA was dissolved in NMP to a concentration of 2 M. Amino acid, HATU and DIPEA are mixed to a final concentration of 0.2 M, 0.2 M and 0.4 M, respectively, and added to the resin. The resin was agitated for 45 min. Coupling steps were repeated once.
- Capping was performed with acetic anhydride. 20 vol% acetic anhydride in DMF was mixed with 2 M DIPEA at a ratio of 3:2 and added to the resin. The resin was agitated for 5 min. The capping step was repeated once.
- Fmoc deprotection was performed with 20 vol% piperidine in DMF for 10 min. The deprotection step was repeated once.

##### 2.6. Gel electrophoresis

Sodium dodecyl sulfate-polyacrylamide gel electrophoresis (SDS-PAGE) was carried out on a Mini-PROTEAN Tetra Cell system (Bio-Rad) or a Criterion Cell connected to the PowerPac Basic (Bio-Rad) power supply. Reducing samples were treated with sample buffer (Laemmli 2x or 4x Concentrate, Sigma Aldrich), heated at 95 °C for 5 min and used for separation. A 10-180 kDa pre-stained protein ladder (Thermo Fisher) was applied to at least one well of each gel (4 µL). Samples were separated on 8-16% gradient Mini-PROTEAN TGX Precast gels (Bio-Rad) for 30 minutes at 200 V or 8-16% gradient Criterion TGX Precast gels (Bio-Rad) for 60 min at 180 V at 4 °C. Gels were stained with Coomassie (0.1% Coomassie Brilliant Blue R, 40% MeOH, 10% acetic acid) with agitation for 1h and subsequent destaining (40% MeOH, 10% acetic acid), or with Sypro-Ruby (Bio-Rad) according to the manufacturers procedure. Destained gels were imaged on a Bio-Rad Molecular Imager Pharos FX (Coomassie Blue). Gels were quantified by gel densitometry using ImageLab (Bio-Rad). Gels were cropped in Adobe Illustrator. Full-gels are shown in the Source Data.

##### 2.7. Western blotting

Proteins were electrophoretically transferred from resolved gels to low-fluorescence PVDF membranes using TransBlot (Bio-Rad). For blots anti FLAG, ATG5, ATG16L1 and β-actin, transfer was performed using Mixed MW method (7 min, 1.3 A). For blots anti LC3A/B transfer was performed using Low MW method (5 min, 1.3 A). Blots were blocked with 5 wt% dry skim milk powder in TBS-T for 2 h at room temperature. Blots were washed with TBS-T and then incubated with antibodies at indicated dilutions in TBS-T with 1 wt% BSA for 18 h at 4 °C. Blots were exhaustively washed with TBS-T and incubated with secondary antibody at indicated dilutions in TBS-T with 1 wt% BSA for 1.5 h at room temperature. Blots were exhaustively washed and incubated with Clarity Western ECL (Bio-Rad) and visualized using Bio-Rad Molecular Imager Pharos FX (Chemiluminescence). Blots were cropped in Adobe Illustrator. Full-blots are shown in the Source Data.

#### 2.8. Antibodies

| Plasmid | Supplier | Identifier | Dilution |
|---|---|---|---|
| Anti-FLAG | Thermo-Fisher | #MA1-91878 | 1:1,000 |
| Anti-LC3/AB | Cell-Signalling | #12741 | 1:1,000 |
| Anti-ATG5 | Cell-Signalling | #D5F5U | 1:1,000 |
| Anti-ATG16L1 | Cell-Signalling | #D6D5 | 1:1,000 |
| Anti-β-Actin | Thermo-Fisher | #BA3R | 1:2,000 |
| Anti-Mouse | Thermo-Fisher | #62-6520 | 1:3,000 |
| Anti-Goat | Cell-Signalling | #91196 | 1:3,000 |

**Supplementary Table S1** Antibodies used in this study.

#### 2.9. Circular dichroism spectroscopy

CD spectra were recorded on a JASCO J-1500 CD spectrometer in a 0.1 cm cuvette. Samples were measured in potassium phosphate (20 mM, pH 7.2) at 20 °C. Data were recorded between 180-300 nm with a measuring speed of 20 nm/min. Measurements were repeated (5x) and averaged. Background spectrum was recorded and subtracted from measured spectra. Data were converted to mean residue ellipticity.

#### 2.10. Gene constructs

Ubiquitin (P0CG48) was synthesized by ATG-Biosynthetics, codon-optimized for *E. coli*. SUMO2 (P61956, Addgene plasmid #53142) was a gift from Dieter Willibold. GABARAP was a gift from Dieter Willibold (O95166, Addgene plasmid #73948), LC3A was a gift from Dieter Willibold (Q9H492, Addgene plasmid #73946), Ube2K C170S (P61086, Addgene plasmid #18892) was a gift from Cecile Pickart, ATG3 was a gift from Robin Ketteler (Q9NT62, Addgene plasmid #129296), SENP1 cat. domain (Q9P0U3, Addgene plasmid #16356) was a gift from Guy Salvesen, ATG7 was a gift from Eiki Kominami and Isei Tanida (O95352, Addgene plasmid #87867). Ubiquitin was subcloned into *Mxe*-GyrA-$His_6$ pET28a (Merck Novagen) using Gibson assembly. GABARAP and LC3A were subcloned into *Mxe*-GyrA-$His_6$ pET28a (Merck Novagen) with an N-terminal SUMO2 solubilizing tag to access protein hydrazides. GABARAP and LC3A were subcloned into pOPINS to express GABARAP and LC3 wild-type proteins. ATG3 was subcloned into pOPINS using Gibson assembly. Mutations were introduced using inverse PCR. All sequences were confirmed by DNA Sanger sequencing (Microsynth, Plasmidsaurus).

| Plasmid | Purpose |
|---|---|
| pET28a Ub(ΔGG)–*Mxe* GyrA-His6 | Ub(ΔGG) intein expression to generate Ub(ΔGG)-$NHNH_2$ |
| pET28a SUMO2-GABARAP(ΔG116)–*Mxe* GyrA-His6 | SUMO-GABARAP(ΔG116) intein expression to generate GABARAP(ΔG116)-$NHNH_2$ |
| pET28a SUMO2-LC3A(ΔG120)–*Mxe* GyrA-His6 | SUMO-LC3A(ΔG116) intein expression to generate LC3A(ΔG116)-$NHNH_2$ |
| pET28 His6-TEV-Ube2K | Ube2K expression |
| pOPINS His6-SUMO-ATG3 | ATG3 wt expression |
| pOPINS His6-SUMO-ATG3 C264A | ATG3 C264A expression |
| pOPINS His6-SUMO-ATG3 ΔLIR | ATG3 ΔLIR (Δ90-112) expression |
| pOPINS His6-SUMO-ATG3 C1 | ATG3 expression containing only C264 |
| pOPINS His6-SUMO-ATG3 C1 C264A | ATG3 expression containing no cysteines |
| pOPINS His6-SUMO-ATG3 C1 ΔLIR | ATG3 ΔLIR (Δ90-112) expression containing only C264 |
| pCMV-3XFLAG-ATG3 | Mammalian expression of 3x-FLAG ATG3 |
| pCMV-3XFLAG-ATG3 C264A | Mammalian expression of 3x-FLAG ATG3 C264A |
| pCMV-3XFLAG-ATG3 ΔLIR | Mammalian expression of 3x-FLAG ATG3 ΔLIR |
| pCMV-3XFLAG-ATG7 | Mammalian expression of 3x-FLAG ATG7 |

**Supplementary Table S2** Table showing plasmids used in this study. Plasmids for Ala-screen (Fig. 4) are derived from plasmid pOPINS His6-SUMO-ATG3 C1.

##### 2.11. Protein expression

Chemically competent BL21 (DE3) cells (Ub, Ube2K) or Rosetta-CodonPlus (DE3)-RIL (GABARA, LC3A, ATG3, SENP1) were heat-shock transformed with the plasmids and single colonies were used to inoculate overnight precultures in selective lysogeny broth (LB) Miller medium. For Ube2K, ATG3, Ubiquitin-*Mxe*-GyrA, preculture was diluted 1:200 with fresh selective LB Miller medium, cultures were grown in baffled shake flasks at 37 °C. Ub-*Mxe*-GyrA fusion proteins were grown to $OD_{600}$ 0.6. The temperature was reduced to 25 °C and protein expression induced by addition of isopropyl β-D-1-thiogalactopyranoside at a final concentration of 0.5 mM. Expression was allowed to proceed for 18 h. Cells were harvested (4,500 g, 35 min, 4 °C) and resuspended in intein lysis buffer (20 mM Tris, 200 mM NaCl, pH 7.2). Cells were lysed by addition of lysoszyme, DNase I. Cells were further lysed by sonication and suspension clarified by centrifugation (20,000 g, 35 min, 4 °C). Intein fusion proteins were purified by gravity Ni-NTA purification. Resin was washed with excess intein lysis buffer and eluted using intein elution buffer (20 mM Tris, 200 mM NaCl, 300 mM imidazole pH 7.2). The isolated fusion protein was dialyzed against intein cleavage buffer (see below). Ube2K was expressed and purified as previously reported.[1] For SUMO-GABARAP/LC3– *Mxe*-GyrA preculture was diluted 1:200 with fresh autoinduction media. Cells were grown to $OD_{600}$ 1.0, the temperature was reduced to 18 h and cells were grown for an additional 36 h. Cell lysis was performed as for ubiquitin. ATG3 was grown to grown to $OD_{600}$ 0.6. Temperature was reduced to 25 °C and protein expression induced by addition of isopropyl β-D-1-thiogalactopyranoside at a final concentration of 0.5 mM. Cultures were grown for 18 h. Cells were harvested (4,500 g, 35 min, 4 °C) and resuspended in intein lysis buffer (50 mM Tris, 350 mM NaCl, 10 vol% glycerol, 30 mM imidazole, pH 7.4). Cells were lysed by addition of lysoszyme, DNase I and DTT. Cells were further lysed by sonication and suspension clarified by centrifugation (20,000 g, 35 min, 4 °C). Following Ni-NTA purification. Protein was diluted with 25 mM HEPES pH 7.0 and SENP1 was added to cleave $His_6$-SUMO. Proteins were further purified by anion exchange chromatography using a MonoQ column (Cytiva) on an Äkta Pure (GE Healthcare) with a gradient of 25 mM HEPES pH 7.0 to 25 mM HEPES, 500 mM NaCl, pH 7.0 over 25 column volumes. Product fractions were pooled and concentrated. For-SUMO2-GABARAP/LC3A wild-tyoe, proteins were expressed according to protocol of intein fusion proteins. After Ni-NTA, SENP1 (1 wt%) was added and protein diluted with 25 mM HEPES pH 7.0. Following complete SUMO-cleavage, GABARAP/LC3A were further purified by anion exchange chromatography using a MonoQ column (Cytiva) on an Äkta Pure (GE Healthcare) with a gradient of 25 mM HEPES pH 7.0 to 25 mM HEPES, 500 mM NaCl, pH 7.0 over 25 column volumes. Protein concentrations were estimated by absorption at 280 nm using extinction coefficients calculated using the online tool ProtParam (https://web.expasy.org/protparam). Proteins were aliquoted, flash frozen in liquid $N_2$ and stored at –80 °C until use.

##### 2.12. Mammalian cell culture

HEK293T and HCT116 cells were obtained from ATCC. Cells were cultured in DMEM supplemented with FBS and penicillin/streptomycin. Cells were seeded at appropriate densities in 6-well, T25 or T75 flasks. Cells were maintained at 37 °C with 5 vol% $CO_2$.

#### 3. General methods

##### 3.1. Acylation of hydrazides with anhydrides

Proteins were dissolved or diluted with acylation buffer (100 mM NaPhos, 50 mM NaCl, pH 3.0) to a concentration of 100 μM. The anhydride was dissolved in THF or DMF to a final concentration of 500 mM. The anhydride was added to the desired concentration (1 mM – 20 mM) and mixed. The reaction was allowed to proceed for ca. 5 min. The reaction was analyzed by RP-HPLC or LC-MS analysis. The reaction was purified by dialysis at 4 °C or by buffer exchange using desalting columns (Cytiva). Products could be used without further purification.

For GABARAP(ΔG116)–NHNH $\alpha$-chloroacetyl 200 equiv of $\alpha$-chloroacetic acid anhydride were added.

For GABARAP(ΔG116)–NHNH methyl fumarate 85 equiv methyl fumarate anhydride was added.

For LC3A(ΔG120)–NHNH $\alpha$-chloroacetyl 200 equiv $\alpha$-chloroacetic acid anhydride was added.

For LC3A(ΔG120)–NHNH methyl fumarate 20 equiv methyl fumarate anhydride was added.

##### 3.2. Generation of Ub hydrazides from Ub-*Mxe* GyrA-$His_6$ fusions

Ub-*Mxe* GyrA-$His_6$ fusion after Ni-NTA purification was dialyzed against intein cleavage buffer (25 mM HEPES, 200 mM NaCl, 1 mM EDTA pH 7.2 at 4 °C). After dialysis hydrazine monohydrate (100 mM) was added and the pH adjusted to 7.8. The solution was allowed to stand at room temperature and monitored using LC-MS analysis. Upon completion the solution was dialyzed against 25 mM HEPES, 200 mM NaCl pH 7.2 at 4 °C. Cleaved intein was removed using gravity Ni-NTA purification. The flow-through was collected and combined with resin washes. The combined fractions were dialyzed and further purified using ion exchange.

##### 3.3. Generation of GABARAP and LC3A hydrazides from SUMO-GABARAP/LC3A-*Mxe* GyrA-$His_6$ fusions

SUMO-GABARAP/LC3A-*Mxe* GyrA-$His_6$ fusion after Ni-NTA purification was dialyzed against intein cleavage buffer (25 mM HEPES, 500 mM NaCl, 1 mM EDTA pH 7.2 at 4 °C). After dialysis hydrazine monohydrate (75 mM) and MESNa (10 mM) was added and the pH adjusted to 7.8. The solution was allowed to stand at room temperature and monitored using LC-MS analysis. Upon completion the solution was dialyzed against 25 mM HEPES, 500 mM NaCl pH 7.2 at 4 °C. SENP1 (1 wt%) was added to cleave the SUMO-tag. The cleaved intein was removed using gravity Ni-NTA purification. The flow-through was collected and combined with resin washes. The combined fractions were dialyzed and further purified using cation exchange using a buffer 25 mM Tris pH 8.0 and a gradient of 0–500 M NaCl in the same buffer over 25 column volumes. The product fractions were pooled, concentrated, and stored at –80 °C until further use.

##### 3.4. Synthesis of symmetrical anhydrides

The carboxylic acid was dissolved in $CH_2Cl_2$ (300 mM) and dicyclohexylcarbodiimide (0.5 eqiv) was added in one portion. The reaction was stirred under $N_2$ for 18 h. The suspension was filtered and concentrated to ~10% of the initial volume. The solution was placed at 4 °C for 1 h and filtered again. The filtrate was concentrated to obtain the symmetrical anhydride which was used without further purification. Anhydrides can be stored at –20 °C for weeks.

##### 3.5. Reaction of modified Ubl probes with E2s

Ubl probes (15 µM) were mixed with E2s (15 µM) in PBS. The mixture was incubated at 37 °C in a water bath for 0.5-3 h. The reaction was quenched by addition of 2x Laemmli buffer and boiled at 95 °C for 5 min. Samples were resolved by SDS-PAGE (8-16%, Bio-Rad) and visualized by Coomassie staining. If required, gel bands were quantified by gel densitometry using ImageLab (Bio-Rad).

##### 3.6. Competition experiment in the presence of $LIR^{ATG3}$ peptides

ATG3 C1 (15 µM) was mixed with peptides **5** or **6** (1 µM – 1 mM) in PBS. GABARAP methyl fumarate **2** (15 µM) was added and the reaction mixture was incubated at 37 °C for 30 min. The reaction was quenched by addition of 2x Laemmli buffer and boiled at 95 °C for 5 min. Samples were resolved by

SDS-PAGE (8-16%, Bio-Rad) and visualized by Coomassie staining. Gel bands were quantified by gel densitometry using ImageLab (Bio-Rad).

##### 3.7. Alanine-screen

ATG3 C1 (15 µM) was mixed with GABARAP methyl fumarate **2** (15 µM) in PBS, and the reaction mixture was incubated at 37 °C for 30 min. The reaction was quenched by addition of 2x Laemmli buffer and boiled at 95 °C for 5 min. Samples were resolved by SDS-PAGE (8-16%, Bio-Rad) and visualized by Coomassie staining. Gel bands were quantified by gel densitometry using ImageLab (Bio-Rad).

##### 3.8. NaCl dependence

ATG3 C1 (15 µM) was mixed with phosphate buffer with varying concentrations of NaCl (0-500 mM). GABARAP methyl fumarate **2** (15 µM) was added and the reaction mixture was incubated at 37 °C for 30 min. The reaction was quenched by addition of 2x Laemmli buffer and boiled at 95 °C for 5 min. Samples were resolved by SDS-PAGE (8-16%, Bio-Rad) and visualized by Coomassie staining. Gel bands were quantified by gel densitometry using ImageLab (Bio-Rad).

##### 3.9. Structure prediction

The ATG3-GABARAP complex was predicted using the publicly available ColabFold work-book (https://colab.research.google.com/github/sokrypton/ColabFold/blob/main/AlphaFold2.ipynb). ATG3 sequence (Q9NT62) and mature-GABARAP (O95166) were used as input sequences. Structure prediction was performed in no-template mode with msa_mode “MMseqs2 (UniRef+Environmental)” and pair_mode “unpaired+paired”.

##### 3.10. Synthesis of linear peptides

Amino acids were loaded according to the general methods. Automated peptide elongation was carried out on a Multisyntech Syro I parallel synthesizer according to the general peptide methods. After coupling of the final amino acid Fmoc group was removed by treatment with 20 vol% piperidine. The resin was thoroughly washed. Acetic anhydride (2 equiv) and DIPEA (4 equiv) in DMF were added to the resin and agitated for 2h. The peptide was cleaved form the resin using TFA/DODT/$H_2O$ (95:2.5:2.5, v/v) for 1 h. The resin was removed by filtration and the filtrate concentrated under reduced pressure. The solution was triturated with $Et_2O$ and centrifuged to obtain crude peptide. The crude peptide was dissolved in $H_2O/CH_3N$ (1:1, v/v) + 0.1% (v/v) TFA and purified using preparative HPLC.

##### 3.11. Synthesis of cyclic peptides

Amino acids were loaded according to the General Methods. Automated peptide elongation was carried out on a Multisyntech Syro I parallel synthesizer according to the general peptide methods. After coupling of the final amino acid Fmoc group was removed by treatment with 20 vol% piperidine. The resin was thoroughly washed. $\alpha$-Chloroacetic anhydride (2 equiv) and DIPEA (4 equiv) in DMF were added to the resin and agitated for 2h. The resin was thoroughly washed. The peptide was cleaved form the resin using TFA/DODT/$H_2O$ (95:2.5:2.5, v/v) for 1 h. The resin was removed by filtration and the filtrate concentrated under reduced pressure. The solution was triturated with $Et_2O$ and centrifuged to obtain crude peptide. The peptide was dissolved in 6 M Gdn HCl, 200 mM $NaCO_3$ (pH 9.5, 20 mL, 5 mM with respect to initial resin loading) and shaken at rt for 1 h. The reaction mixture was diluted with $H_2O/CH_3N$ (1:1, v/v) + 0.1% (v/v) TFA and purified using preparative HPLC.

##### 3.12. Synthesis of fluorescently labelled peptides

Fmoc-Lys(alloc)-OH was loaded according to the General Methods. Automated peptide elongation was carried out on a Multisyntech Syro I parallel synthesizer according to the general peptide methods. The side-chain alloc group was removed by treatment with $Pd(PPh_3)_4$ (0.25 equiv) and $PhSiH_3$ (24 equiv) in $CH_2Cl_2$. The deprotection step was repeated once. The resin was thoroughly washed. FITC (2 equiv) and DIPEA (4 equiv) were added in DMF and shaken for 2 h at rt. All steps following FITC coupling were performed in the dark. The resin was removed by filtration and the filtrate concentrated under reduced pressure. The solution was triturated with $Et_2O$ and centrifuged to obtain crude peptide. The crude peptide was dissolved with $H_2O/CH_3N$ (1:1, v/v) + 0.1% (v/v) TFA and purified using preparative HPLC.

##### 3.13. Fluorescence polarization measurement

Fluorescent peptides were dissolved in DMSO and PBS. The concentration was determined by UV-absorption and extinction coefficient at 280 nm. The peptides were diluted to a 2X concentration of 20 nM using PBS. Protein concentration was determined using UV-absorption and extinction coefficient at 280 nm. Proteins were diluted to to twice their final concentration (40, 20, 10, 4, 3, 2, 1.5, 1, 0.75, 0.5, 0.375, 0.25 µM) . Peptides and proteins were mixed 1:1 to a final peptide concentration of 10 nM and protein concentrations of 20, 10, 5, 2, 1.5, 1, 0.75, 0.5, 0.375, 0.25, 0.1875, 0.125 µM. The reactions were incubated at room temperature for 45 min. Fluorescence polarization and intensity was measured on a Victor Nivo Plate Reader (Perkin Elmer). Fluorescence polarization was plotted against concentration and the binding curve was fitted in PRISM (GraphPad) using least square regression to extract binding affinities. The measurements were performed in triplicates and plotted as such. Errors were estimated based on these triplicates.

##### 3.14. Co-crystallization of GABARAP with LIR$^{ATG3}$ peptide

GABARAP–OH in 25 mM Tris, 150 mM NaCl, pH 8.0 was concentrated using centrifugal filters (Amicon) to 730 µM ATG3 LIR peptide (Y90-H112) was resuspended in DMSO (50 mM) and 3 equivalents with respect to GABARAP were added. The reaction was dialyzed against 25 mM Tris, 150 mM NaCl, pH 8.0 at room temperature for 20 h. Crystallization was performed using hanging-drop vapour diffusion in 24-well crystallization flasks (Hampton Research). Protein solution (2 µL) was mixed with reservoir solution (2 µL) on siliconized cover-slips (Hampton Research). Crystallizations were performed at room temperature. Crystals were observed after 2 days. Crystals were obtained for 0.2 M NaOAc, 0.1 M sodium cacodylate, 30 vol% PEG 8,000, pH 6.5. Crystals were cryo-protected with 25 vol% glycerol and flash-frozen. Crystal diffraction was measured at the Swiss Light Source (Paul Scherrer Institute, Villigen, Switzerland) beamline X06DA (PXIII) equipped with the PILATUS 2M-F detector system (Dectris, Switzerland) at a wavelength of 1.0 Å, while the crystal was kept at 100 K. XDS software was used for data processing[2]. Molecular replacement using the GABARAP structure from PDB structure 6HB9 as the search model was performed using Phaser[3,4]. The asymmetric unit contained eight copies of the GABARAP–LIR$^{ATG3}$ peptide complex. Refinement was performed iteratively using phenix.refine from the Phenix[5] software package and Coot[6]. Protein structure visualization and analysis (RMSD, Coulombic potential, surface area) was performed using the UCSF ChimeraX software package[7].

##### 3.15. Generation of HCT116 ATG3 knock-down cell-line using CRISPRi

HCT116 cells expressing dCas9-KRAB were prepared according to published procedures.[8] sgRNA were incubated with T4 polynucleotide kinase (NEB) at 37 °C for 30 min and then annealed by incubation at 95 °C for 5 min and cooling to 25 °C at 5 °C/min. Annealed sgRNAs were cloned into pCRISPRia–v2 (Addgene #84832) using BstXI and BlpI restriction enzyme sites. Correct insertion was validated by Sanger sequencing.

Lentiviral packaging was performed in HEK293T cells using Xtreme-Gene HP (Roche) transfection reagent according the manufacturer's procedure. pVSVG (Addgene #8454), pR8.2 dvpr (Addgene #8455) and pCRISPRia–v2 carrying the sgRNA of interest were transfected using a ration of 4:1:3.5 (w/w/w). The media was changed after 24 h and lentiviral supernantant harvested after an additional 24 h. HCT116 dCas9-KRAB cells were transfected with lentiviral supernatant in the presence of 10 µg/mL polybrene. After 24 h cell media was changed. After additional 24 h, selection was initiated by supplementing cell media with 2.5 µM puromycin. Selection was performed for additional 3 d until all non-transfected control cells had died. ATG3 knock-down was confirmed using Anti-ATG3 Western blotting. sgRNA 1 showed most efficient knock-down and was used for all experiments. Used primer pairs are listed in Supplementary Table S5

##### 3.16. Generation of HEK293T ATG3 -/- using CRISPR-Cas9

ATG3-deficient HEK293T cell lines were derived by nucleofection of recombinant Cas9 and in vitro transcribed (IVT) sgRNAs as described previously[9]. In brief, IVT templates were generated by extension PCR using Phusion Polymerase (New England Biolabs) with oligonucleutides T7FwdAmp (2 µM), T7 RevAmp (2 µM), T7RevLong (0.02 µM) and one of two oligonucleotides bearing an ATG3-targeting protospacer (T7FwdVar_sgATG3.KO3 and T7FwdVar_sgATG3.KO4, each 0.02 µM). Protospacers were chosen based on previously optimized design rules[10]. Crude PCR products were subjected to IVT (HiScribe T7 High yield RNA Synthesis kit, New England Biolabs, 37°C overnight), dephosphorylated (Quick CIP, New England Biolabs, 5U per reaction, 37°C, 30 min) and column purified (RNeasy Mini kit, QIAGEN).

120 pmol IVT product were complexed with 100 pmol recombinant His$_6$-Cas9-NLS (kindly provided by the Genome Engineering and Measurement Lab, GEML, Functional Genomics Center Zürich) for 20 min at room temperature in RNP assembly buffer (150 mM KCl, 20 mM HEPES, 1mM $MgCl_2$, pH 7.5) followed by delivery using a 4D-nucleofector device (Lonza). Sub-clonal cell lines were isolated by cell sorting (SONY SH-800). Genomic DNA of resulting clonal cell lines was isolated (QuickExtract, Lucigen) followed by PCR of target regions with Q5 Polymerase (New England Biolabs) using one of two primer pairs (sgATG3.KO3_seq or sgATG3.KO4_seq). Homozygous knockout lines were identified by sanger sequencing and deconvolution of composite chromatograms (ICE Analysis Tool, Synthego). All used primer pairs are listed in Supplementary Table S6.

##### 3.17. *In cellulo lipidation* of LC3A/B

HEK293T ATG3 -/- cells were grown in a 6-well plate. pCMV-3xFLAG-ATG3 plasmids containing ATG3 wt, C264A or ΔLIR were transfected using Xtreme-Gene HP (Roche) in OptiMem. After 24 h media was changed to DMEM. After an additional 24 h the media was changed to HBSS supplemented with 40 µM chloroquine and incubated for 60 min. Cells were directly lysed using RIPA-buffer (50 mM Tris, 150 mM NaCl, 1 vol% Nonidet P-40, 0.5 wt% deoxycholate, 0.1 wt% SDS, pH 7.5) in the presence of protease inhibitors (Thermo-Fisher). Insoluble cell components were removed by centrifugation. Protein concentrations were measured by BCA according to the manufacturers protocol. Cell lysates were diluted to equal protein concentrations using RIPA-buffer. 10 µg of total protein was loaded and used for SDS-PAGE and Western bot analysis.

##### 3.18. Co-immunoprecipitation of ATG3

HCT116 ATG3 knock-down cells, generated by CRISPR inhibition, were grown in T25 flasks. pCMV-3xFLAG-ATG3 plasmids containing ATG3 wt, C264A or ΔLIR were transfected using Xtreme-Gene

HP (Roche) in OptiMem. After 24 h media was changed to DMEM. After an additional 24 h the media was changed to HBSS supplemented with 40 µM chloroquine and incubated for 60 min. Cells were directly lysed using NP-40 buffer (50 mM Tris, 150 mM NaCl, 1 vol% nonidet-P40, pH 8.0) in the presence of protease inhibitors (Thermo-Fisher). Insoluble cell components were removed by centrifugation. An aliquot of lysate was removed for further analysis. The remaining cell lysate was incubated with Anti-FLAG M2 magnetic beads (20 µL, Sigma-Aldrich) for 2 h at 4 °C. The resin was washed with NP40 lysis buffer and PBS. Proteins were eluted by addition of 4x non-reducing Laemmli buffer and boiling at 95 °C for 5 min. β-mercaptoethanol was added to the eluted proteins. Input and elution were analyzed by SDS-PAGE and Western blot analysis.

##### 3.19. Analysis of *in cellulo* ATG3 thioester formation

HEK293T ATG3 -/- cells were grown in a 6-well plate. pCMV-3xFLAG-ATG3 plasmids containing ATG3 wt, C264A or ΔLIR were transfected using Xtreme-Gene HP (Roche) in OptiMem. After 24 h media was changed to DMEM. After an additional 24 h the media was changed to HBSS supplemented with 40 µM chloroquine and incubated for 60 min. Cells were directly lysed using NP-40 buffer (50 mM Tris, 150 mM NaCl, 1 vol% nonidet-P40, pH 8.0) in the presence of protease inhibitors (Thermo-Fisher). Insoluble cell components were removed by centrifugation. Aliquots of cell lysate were removed and directly quenched by the addition of either 4x reducing Laemmli buffer or 4x non-reducing Laemmli buffer. Samples containing reducing Laemmli buffer were further boiled for 5 min at 95 °C. Samples were analyzed by SDS-PAGE and Western blot analysis.

##### 3.20. ATG3 pull-down using 3xFLAG-ATG7

HCT116 cells were grown in T25 flasks. pCMV-3xFLAG-ATG7 plasmid was transfected using Xtreme-Gene HP (Roche) in OptiMem. After 24 h media was changed to DMEM and incubated for additional 24 h. Cells were directly lysed using NP-40 buffer (50 mM Tris, 150 mM NaCl, 1 vol% nonidet-P40, pH 8.0) in the presence of protease inhibitors (Thermo-Fisher). Insoluble cell components were removed by centrifugation. Cell lysate was incubated with Anti-FLAG M2 magnetic beads (40 µL, Sigma-Aldrich) for 2 h at 4 °C. The resin was washed with NP40 lysis buffer and PBS. Recombinant ATG3 variants (wt, ΔLIR, W107A, 15 µM) were added to immobilized ATG3 resin and incubated for 1 h at 4 °C. The resin was washed with NP40 lysis buffer and PBS. Proteins were eluted by addition of 4x non-reducing Laemmli buffer and boiling at 95 °C for 5 min. β-mercaptoethanol was added to the eluted proteins. Input and elution were analyzed by Sypro-Ruby staining and SDS-PAGE analysis.

##### 3.21. Preparation of Fluorescein-GABARAP

Cysteine-GABARAP was expressed as a His-SUMO linear fusion as described above. Following SUMO-tag cleavage, GABARAP was concentrated to 100 µM and mixed with fluorescein-iodoacetamide in DMF (3 mg) and incubated at room temperature. Reaction progress was monitored using LC-MS analysis. After 3 h excess small molecule was removed by buffer exchange to PBS using MidiPrep desalting columns (Cytiva). Buffer exchange was repeated (2x) to provide fluorescein-GABARAP.

##### 3.22. Pulse-chase assay of GABARAP transfer to ATG3

Fluorescein-GABARAP (10 µM) was mixed with ATG7 (1 µM) in 25 mM Tris, 150 mM NaCl, pH 7.5. ATP-$MgCl_2$ (5 mM) was added and the mixture was incubated at 4 °C for 15 min. The solution was diluted 3-fold with 25 mM Tris, 150 mM NaCl, 50 mM EDTA, pH 7.5 and buffer exchanged using a Zeba-desalting column to 25 mM Tris, 150 mM NaCl, 50 mM EDTA, pH 7.5. The solution was added to recombinant ATG3 (0.5 µM) variants and incubated for 5 min. Aliquots were removed after 1, 2.5 and 5 min and mixed with either 4x non-reducing Laemmli buffer or 4x reducing Laemmli buffer. Samples were resolved using SDS-PAGE and analyzed by in-gel fluorescence and Stain-free visualization. ATG3~GABARAP band was quantified by gel densitometry (Image-Lab, BioRad) and normalized to ATG3~GABARAP band for wild-type ATG3 at 5 min.

#### 4. Characterization of peptides and proteins

##### 4.1. Linear LIR$^{ATG3}$ peptide 5

**Leu-Glu-Ala-Ile-Ile-Glu-Glu-Asp-Asp-Gly-Asp-Gly-Gly-Trp-Val-Asp-Thr-Tyr-His-Ala-Gly-Gly-Gly-Lys**

HRMS (ESI): calculated for $[C_{110}H_{162}N_{28}O_{42}]^{2+}$: m/z 1273.5695, found: m/z 1273.5712

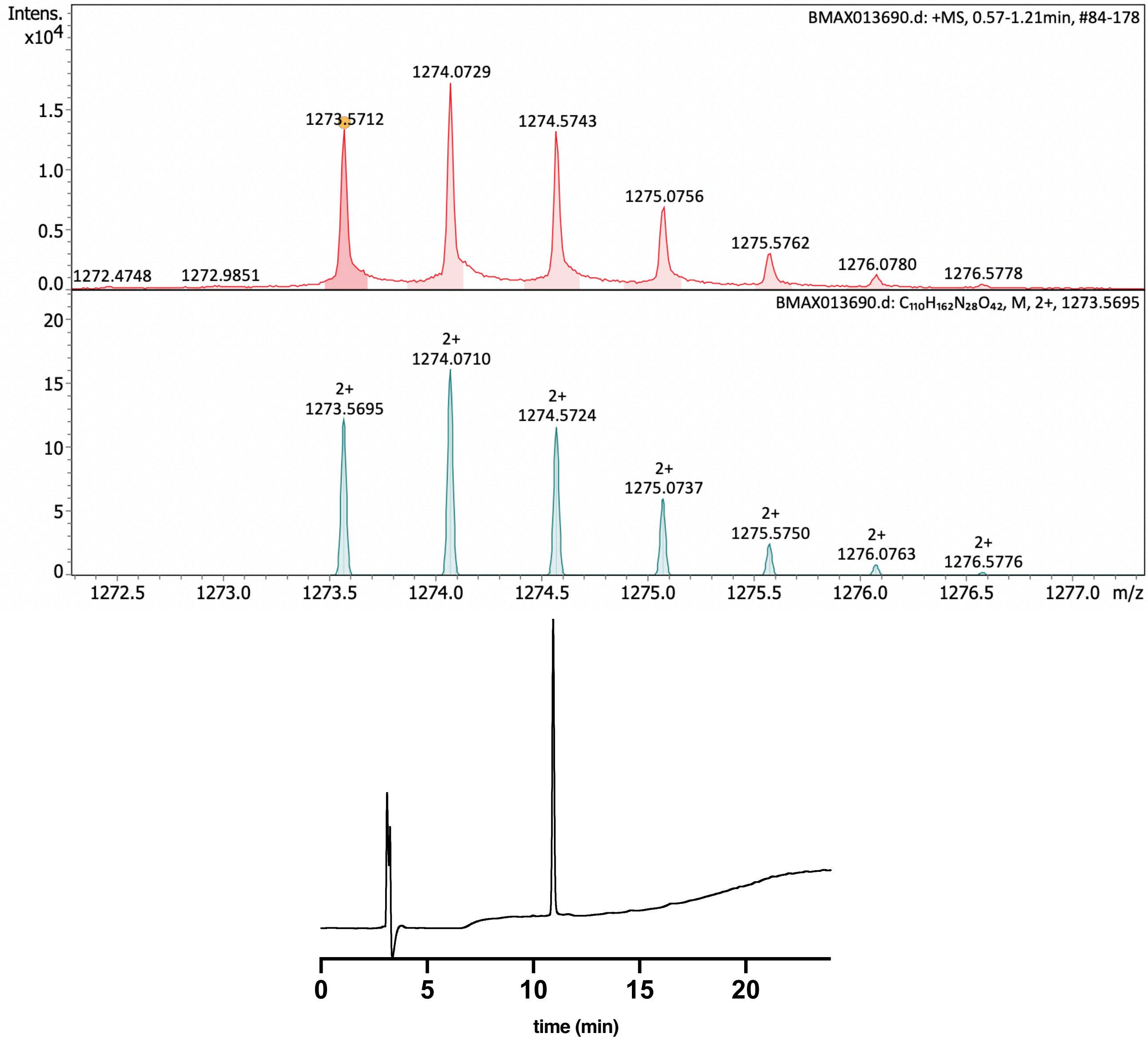

**Supplementary Figure 5 Characterization of peptide 5** HRMS (ESI) spectrum of purified peptide showing recorded mass spectrum (upper panel) and calculated spectrum (lower panel). Analytical RP-HPLC trace of **5**. Absorption at 220 nm is shown.

##### 4.2. Cyclic LIR$^{ATG3}$ peptide 6

**Leu-Glu-Ala-Ile-Ile-Glu-Glu-Asp-Asp-Gly-Asp-Gly-Gly-Trp-Val-Asp-Thr-Tyr-His-Cys**

HRMS (ESI): calculated for $[C_{95}H_{134}N_{22}O_{37}S]^{2+}$: m/z 1103.4495, found: m/z 1103.4492

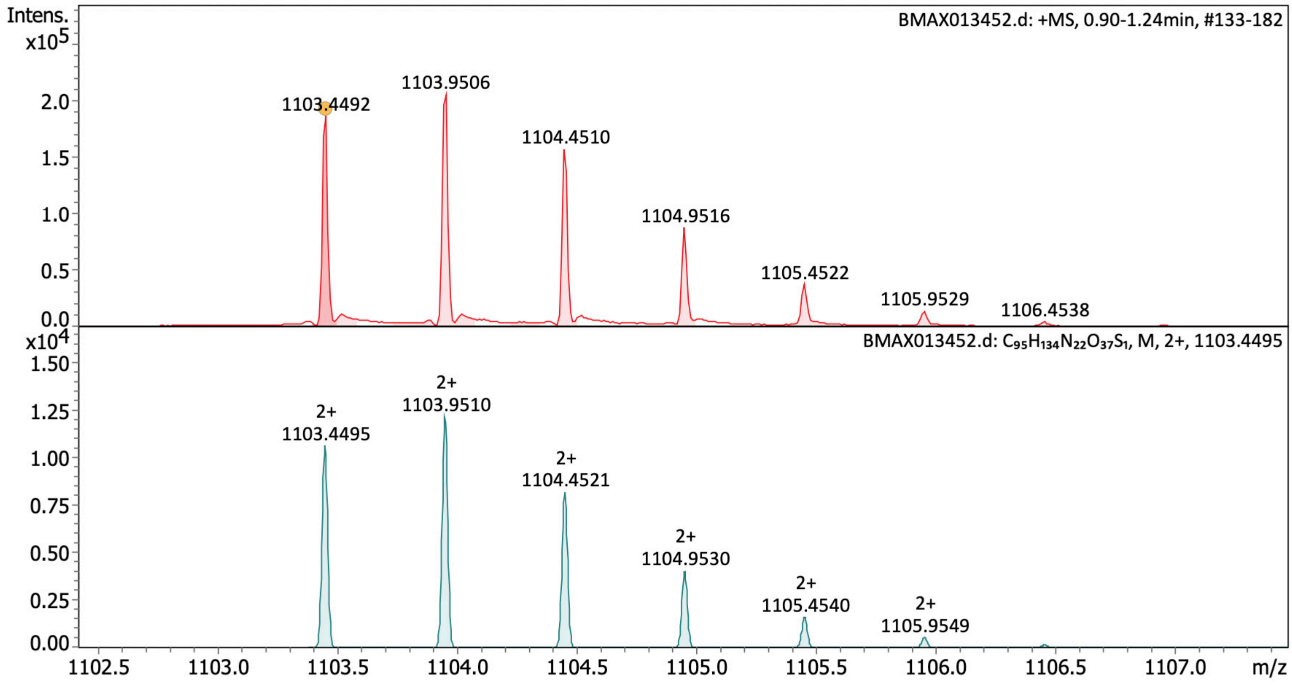

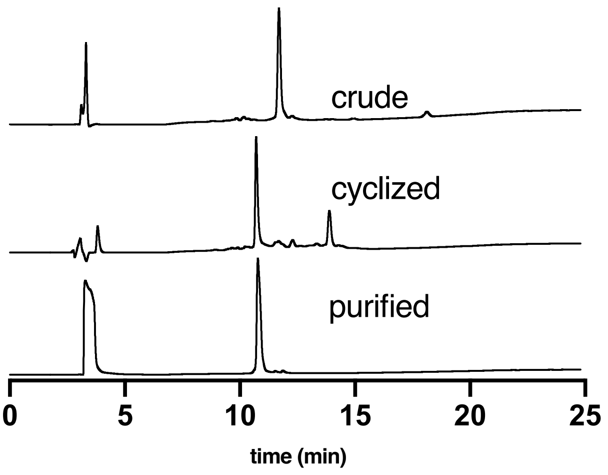

**Supplementary Figure 6 Characterization of peptide 6** HRMS (ESI) spectrum of purified peptide showing recorded mass spectrum (upper panel) and calculated spectrum (lower panel). RP-HPLC of linear peptide after resin cleavage, crude RP-HPLC trace during peptide cyclization, RP-HPL of purified cyclized peptide **6**. Absorption at 220 nm is shown.

##### 4.3. Fluorescent linear LIR$^{ATG3}$ peptide

**Leu-Glu-Ala-Ile-Ile-Glu-Glu-Asp-Asp-Gly-Asp-Gly-Gly-Trp-Val-Asp-Thr-Tyr-His-Ala-Gly-Gly-Gly-Lys**

The peptide was obtained as a white solid.
HRMS (ESI): calculated for $[C_{131}H_{173}N_{29}O_{47}S]^{2+}$: m/z 1468.0874, found: m/z 1468.0909

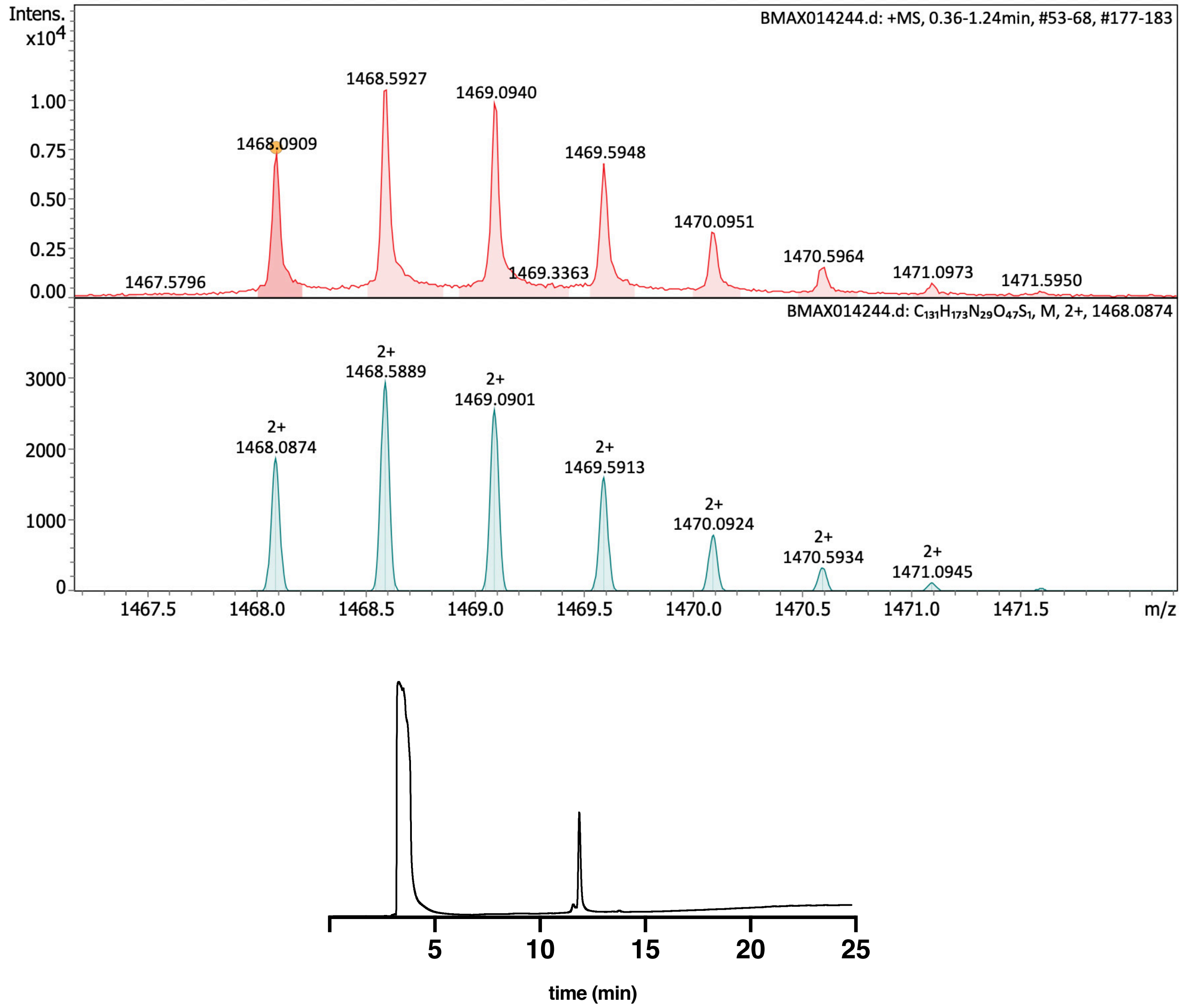

**Supplementary Figure 7 Characterization of fluorescent linear LIR$^{ATG3}$ peptide** HRMS (ESI) spectrum of purified peptide **5** showing recorded mass spectrum (upper panel) and calculated spectrum (lower panel). Analytical RP-HPLC trace of purified peptide. Absorption at 220 nm is shown.

##### 4.4. Fluorescent cyclic LIR$^{ATG3}$ peptide

**Leu-Glu-Ala-Ile-Ile-Glu-Glu-Asp-Asp-Gly-Asp-Gly-Gly-Trp-Val-Asp-Thr-Tyr-His-Cys-Gly-Gly-Gly-Lys**

HRMS (ESI): calculated for $[C_{131}H_{171}N_{29}O_{47}S2]^{2+}$: m/z 1483.0656, found: m/z 1483.0690

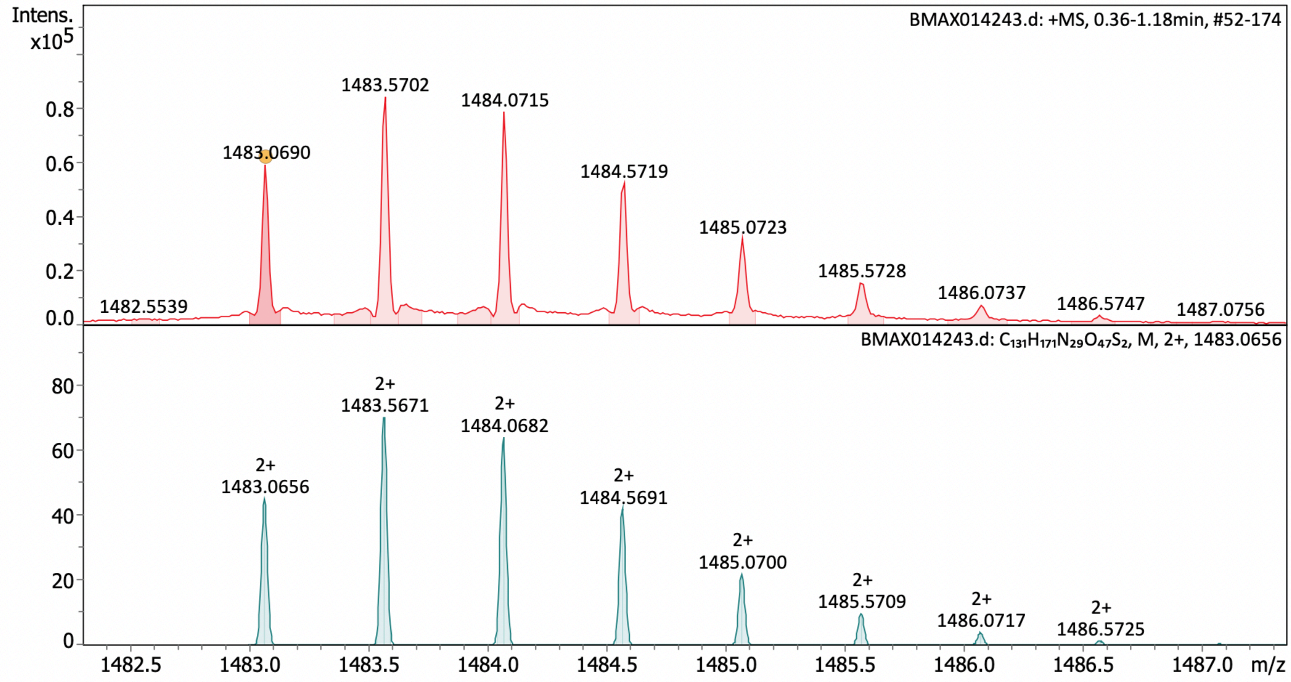

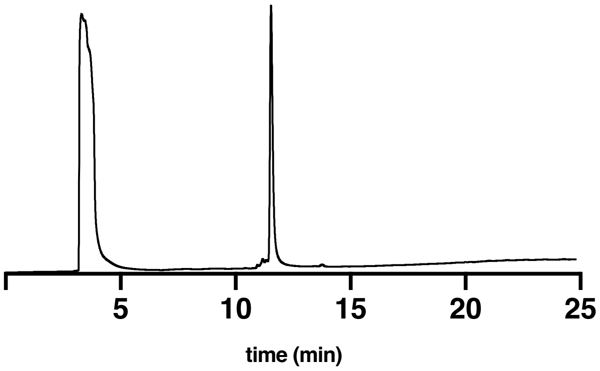

**Supplementary Figure 8 Characterization of fluorescent cyclic LIR$^{ATG3}$ peptide** HRMS (ESI) spectrum of purified peptide showing recorded mass spectrum (upper panel) and calculated spectrum (lower panel). Analytical RP-HPLC trace of purified peptide. Absorption at 220 nm is shown.

##### 4.5. LIR$^{ATG3}$ peptide for co-crystallization

**Tyr-Ser-Asp-Glu-Leu-Glu-Ala-Ile-Ile-Glu-Glu-Asp-Asp-Gly-Asp-Gly-Gly-Trp-Val-Asp-Thr-Tyr-His-Gly**

The peptide was obtained as a white solid (30 mg, 11%).
HRMS (ESI): calculated for $[C_{116}H_{163}N_{27}O_{47}]^{2+}$: m/z 1343.0592, found: m/z 1343.0592

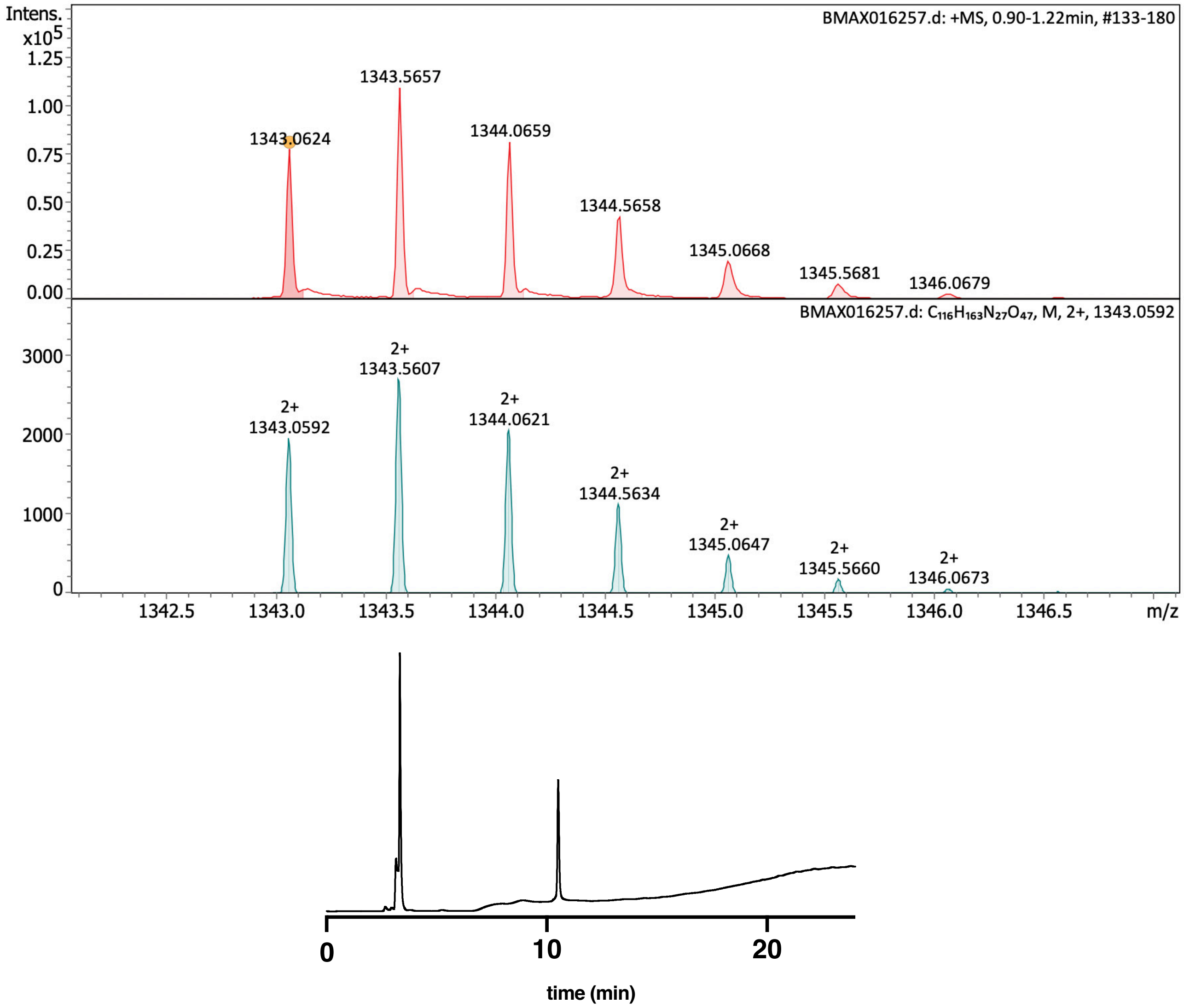

**Supplementary Figure 9 Characterization of LIR$^{ATG3}$ peptide** HRMS (ESI) spectrum of purified peptide showing recorded mass spectrum (upper panel) and calculated spectrum (lower panel). Analytical RP-HPLC trace of purified peptide. Absorption at 220 nm is shown.

#### 4.6. GABARAP

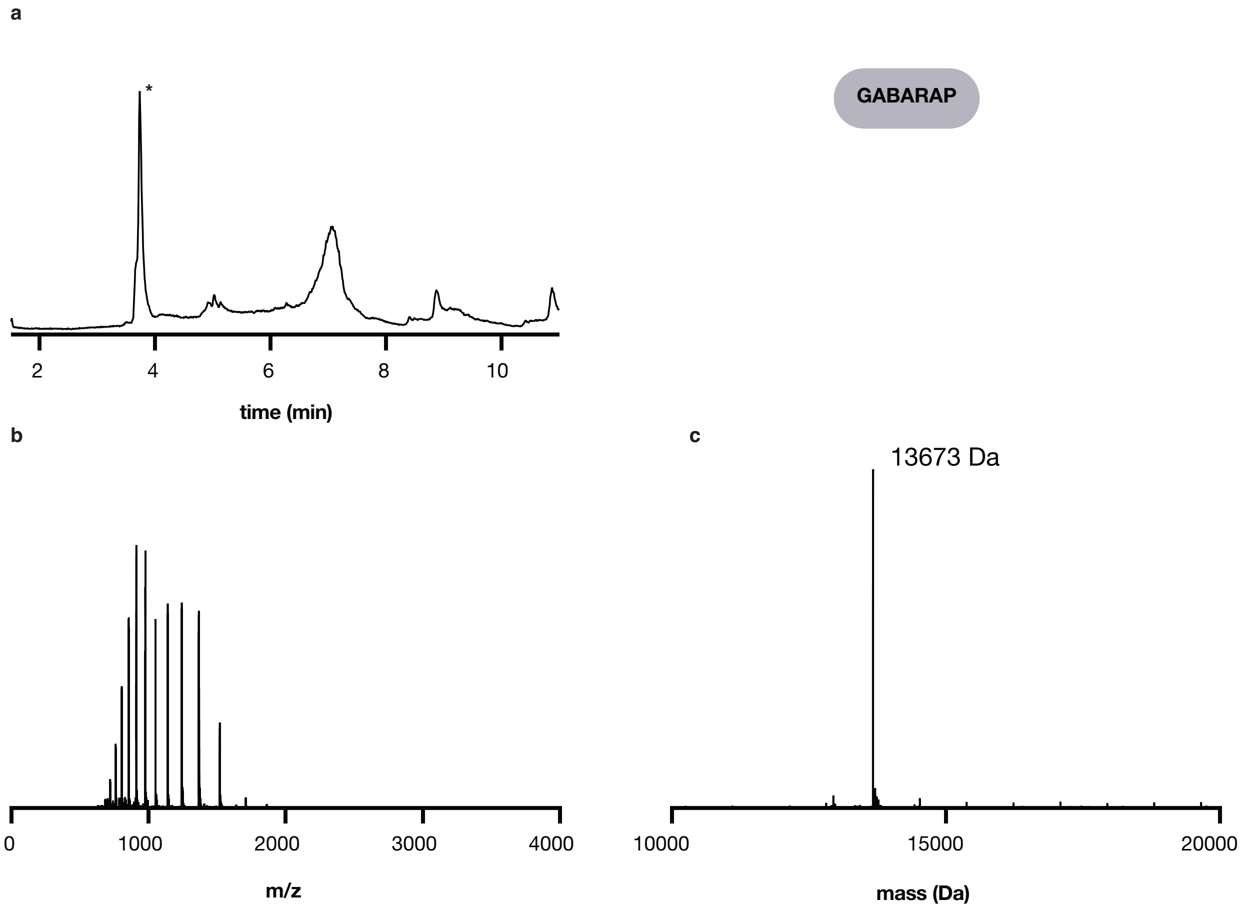

**Supporting Figure 10** Characterization of GABARAP. **a** LC-chromatogram of recombinantly expressed GABARAP. TIC is shown. Protein peak is marked with a star. **b** Mass-spectrum (ESI) of recombinantly expressed GABARAP. **c** Deconvoluted mass spectrum of GABARAP. Expected mass 13673 Da.

##### 4.7. GABARAP(ΔG116)–$NHNH_2$

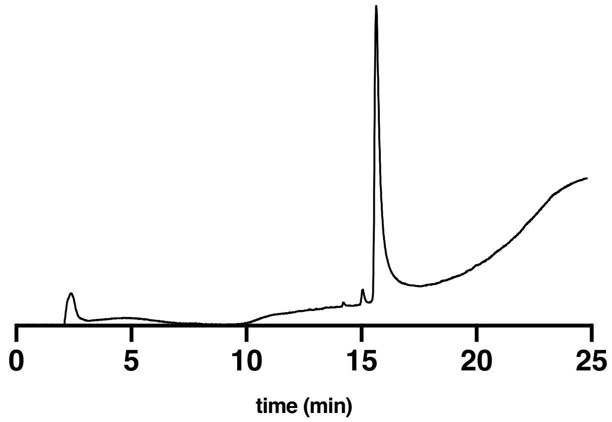

HRMS (ESI): calculated for $[C_{631}H_{965}N_{165}O_{171}S]^{+}$: m/z 13622.1608, found: m/z 13622.2880

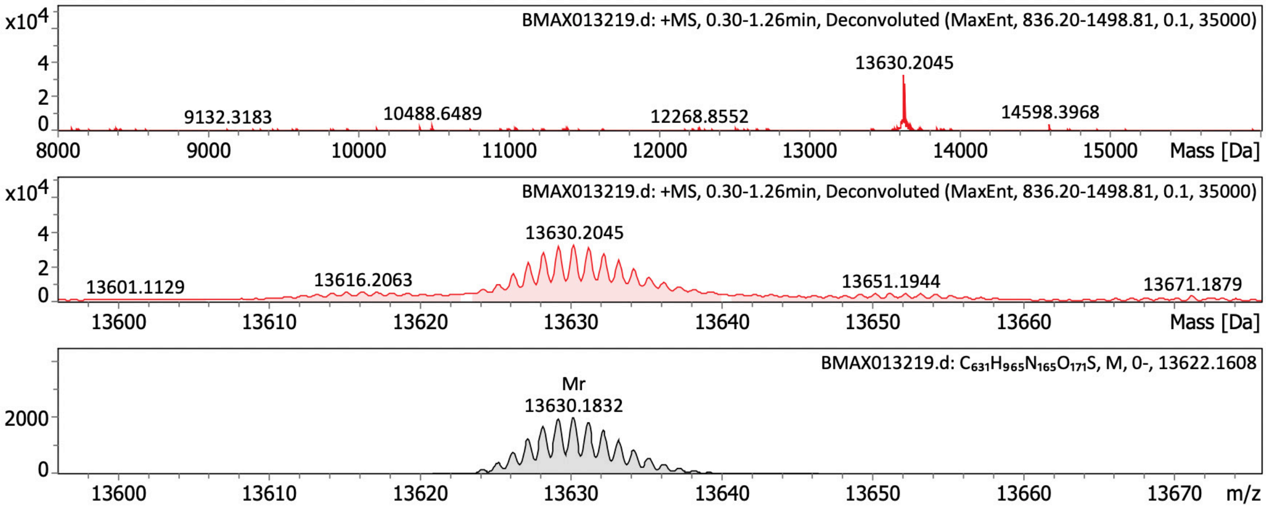

**Supporting Figure 11** Characterization of GABARAP(ΔG116)–$NHNH_2$ Analytical RP-HPLC of purified GABARAP ΔG116 $NHNH_2$. UV absorption at 220 nm is shown. HRMS (ESI) spectrum of GABARAP ΔG116 $NHNH_2$ showing recorded mass spectrum (upper panel), zoom in of recorded spectrum (middle panel) and calculated spectrum (lower panel).

##### 4.8. GABARAP(ΔG116)–NHNH α-chloroacetyl 1

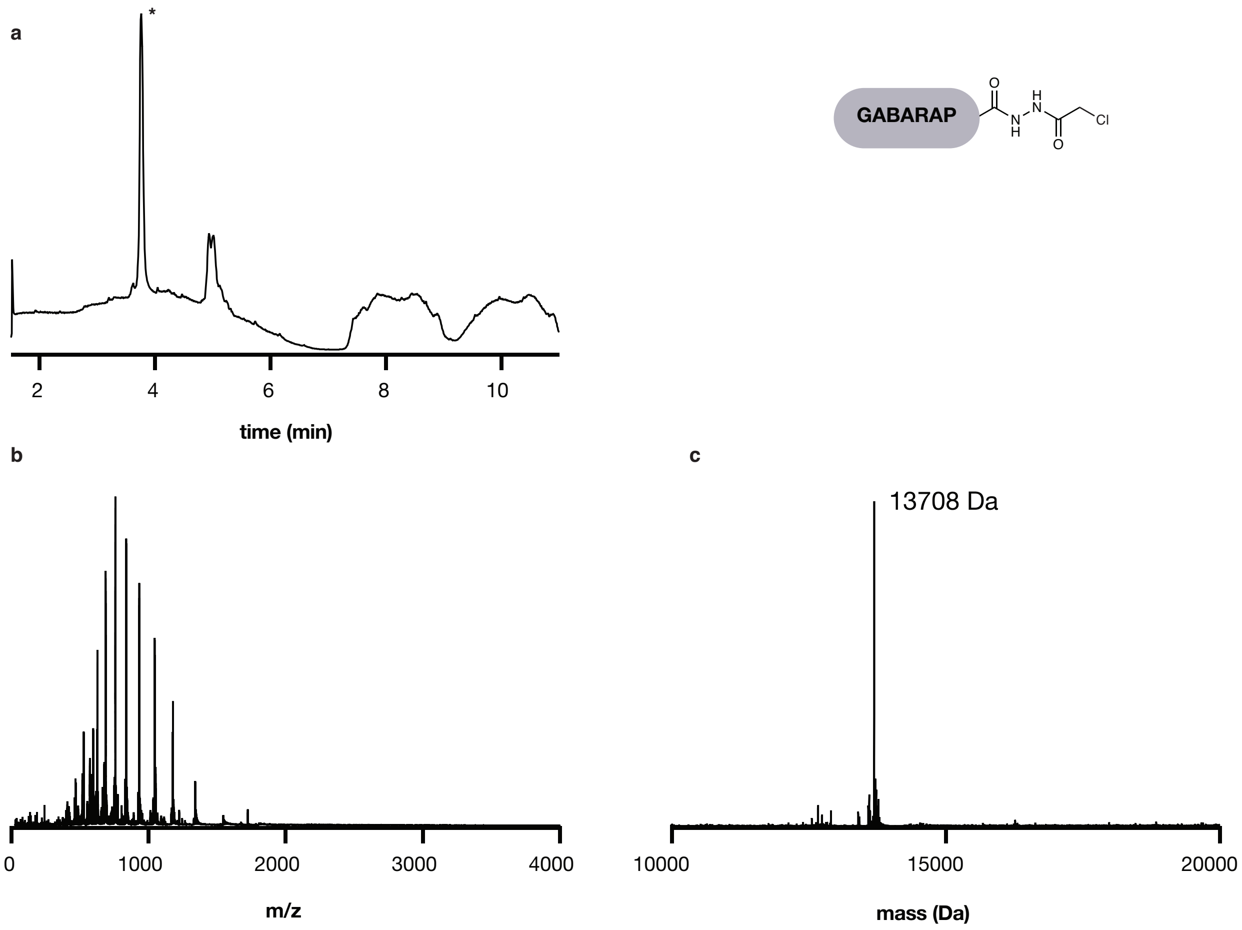

**Supplementary Figure 12** Characterization of GABARAP(ΔG116)–NHNH α-chloroacetyl **a** LC-chromatogram of GABARAP(ΔG116)–NHNH α-chloroacetyl. TIC is shown. Protein peak is marked with a star. **b** Mass-spectrum (ESI) of GABARAP(ΔG116)–NHNH α-chloroacetyl. **c** Deconvoluted mass spectrum of GABARAP(ΔG116)–NHNH α-chloroacetyl. Expected mass 13707 Da.

##### 4.9. GABARAP(ΔG116)–NHNH methyl fumarate 2

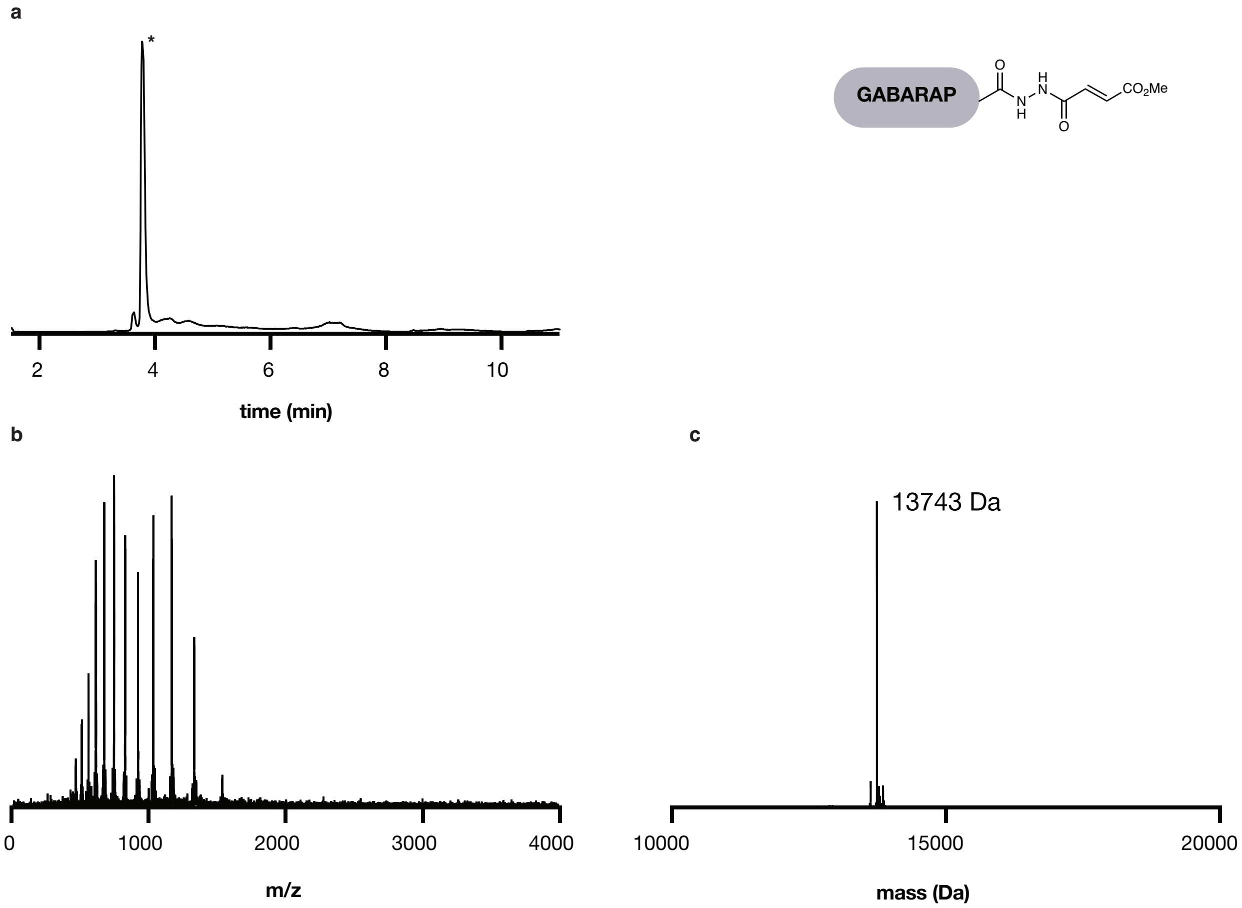

**Supplementary Figure 13** Characterization of GABARAP(ΔG116)–NHNH methyl fumarate. **a** LC-chromatogram of GABARAP(ΔG116)–NHNH methyl fumarate. TIC is shown. Protein peak is marked with a star. **b** Mass-spectrum (ESI) of GABARAP(ΔG116)–NHNH methyl fumarate. **c** Deconvoluted mass spectrum of GABARAP(ΔG116)–NHNH methyl fumarate. Expected mass 13742 Da.

**4.10. LC3A**

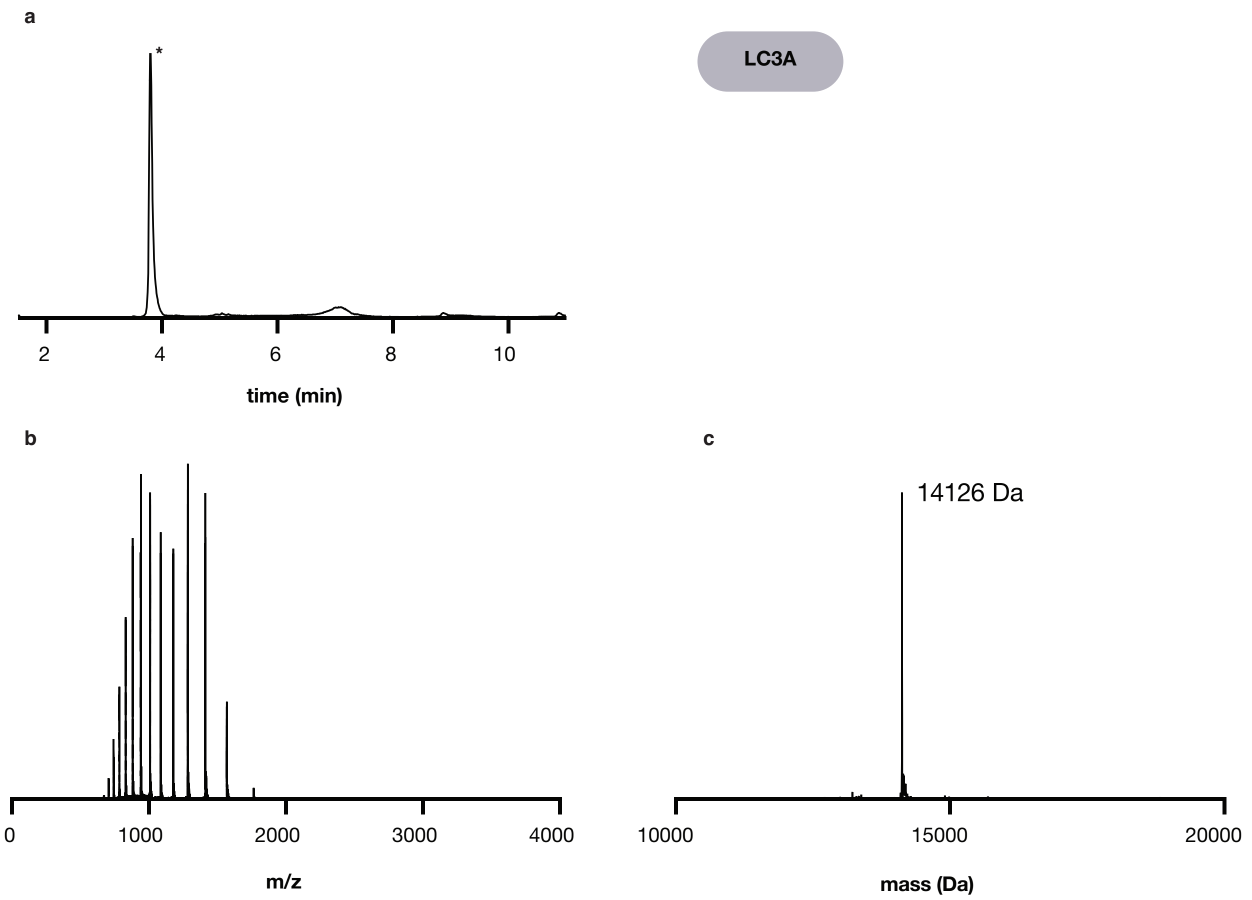

**Supplementary Figure 14** Characterization of LC3A. **a** LC chromatogram of recombinantly expressed LC3A. TIC is shown. Protein peak is marked with a star. **b** Mass-spectrum (ESI) of recombinantly expressed LC3A. **c** Deconvoluted mass spectrum of LC3A. Expected mass 14125 Da.

##### 4.11. LC3A(ΔG120)–$NHNH_2$

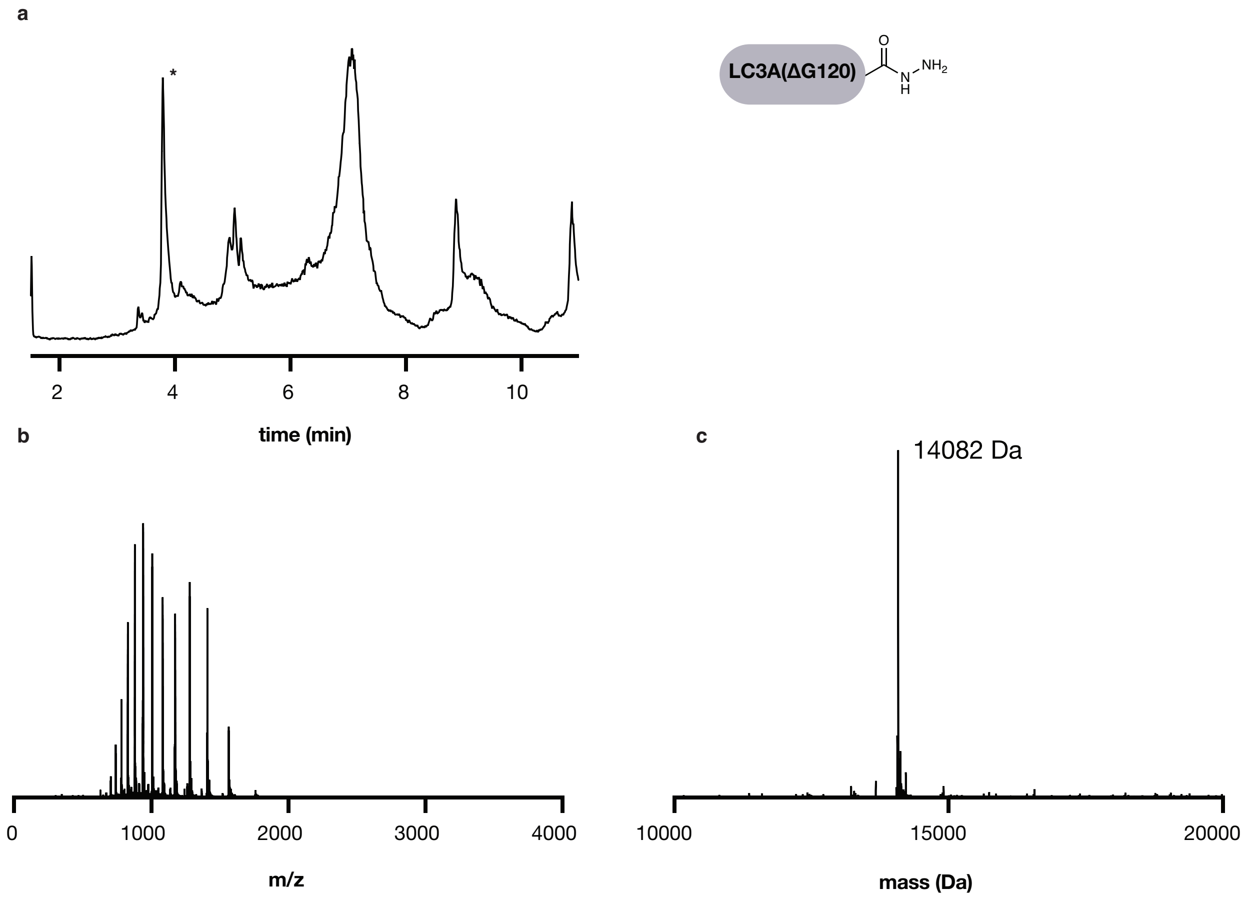

**Supplementary Figure 15** Characterization of LC3A(ΔG120)–$NHNH_2$ **a** LC-chromatogram of LC3A(ΔG120)–$NHNH_2$. TIC is shown. Protein peak is marked with a star. **b** Mass-spectrum (ESI) of LC3A(ΔG120)–$NHNH_2$. **c** Deconvoluted mass spectrum of LC3A(ΔG120)–$NHNH_2$. Expected mass 14082 Da.

##### 4.12. LC3A(ΔG120)–NHNH α-chloroacetyl 3

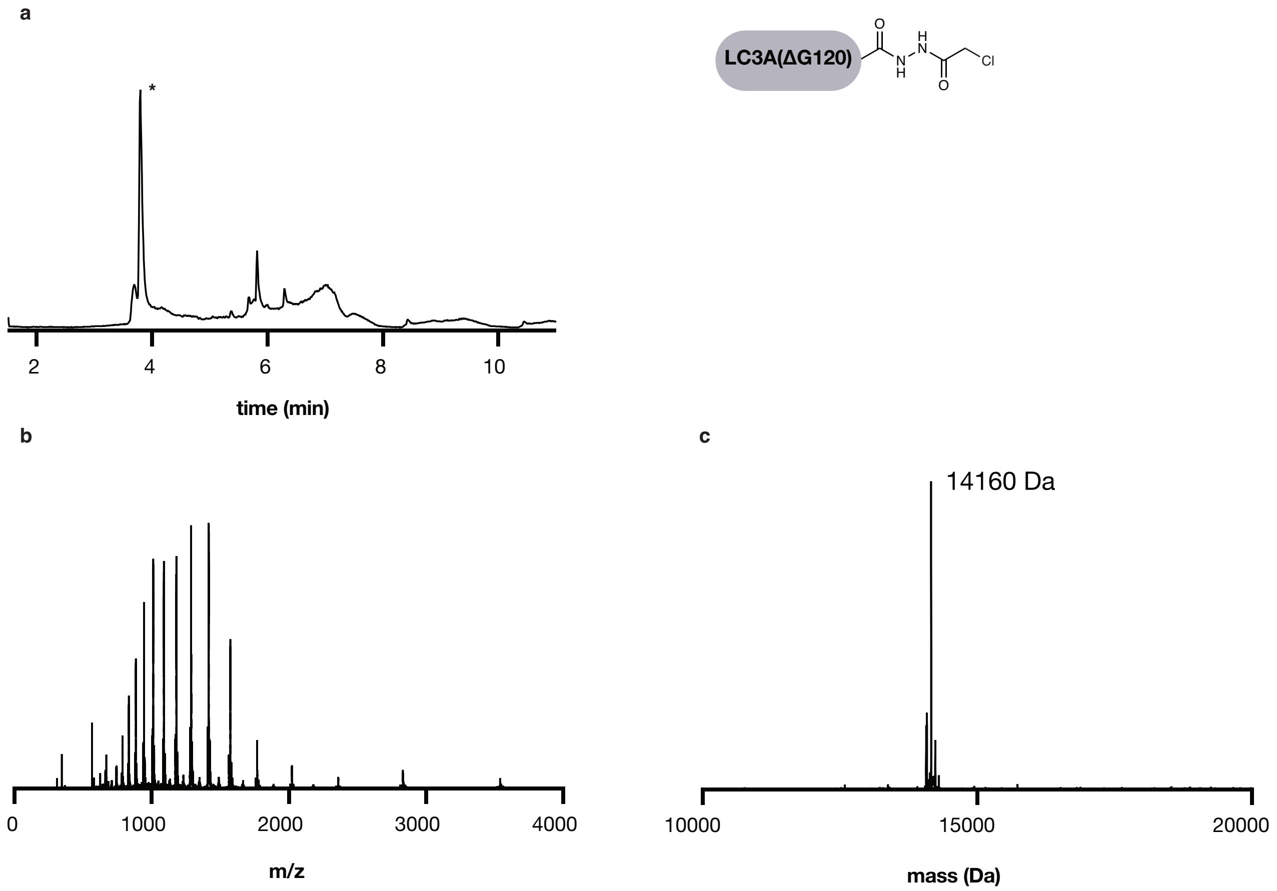

**Supplementary Figure 16** Characterization of LC3A(ΔG120)–NHNH α-chloroacetyl **a** LC-chromatogram of LC3A(ΔG120)–NHNH α-chloroacetyl. TIC is shown. Protein peak is marked with a star. **b** Mass-spectrum (ESI) of LC3A(ΔG120)–NHNH α-chloroacetyl. **c** Deconvoluted mass spectrum of LC3A(ΔG120)–NHNH α-chloroacetyl. Expected mass 14158 Da.

##### 4.13. LC3A(ΔG120)–NHNH methyl fumarate 4

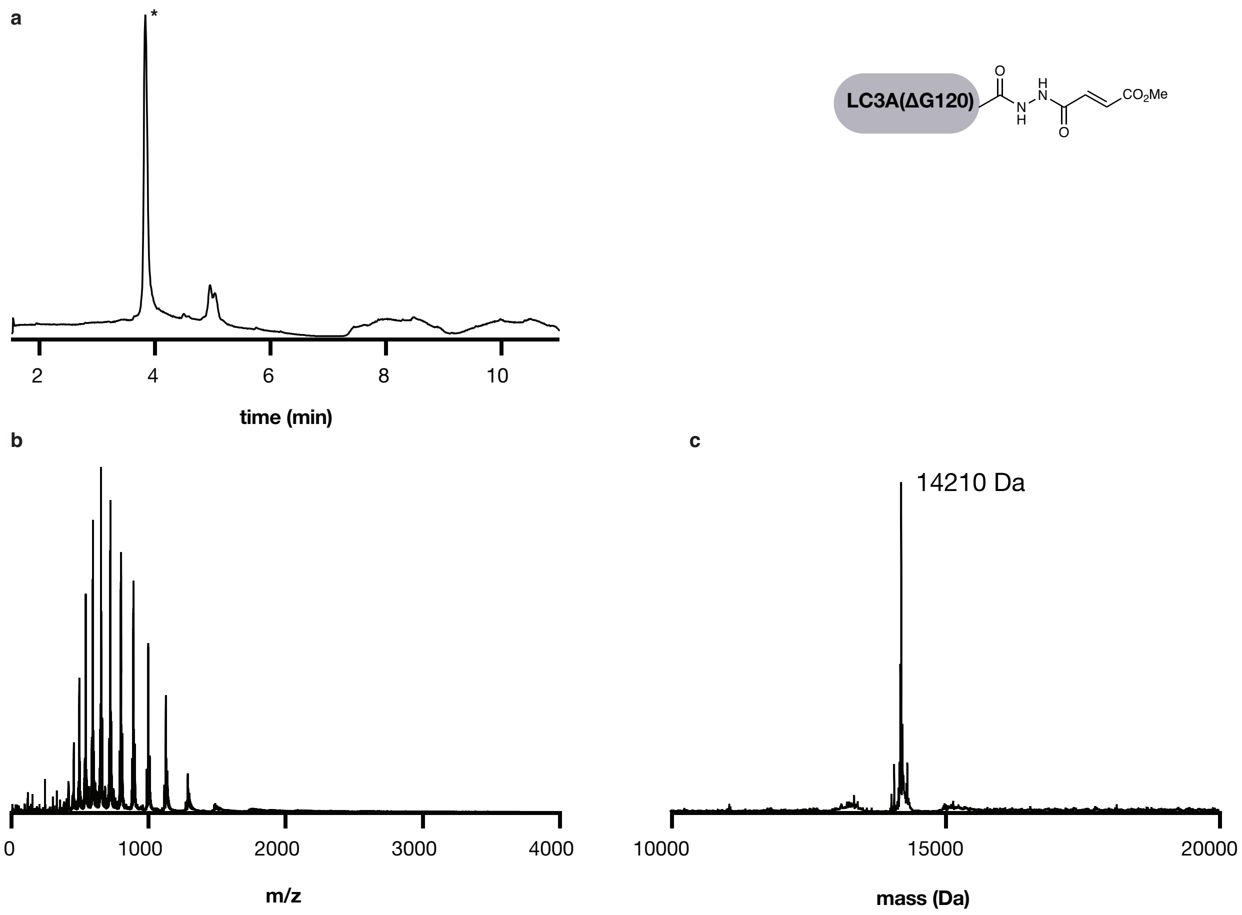

**Supplementary Figure 17** Characterization of LC3A(ΔG120)–NHNH methyl fumarate **a** LC-chromatogram of LC3A(ΔG120)–NHNH methyl fumarate. TIC is shown. Protein peak is marked with a star. **b** Mass-spectrum (ESI) of LC3A(ΔG120)–NHNH methyl fumarate. **c** Deconvoluted mass spectrum of LC3A(ΔG120)–NHNH methyl fumarate. Expected mass 14210 Da.

##### 4.14. GABARAP fluorescein

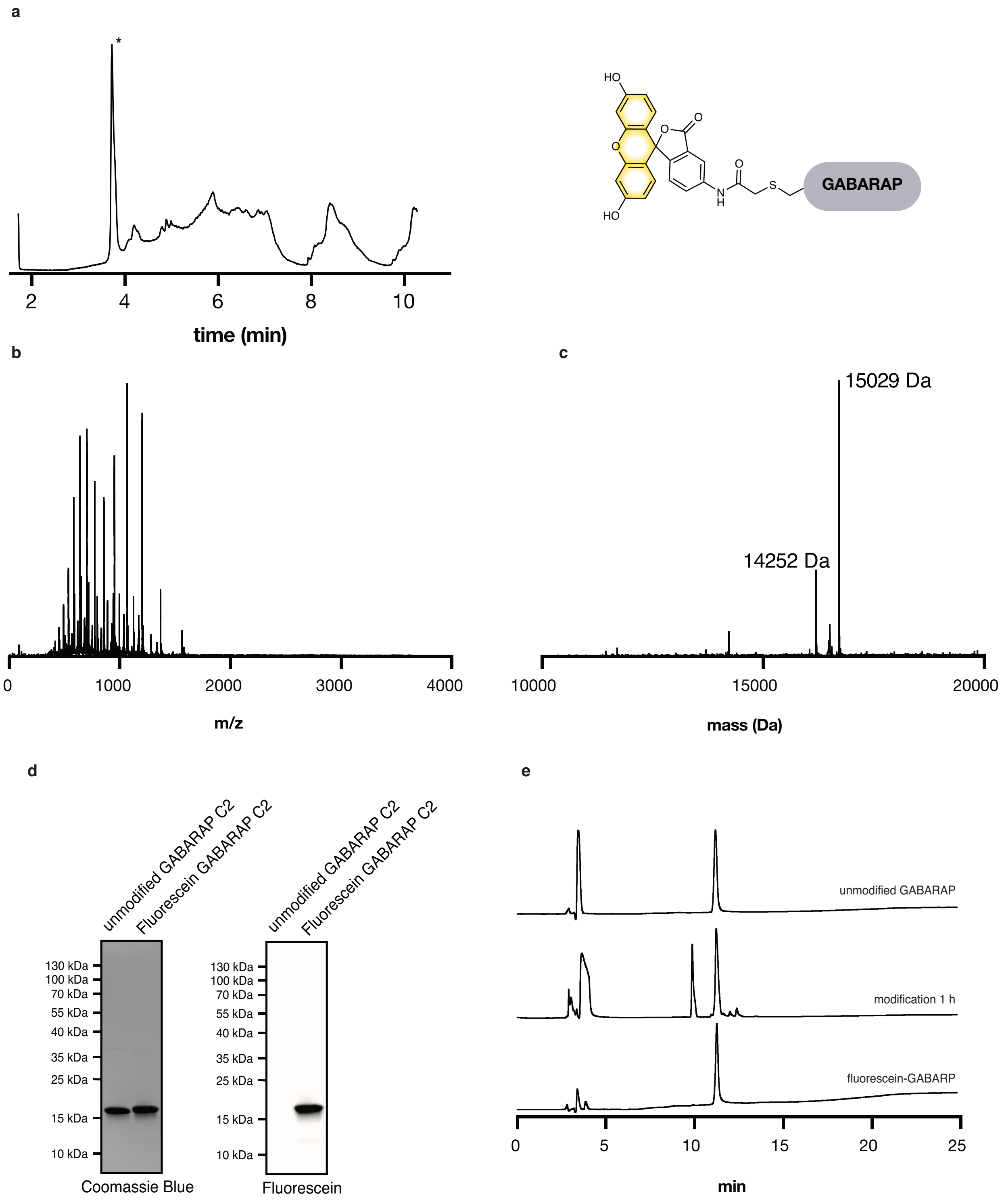

**Supplementary Figure 18** Characterization of fluorescein-GABARAP **a** LC-chromatogram of fluorescein-GABARAP. TIC is shown. Protein peak is marked with a star. **b** Mass-spectrum (ESI) of fluorescein-GABARAP. **c** Deconvoluted mass spectrum of fluorescein-GABARAP. Expected mass 15029 Da. **d** SDS-PAGE gel analysis of GABARAP before and after modification with 5-Iodoacetamide fluorescein. Analysis by Coomassie Blue staining (left gel) and in-gel fluorescence (right gel). **E** RP-HPLC chromatograms showing unmodified GABARAP, crude chromatogram of GABARAP modification with 5-Iodoacetamide fluorescein, and purified fluorescein-GABARAP.

#### 5. Amino acid sequences of recombinantly expressed proteins

| Protein | Amino acid sequence | Comments |
|---|---|---|
| Ubiquitin (ΔGG)-**GyrA-His$_6$** | MQIFVKTLTGKTITLEVEPSDTIENVKAKIQDKEGIPPDQQRLIFAGKQL<br>EDGRTLSDYNIQKESTLHLVLRLR**CITGDALVALPEGESVRIADIVPGAR**<br>**PNSDNAIDLKVLDRHGNPVLADRLFHSGEHPVYTVRTVEGLRVTGTANHP**<br>**LLCLVDVAGVPTLLWKLIDEIKPGDYAVIQRSAFSVDCAGFARGKPEFAP**<br>**TTYTVGVPGLVRFLEAHHRDPDAQAIADELTDGRFYYAKVASVTDAGVQP**<br>**VYSLRVDTADHAFITNGFVSHALEHHHHHH** | *Mxe*-GyrA intein with C-terminal His$_6$ |
| SUMO-GABARAP (ΔG116)-**GyrA-His$_6$** | **MGSDSEVNQEAKPEVKPEVKPETHINLKVSDGSSEIFFKIKKTTPLRRLM**<br>**EAFAKRQGKEMDSLRFLYDGIRIQADQTPEDLDMEDNDIIEAHREQIGG**K<br>FVYKEEHPFEKRRSEGEKIRKKYPDRVPVIVEKAPKARIGDLDKKKYLVP<br>SDLTVGQFYFLIRKRIHLRAEDALFFFVNNVIPPTSATMGQLYQEHHEED<br>FFLYIAYSDESVY**CITGDALVALPEGESVRIADIVPGARPNSDNAIDLKV**<br>**LDRHGNPVLADRLFHSGEHPVYTVRTVEGLRVTGTANHPLLCLVDVAGVP**<br>**TLLWKLIDEIKPGDYAVIQRSAFSVDCAGFARGKPEFAPTTYTVGVPGLV**<br>**RFLEAHHRDPDAQAIADELTDGRFYYAKVASVTDAGVQPVYSLRVDTADH**<br>**AFITNGFVSHALEHHHHHH** | *Mxe*-GyrA intein with C-terminal His$_6$ |
| **His$_6$-SUMO**-GABARAP | **MGSSHHHHHHGSDSEVNQEAKPEVKPEVKPETHINLKVSDGSSEIFFKIK**<br>**KTTPLRRLMEAFAKRQGKEMDSLRFLYDGIRIQADQTPEDLDMEDNDIIE**<br>**AHREQIGG**KFVYKEEHPFEKRRSEGEKIRKKYPDRVPVIVEKAPKARIGD<br>LDKKKYLVPSDLTVGQFYFLIRKRIHLRAEDALFFFVNNVIPPTSATMGQ<br>LYQEHHEEDFFLYIAYSDESVYG | N-terminal His$_6$ with SUMO-tag |
| SUMO-LC3A(ΔG120) **-GyrA-His$_6$** | **MGSDSEVNQEAKPEVKPEVKPETHINLKVSDGSSEIFFKIKKTTPLRRLM**<br>**EAFAKRQGKEMDSLRFLYDGIRIQADQTPEDLDMEDNDIIEAHREQIGG**M<br>PSDRPFKQRRSFADRCKEVQQIRDQHPSKIPVIIERYKGEKQLPVLDKTK<br>FLVPDHVNMSELVKIIRRRLQLNPTQAFFLLVNQHSMVSVSTPIADIYEQ<br>EKDEDGFLYMVYASQETF**CITGDALVALPEGESVRIADIVPGARPNSDNA**<br>**IDLKVLDRHGNPVLADRLFHSGEHPVYTVRTVEGLRVTGTANHPLLCLVD**<br>**VAGVPTLLWKLIDEIKPGDYAVIQRSAFSVDCAGFARGKPEFAPTTYTVG**<br>**VPGLVRFLEAHHRDPDAQAIADELTDGRFYYAKVASVTDAGVQPVYSLRV**<br>**DTADHAFITNGFVSHALEHHHHHH** | *Mxe*-GyrA intein with C-terminal His$_6$ |
| **His$_6$-SUMO**-ATG3 wt | **MGSSHHHHHHGSDSEVNQEAKPEVKPEVKPETHINLKVSDGSSEIFFKIK**<br>**KTTPLRRLMEAFAKRQGKEMDSLRFLYDGIRIQADQTPEDLDMEDNDIIE**<br>**AHREQIGG**QNVINTVKGKALEVAEYLTPVLKESKFKETGVITPEEFVAAG<br>DHLVHHCPTWQWATGEELKVKAYLPTGKQFLVTKNVPCYKRCKQMEYSDE<br>LEAIIEEDDGDGGWVDTYHNTGITGITEAVKEITLENKDNIRLQDCSALC<br>EEEEDEDEGEAADMEEYEESGLLETDEATLDTRKIVEACKAKTDAGGEDA<br>ILQTRTYDLYITYDKYYQTPRLWLFGYDEQRQPLTVEHMYEDISQDHVKK<br>TVTIENHPHLPPPPMCSVHPCRHAEVMKKIIETVAEGGGELGVHMYLLIF<br>LKFVQAVIPTIEYDYTRHFTM | N-terminal His$_6$ with SUMO-tag |
| **His$_6$-SUMO**-ATG3 ΔLIR | **MGSSHHHHHHGSDSEVNQEAKPEVKPEVKPETHINLKVSDGSSEIFFKIK**<br>**KTTPLRRLMEAFAKRQGKEMDSLRFLYDGIRIQADQTPEDLDMEDNDIIE**<br>**AHREQIGG**QNVINTVKGKALEVAEYLTPVLKESKFKETGVITPEEFVAAG<br>DHLVHHCPTWQWATGEELKVKAYLPTGKQFLVTKNVPCYKRCKQMEYSDE<br>LRGHNTGITGITEAVKEITLENKDNIRLQDCSALCEEEEDEDEGEAADME<br>EYEESGLLETDEATLDTRKIVEACKAKTDAGGEDAILQTRTYDLYITYDK<br>YYQTPRLWLFGYDEQRQPLTVEHMYEDISQDHVKKTVTIENHPHLPPPPM<br>CSVHPCRHAEVMKKIIETVAEGGGELGVHMYLLIFLKFVQAVIPTIEYDY<br>TRHFTM | N-terminal His$_6$ with SUMO-tag |
| **His$_6$-SUMO**-ATG3 C1 | **MGSSHHHHHHGSDSEVNQEAKPEVKPEVKPETHINLKVSDGSSEIFFKIK**<br>**KTTPLRRLMEAFAKRQGKEMDSLRFLYDGIRIQADQTPEDLDMEDNDIIE**<br>**AHREQIGG**NVINTVKGKALEVAEYLTPVLKESKFKETGVITPEEFVAAGD<br>HLVHHAPTWQWATGEELKVKAYLPTGKQFLVTKNVPAYKRAKQMEYSDEL<br>EAIIEEDDGDGGWVDTYHNTGITGITEAVKEITLENKDNIRLQDASALAE<br>EEEDEDEGEAADMEEYEESGLLETDEATLDTRKIVEAAKAKTDAGGEDAI<br>LQTRTYDLYITYDKYYQTPRLWLFGYDEQRQPLTVEHMYEDISQDHVKKT<br>VTIENHPHLPPPPMASVHPCRHAEVMKKIIETVAEGGGELGVHMYLLIFL<br>KFVQAVIPTIEYDYTRHFTM | N-terminal His$_6$ with SUMO-tag |

| | | |
|---|---|---|
| **His$_6$-SUMO-** ATG3 C1 ΔLIR | **MGSSHHHHHHGSDSEVNQEAKPEVKPEVKPETHINLKVSDGSSEIFFKIK**<br>**KTTPLRRLMEAFAKRQGKEMDSLRFLYDGIRIQADQTPEDLDMEDNDIIE**<br>**AHREQIGG**NVINTVKGKALEVAEYLTPVLKESKFKETGVITPEEFVAAGD<br>HLVHHAPTWQWATGEELKVKAYLPTGKQFLVTKNVPAYKRAKQMEYSDEL<br>RGHNTGITGITEAVKEITLENKDNIRLQDASALAEEEEDEDEGEAADMEE<br>YEESGLLETDEATLDTRKIVEAAKAKTDAGGEDAILQTRTYDLYITYDKY<br>YQTPRLWLFGYDEQRQPLTVEHMYEDISQDHVKKTVTIENHPHLPPPPMA<br>SVHPCRHAEVMKKIIETVAEGGGELGVHMYLLIFLKFVQAVIPTIEYDYT<br>RHFTM | N-terminal His$_6$ with SUMO-tag |
| **His$_6$-SUMO-** LC3A | **MGSSHHHHHHGSDSEVNQEAKPEVKPEVKPETHINLKVSDGSSEIFFKIK**<br>**KTTPLRRLMEAFAKRQGKEMDSLRFLYDGIRIQADQTPEDLDMEDNDIIE**<br>**AHREQIGG**MPSDRPFKQRRSFADRCKEVQQIRDQHPSKIPVIIERYKGEK<br>QLPVLDKTKFLVPDHVNMSELVKIIRRRLQLNPTQAFFLLVNQHSMVSVS<br>TPIADIYEQEKDEDGFLYMVYASQETFG | N-terminal His$_6$ with SUMO-tag |
| His-TEV-Ube2K (C170S) | MGSSHHHHHHSSGAENLYFQGMANIAVQRIKREFKEVLKSEETSKNQIKV<br>DLVDENFTELRGEIAGPPDTPYEGGRYQLEIKIPETYPFNPPKVRFITKI<br>WHPNISSVTGAICLDILKDQWAAAMTLRTVLLSLQALLAAAEPDDPQDAV<br>VANQYKQNPEMFKQTARLWAHVYAGAPVSSPEYTKKIENLSAMGFDRNAV<br>IVALSSKSWDVETATELLLSN | N-terminal His$_6$ with TEV cleavage site |
| His-thrombin-SENP1 (419-644) | MGSSHHHHHHSSGLVPRGSHMEFPEITEEMEKEIKNVFRNGNQDEVLSEA<br>FRLTITRKDIQTLNHLNWLNDEIINFYMNMLMERSKEKGLPSVHAFNTFF<br>FTKLKTAGYQAVKRWTKKVDVFSVDILLVPIHLGVHWCLAVVDFRKKNIT<br>YYDSMGGINNEACRILLQYLKQESIDKKRKEFDTNGWQLFSKKSQEIPQQ<br>MNGSDAGMFACKYADCITKDRPINFTQQHMPYFRKRMVWEILHRKLL | N-terminal His$_6$ with thrombin cleavage site |

**Supplementary Table S3** Amino acid sequences of recombinantly expressed proteins used in this study. Purification handles cleaved after expression are shown in bold.

#### 6. Primer sequences

| Protein | Mutation | Primer | Sequence |
|---|---|---|---|
| **His$_6$-SUMO-** ATG3 | C264A | Forward primer (5'–3') | AGTTCACCCAGCGAGGCATGCTGAGG |
| | | Reverse primer (5'–3') | GAACACATGGGAGGTGGT |
| **His$_6$-SUMO-** ATG3 | LIR deletion | Forward primer (5'–3') | GGCCACAACACAGGTATTACAGGAA |
| | | Reverse primer (5'–3') | CCGCAATTCATCTGAATATTCCATCTGTTT |
| **His$_6$-SUMO-** ATG3 | Y90A | Forward primer (5'–3') | GCGTCAGATGAATTGGAAGCTATCATTGAAGAAGA |
| | | Reverse primer (5'–3') | TTCCATCTGTTTCGCCCGC |
| **His$_6$-SUMO-** ATG3 | D92A | Forward primer (5'–3') | GCGGAATTGGAAGCTATCATTGAAGAAGATGATGG |
| | | Reverse primer (5'–3') | TGAATATTCCATCTGTTTCGCCCG |
| **His$_6$-SUMO-** ATG3 | E95A | Forward primer (5'–3') | AGATGAATTGGCGGCTATCATTGAAG |
| | | Reverse primer (5'–3') | GAATATTCCATCTGTTTCG |
| **His$_6$-SUMO-** ATG3 | I97A | Forward primer (5'–3') | ATTGGAAGCTGCGATTGAAGAAGATGATG |
| | | Reverse primer (5'–3') | TCATCTGAATATTCCATCTG |
| **His$_6$-SUMO-** ATG3 | I98A | Forward primer (5'–3') | GGAAGCTATCGCGGAAGAAGATGATGG |
| | | Reverse primer (5'–3') | AATTCATCTGAATATTCCATCTG |

| | | | |
|---|---|---|---|
| **$His_6$-SUMO-**ATG3 | E99A | Forward primer (5'–3') | GCGGAAGATGATGGTGATGGCGGATG |
| | | Reverse primer (5'–3') | AATGATAGCTTCCAATTCATCTGAATATTCCATC |
| **$His_6$-SUMO-**ATG3 | D102A | Forward primer (5'–3') | GCGGGTGATGGCGGATGGGTAGA |
| | | Reverse primer (5'–3') | ATCTTCTTCAATGATAGCTTCCAATTCATCTG |
| **$His_6$-SUMO-**ATG3 | D104A | Forward primer (5'–3') | GCGGGCGGATGGGTAGATACATATCACA |
| | | Reverse primer (5'–3') | ACCATCATCTTCTTCAATGATAGCTTCCA |
| **$His_6$-SUMO-**ATG3 | W107A | Forward primer (5'–3') | GCGGTAGATACATATCACAACACAGGTATTACAGGA |
| | | Reverse primer (5'–3') | TCCGCCATCACCATCATCTTCT |
| **$His_6$-SUMO-**ATG3 | V108A | Forward primer (5'–3') | GCGGATACATATCACAACACAGGTATTACAGGAA |
| | | Reverse primer (5'–3') | CCATCCGCCATCACCATCATC |
| **$His_6$-SUMO-**ATG3 | D109A | Forward primer (5'–3') | GCGACATATCACAACACAGGTATTACAGGAATAACG |
| | | Reverse primer (5'–3') | TACCCATCCGCCATCACCATCA |
| **$His_6$-SUMO-**ATG3 | T110A | Forward primer (5'–3') | GCGTATCACAACACAGGTATTACAGGAATAACGGA |
| | | Reverse primer (5'–3') | ATCTACCCATCCGCCATCACC |
| **$His_6$-SUMO-**ATG3 | Y111A | Forward primer (5'–3') | GCGCACAACACAGGTATTACAGGAATAACGG |
| | | Reverse primer (5'–3') | TGTATCTACCCATCCGCCATCAC |

**Supplementary Table S4** Primer sequences used to introduce deletions and point mutations for proteins used in this study.

| experiment | primer ID | primer | sequence |
|---|---|---|---|
| sgRNA production | sgATG3.KD1_seq | Forward primer (5'–3') | TTGGGACTCCCTGGCCCCTGACAGTTTAAGAGC |
| sgRNA production | | Reverse primer (5'–3') | TTAGCTCTTAAAC**T**GTCAGGGGCCAGGGAGTCCCAACAAG |
| sgRNA production | sgATG3.KD2_seq | Forward primer (5'–3') | TTGGTCACGTGAGGCCCCGGTGGGTTTAAGAGC |
| sgRNA production | | Reverse primer (5'–3') | TTAGCTCTTAAACCCACCGGGGCCTCACGTGACCAACAAG |
| sgRNA production | sgATG3.KD3_seq | Forward primer (5'–3') | TTGGCCCGGCTGGCAGCACCCGAGTTTAAGAGC |
| sgRNA production | | Reverse primer (5'–3') | TTAGCTCTTAAACTCGGGTGCTGCCAGCCGGGCCAACAAG |
| sgRNA production | sgATG3.KD4_seq | Forward primer (5'–3') | TTGGGAAAGTGCAGCCGTGTCAGGTTTAAGAGC |
| sgRNA production | | Reverse primer (5'–3') | TTAGCTCTTAAACCTGACACGGCTGCACTTTCCCAACAAG |

**Supplementary Table S5** Primer sequences used to generate sgRNAs for ATG3 knock-down. Protospacer sequences are underlined.

| experiment | primer ID | primer | sequence |
|---|---|---|---|
| sgRNA production | T7FwdVar-sgATG3.KO3 | extension PCR template (5'-3') | GGATCCTAATACGACTCACTATAGGTGAACTGAACACATGGGGTTTTAGAGCTAGAA |
| sgRNA production | T7FwdVar-sgATG3.KO4 | extension PCR template (5'-3') | GGATCCTAATACGACTCACTATAGTGGCTGAGTACCTGACCCGTTTTAGAGCTAGAA |
| sgRNA production | T7RevLong | extension PCR template (5'-3') | AAAAAAGCACCGACTCGGTGCCACTTTTTCAAGTTGATAACGGACTAGCCTTATTTTAACTTGCTATTTCTAGCTCTAAAAC |
| sgRNA production | T7FwdAmp | Forward primer (5'–3') | GGATCCTAATACGACTCACTATAG |
| sgRNA production | T7RevAmp | Reverse primer (5'–3') | AAAAAAGCACCGACTCGG |
| clone genotyping | sgATG3.KO3_seq | Forward primer (5'–3') | CTTTCAGCAACGGCAGCCTT |
| clone genotyping | | Reverse primer (5'–3') | AGCACAAGAAAATTTCTAAGGAGGAC |
| clone genotyping | sgATG3.KO4_seq | Forward primer (5'–3') | CACGGCTGCACTTTCCATCC |
| clone genotyping | | Reverse primer (5'–3') | CTCGAGCCCTACTGCCTTCC |

**Supplementary Table S6** Primer sequences used to generate sgRNAs for ATG3 knockout and for PCR amplification of edited loci. Protospacer sequences are underlined.

#### 7. References

1 Raasi, S. & Pickart, C. M. Ubiquitin chain synthesis. *Methods Mol Biol* **301**, 47-55, (2005).

2 Kabsch, W. Xds. *Acta Crystallogr D Biol Crystallogr* **66**, 125-132, (2010).

3 Huber, J. *et al.* An atypical LIR motif within UBA5 (ubiquitin like modifier activating enzyme 5) interacts with GABARAP proteins and mediates membrane localization of UBA5. *Autophagy* **16**, 256-270, (2020).

4 McCoy, A. J. *et al.* Phaser crystallographic software. *J Appl Crystallogr* **40**, 658-674, (2007).

5 Afonine, P. V. *et al.* Towards automated crystallographic structure refinement with phenix.refine. *Acta Crystallogr D Biol Crystallogr* **68**, 352-367, (2012).

6 Emsley, P. & Cowtan, K. Coot: model-building tools for molecular graphics. *Acta Crystallogr D* **60**, 2126-2132, (2004).

7 Pettersen, E. F. *et al.* UCSF ChimeraX: Structure visualization for researchers, educators, and developers. *Protein Sci* **30**, 70-82, (2021).

8 Larson, M. H. *et al.* CRISPR interference (CRISPRi) for sequence-specific control of gene expression. *Nat Protoc* **8**, 2180-2196, (2013).

9 Lingeman, E., Jeans, C. & Corn, J. E. Production of Purified CasRNPs for Efficacious Genome Editing. *Curr Protoc Mol Biol* **120**, 31 10 31-31 10 19, (2017).

10 Michlits, G. *et al.* Multilayered VBC score predicts sgRNAs that efficiently generate loss-of-function alleles. *Nat Methods* **17**, 708-716, (2020).
